# Cross-stress gene expression atlas of *Marchantia polymorpha* reveals the hierarchy and regulatory principles of abiotic stress responses

**DOI:** 10.1101/2021.11.12.468350

**Authors:** Qiao Wen Tan, Peng Ken Lim, Zhong Chen, Asher Pasha, Nicholas Provart, Marius Arend, Zoran Nikoloski, Marek Mutwil

## Abstract

Abiotic stresses negatively impact ecosystems and the yield of crops, and climate change will increase their frequency and intensity. Despite progress in understanding how plants respond to individual stresses, our knowledge of plant acclimatization to combined stresses–typically occurring in nature is still lacking. Here, we used a plant with minimal regulatory network redundancy, *Marchantia polymorpha*, to study how seven abiotic stresses, alone and in 19 pairwise combinations, affect the phenotype, gene expression, and activity of cellular pathways. We found a high divergence of transcriptomic stress responses between *Arabidopsis* and *Marchantia*, suggesting that the stress-specific gene regulatory networks (GRNs) between bryophytes and angiosperms are not strongly conserved. The reconstructed high-confidence GRNs demonstrated that the response to specific stresses dominates those of others by relying on a large ensemble of transcription factors. We also showed that a regression model could accurately predict the gene expression under combined stresses, indicating that *Marchantia* performs arithmetic addition to respond to multiple stresses. Finally, we provide two online resources (https://conekt.plant.tools and http://bar.utoronto.ca/efp_marchantia/cgi-bin/efpWeb.cgi) to facilitate the study of gene expression in Marchantia exposed to abiotic stresses.

## Introduction

The colonization of land by plants, which occurred around 470 Ma, was essential to establish habitable environments on land for all kingdoms of life ^1^. Bryophytes, which include mosses, liverworts, and hornworts, represent the earliest diverging group of non-vascular land plants ^2–4^. Morphology of the earliest land plant fossils, consisting primarily of tissue fragments and spores from the Middle Ordovician around 470 Ma, showed that early land plants were liverwort-like ^5, 6^. As a liverwort, *Marchantia polymorpha* is a valuable model to better understand the emergence and evolution of land plants, as it allows us to compare the biology of aquatic algae and non-vascular plants to vascular, seed, and flowering plants. Studying *Marchantia* can help us better understand the successful terrestrialization event, as *Marchantia* contains traits essential for this task along with increased complexity (e.g., hormones auxin, jasmonate, salicylic acid, protection mechanisms against desiccation, photooxidative damage)^7^ which is still considerably lower than that of vascular plants ^8^.

Besides its interesting evolutionary position among land plants, Marchantia is a valuable model for studying basic plant biology. Mainly due to the lack of whole-genome duplications in the liverwort lineage, *Marchantia* shows a simpler, low-redundancy regulatory genome ^8^, which together with the ease of growth and genetic manipulation ^9^, makes Marchantia an excellent model to study general plant biology. The *Marchantia* genome contains necessary components for most land-plant signaling pathways with low redundancy, making it easier to dissect the pathways ^8^. For example, the auxin signaling network in *Marchantia* is simple yet functional, with all relevant genes existing as single orthologs ^10^. Similarly, cellulose biosynthesis in Marchantia uses the same but simplified machinery; while *Arabidopsis thaliana* contains ten cellulose synthases in multimeric complexes, Marchantia has only two ^11^. Furthermore, since the dominant generation of *Marchantia* is the haploid gametophyte, heterozygosity can be circumvented to directly study mutant and transgenic phenotypes. These, and other features, make *Marchantia* an attractive model for dissecting the function of genes and biological pathways.

Extreme environmental conditions can cause a multitude of biotic and abiotic stresses that can devastate crops and induce the collapse of entire ecosystems ^12, 13^. Plants have evolved sophisticated mechanisms to perceive and respond to the different stresses, which induces an acclimation process that allows the plant to survive the stress ^14^, but often at the cost of reduced growth ^12, 13^. Many studies have analyzed the effect of stress on plant growth by identifying differentially expressed genes between stress-treated and untreated plants (e.g. ^15–17)^, or by identifying single nucleotide polymorphisms associated with stress resistance in *Arabidopsis* and maize ^18–20^. The studies performed on model plants such as *Arabidopsis* can shed light on fundamental processes of stress acclimation, but it is currently unclear whether the acclimation mechanisms are conserved and transferable to crops.

In addition, plants are often exposed to a combination of stresses in the natural environment which may require opposing strategies to mitigate adverse effects. For example, drought causes plants to close their stomata to minimize water loss ^21, 22^, while heat causes the stomata to open to cool down their leaves via transpiration ^17, 23^. Stress signaling is mediated by a diverse ensemble of stress-specific sensors/receptors, networks of protein kinases/phosphatases, calcium channels/pumps, and transcription factors that can be localized to different organelles ^24, 25^. Stress signaling is further communicated and modulated by hormones, signaling molecules (e.g., reactive oxygen species), and other protein modifications (S-nitrosylation, ubiquitination, myristoylation) ^26, 27^. While this complexity renders it challenging to elucidate the molecular basis of stress responses to single or combined stresses, the key features of the model Marchantia provide the means to deepen our understanding of stress acclimation.

In this study, we set to take advantage of *Marchantia*’s less complex regulatory architecture to understand better how plants respond to environmental cues such as stress to modulate the expression of genes and biological pathways. To this end, we constructed an abiotic stress gene expression atlas of *Marchantia* comprising seven abiotic stresses, i.e.darkness, high light, cold, heat, nitrogen deficiency, salt, and mannitol, and their 19 pairwise combinations. For each stress, we identified robustly-responding transcription factors that are likely important for Marchantia’s survival to the stress. Comparing these transcription factors to gene expression responses and biological function of *Arabidopsis thaliana* orthologs revealed poor agreement between the two plants, suggesting a large divergence of stress-related gene regulatory networks. Interestingly, the analysis of differentially expressed genes and biological pathways in the combined stresses revealed that certain stresses (e.g., darkness and heat) induce large transcriptomic responses that dominate other stresses (e.g., salt and mannitol). The dominant stresses express a larger ensemble of transcription factors that change the expression of more genes and pathways than the non-dominant stresses. Importantly, we construct a linear regression model that can explain the gene expression changes of combined stresses, showing that Marchantia performs an arithmetic addition to integrate environmental cues. Finally, to provide bioinformatical resources, we provide i) an eFP browser for Marchantia (http://bar.utoronto.ca/efp_marchantia/cgi-bin/efpWeb.cgi) that allows the visualization of gene expression in organs and stress conditions and ii) an updated CoNekT platform (https://conekt.plant.tools)28, allowing sophisticated comparative gene expression and co-expression analyses.

## Material and Methods

### Maintenance of *Marchantia polymorpha*

Male *Marchantia polymorpha*, accession Takaragaike-1 was propagated on half-strength Gamborg B-5 Basal agar (1% sucrose, pH 5.5, 1.4% agar)(Gamborg et al., 1968) in deep well plates (SPL Biosciences, SPL 310101) at 24°C under continuous LED light at 60 μEm^-2^s^-1^.

### Experimental setup for stress experiment

To determine the ideal stress condition for cross-stress experiments, a series of severity levels was used for heat, cold, salt, osmotic, light, dark, and nitrogen deficiency stresses. For each stress level, three agar plates containing nine gemmae each were plated, where two plates were used as material for RNA sequencing, and one plate was kept for observation until 21 days after plating (DAP).

For salt, osmotic, and nitrogen deficiency stress, the gemmae were plated on half-strength Gamborg B-5 Basal agar (pH 5.5, 1.4% agar, 12.4 mM KNO_3_) supplemented with 20 - 200 mM NaCl (20 mM steps), 50-400 mM mannitol (50 mM steps) and KNO_3_ concentration ranging from 90 to 0% (22.3 - 0 mM, 2.5 mM steps), respectively. The potassium concentration in nitrogen deficiency agar was maintained using equimolar concentrations of KCl. Gemmae subjected to heat, cold, light, and dark stress were plated on normal half-strength Gamborg B5 agar (pH 5.5, 1.4% agar, 12.4 mM KNO_3_, 0.5 mM (NH_4_)_2_SO_4_).

Plates were grown at 24°C under continuous LED light at 60 μEm^-2^s^-1^ from days 0 to 13. For dark stress, plates were moved to the plant growth chamber (HiPoint M- 313) on days 8, 9, 10, 11, 12, 13, and 14 for growth in darkness at 24°C for 7, 6, 5, 4, 3, 2 and 1 day(s), respectively. On day 14, all plates were transferred to the plant growth chamber (HiPoint M-313) for 24 hours. Control and plates for salt, osmotic, and nitrogen deficiency stresses were maintained at 24°C under continuous LED light at 60 μEm^-2^s^-1^ in the plant growth chamber. The following modifications used normal growth conditions for heat, cold, dark, and light stresses. For heat and cold stress, 27°C to 36°C (3°C steps) and 3°C to 12°C (3°C steps) were used, respectively. For light stress, plants were subjected to 20% to 100% chamber capacity of light output (115 to 535 μEm^-2^s^-1^) at steps of 20%, approximating 100 μEm^-2^s^-1^ per step.

On day 15, whole plants were harvested by pooling three plants into 2 mL Eppendorf tubes, flash-frozen in liquid nitrogen, and stored at -80°C. This was done in triplicates, resulting in 3 (replicates) * 3 (pooled plants) used for each RNA-seq sample. Images of the front and back of the observation plates were taken, and the plates were returned to normal growth conditions. Pictures of the front and back of the observation plates were taken on day 21 without cover.

Cross stress experiments were carried out similarly with the following parameters for the various stress combinations - heat (33°C), cold (3°C), salt (40 mM NaCl), osmotic (100 mM mannitol), light (435 μMm^-2^s^-1^), darkness (3 days) and nitrogen deficiency (0 mM KNO_3_).

### Size measurement of *Marchantia polymorpha*

To measure the size of the plants grown under the different stresses, the images of the observation plates were scaled and measured in Adobe Illustrator. The length and breadth of the thallus were taken in relation to the central axis (Figure 1B shows the plants with the central axis in a horizontal position). The thallus’s approximate area was derived from the product of the length and breadth of the thallus. Abnormally small plants (outliers) were excluded from further analysis.

**Figure 1.**
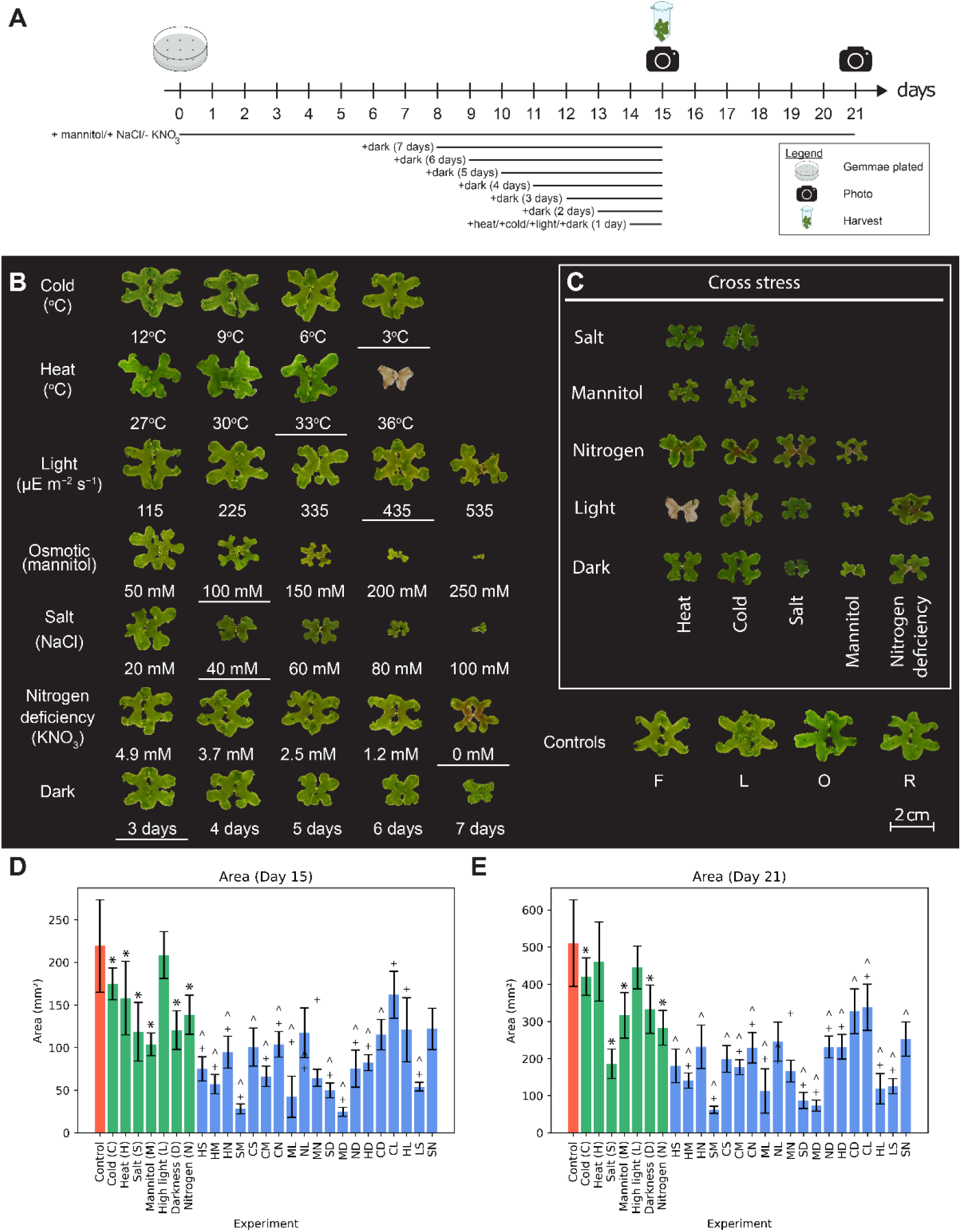
Influence of different abiotic stresses on the growth of *Marchantia polymorpha*. A) Overview of the experimental setup for stress experiments. Black lines below the timeline indicate the duration where plants were exposed to stress. Plants were sampled on day 15. Observation plates were returned to normal growth conditions on day 15, and photographs of the plates were taken on days 15 and 21. B) Phenotype of plants on day 21 for heat, cold, osmotic, salt, light, dark, and nitrogen deficiency stresses. Conditions with underline represent the stress intensities used for combined stress. Controls F and L are representatives from the batches at the start and end of independent stresses in panel B. C) Phenotypes of plants on day 21 for combined stresses. Controls O and R are representatives from the start and end of combined stresses in panel C. D and E) Thallus size of gemmalings on Day 15 and 21 of growth (error bars are represented by standard deviation and Student’s two-tailed t-test, p-value < 0.05. Asterisk (*) represents a significant difference to control, caret (^) represents a significant difference to the first single stress control, and plus sign (+) indicates a significant difference to the second single stress control.

### RNA extraction and sequencing

Using a mortar and pestle, plants from each Eppendorf tube were ground into fine powder in liquid nitrogen. Total RNA was extracted using the Spectrum^TM^ Plant Total RNA Kit (Sigma, STRN-250) using Protocol A (750 μL Binding Solution) according to manufacturer’s instruction with on-column DNase digestion using 60 μL of DNase mixture (15 μL RQ1 RNase-Free DNase (Promega, M6101), 6 μL RQ1 DNase 10X Reaction Buffer and 19 μL nuclease-free water) per column.

Preliminary quality control of the extracted RNA (triplicates for each condition) was done using Nanodrop before further quality control checks by Novogene (Singapore) for sample quantitation, integrity, and purity using Nanodrop agarose gel electrophoresis and Agilent 2100 Bioanalyzer. Library construction from total RNA, including eukaryotic mRNA enrichment by oligo(dT) beads, library size selection, and PCR enrichment, was performed by Novogene using NEBNext® Ultra™ II Directional RNA Library Prep Kit for Illumina®. The libraries were then sequenced with Illumina Novaseq-6000, paired-end sequencing at 150 base pairs, and sequencing depths of ∼20 million reads per sample.

### Expression quantification

RNA sequencing data were mapped against the *Marchantia polymorpha* CDS (v5.1 revision 1, MarpolBase), quantified, and TPM-normalised (transcript per million) using Kallisto v 0.46.1^29^.

### Identification of differentially expressed genes

Non-normalized counts from Kallisto were used to analyze differentially expressed genes (DEGs) using R package DESeq2 ^30^, where various stress conditions were compared against their respective controls.

For our *M. polymorpha* dataset, only genes that were found to be differentially expressed against controls from two different batches were considered for further analysis. For downstream analysis, only genes with a Benjamini-Hochberg^31^ adjusted p-value < 0.05 and a −1> log_2_-fold >1 were considered as differentially expressed.

### Identification of transcription factors and differentially expressed pathways

The biological function and pathway membership of genes of *Marchantia polymorpha* were annotated using Mercator 4 v2.0 ^32^. The annotation of *M. polymorpha* transcription factors was retrieved from PlantTFDB v5.0 ^33^.

Significantly differentially expressed pathways were determined through a permutation analysis, where the observed number of DEGs in a pathway was compared to the permuted number of DEGs. The p-values were adjusted for multiple testing using Benjamini Hochberg correction (p-value < 0.05) ^31^.

### Construction of stress-specific gene regulatory networks

To reconstruct the gene regulatory network (GRN), we selected DEGs expressed in more than five experiments to ensure sufficient variability in our dataset needed for statistical modeling (Figure S8A). Apart from using all the experiments, we also reconstructed stress-specific networks by employing a subset of experiments that included the respective stresses (Figure S8B). For example, the heat-specific network is based only on data from experiments where heat stress was involved (i.e., Heat, Heat-Mannitol, Heat-Salt, Heat-Dark, and Heat-Nitrogen deficiency).

The GRN was reconstructed using linear regression with elastic net regularization ^34^, which provided a good compromise between model sparsity (i.e., feature selection corresponding to the inclusion of transcription factors in the model) and model explanatory power. Three- and five-fold cross-validation was used for the stress-specific data and all data, respectively, to determine the optimal λ and α for each model. We filtered for good quality models with R^2^ > 0.8 (FigS9A).

### Evaluating the accuracy of the GRNs

We evaluated our GRN network against the known curated gene regulatory networks in *Arabidopsis* retrieved from AGRIS ^35^. Orthogroups of *Arabidopsis* and *Marchantia* genes were identified using Orthofinder v2.3.1 ^36^ and used as the basis for comparing the *Arabidopsis* and *Marchantia* GRNs. We used the Jaccard index to quantify the similarity between the GRNs, where TF-target edges were converted into orthogroup-orthogroup tuples that were used in the set comparisons. The union of the stress-specific networks produced the highest Jaccard index score between our networks and the AGRIS network (Fig S9B). To test the significance of the similarity between our GRN and the AGRIS network, we calculated empirical p-values by shuffling the TF and gene pair in the AGRIS network 1000 times and calculated the resulting Jaccard index for each shuffling. The empirical p-value was calculated by comparing the 1000 shuffled Jaccard index values to the observed Jaccard index value.

### Construction of the unified stress GRN

For each gene, we identified the transcription factor with the highest absolute coefficient (Fig. 3A) in the merged network. In the merged network, the coefficients in different stress-specific networks are used to determine whether the transcription factor is an activator (all coefficients are positive), repressor (all coefficients are negative) or if the regulation is ambiguous (mixture of positive and negative coefficients). For example, if TF X regulates gene Y in 4 different stress-specific networks with a positive coefficient, the TF is considered an activator.

**Figure 3.**
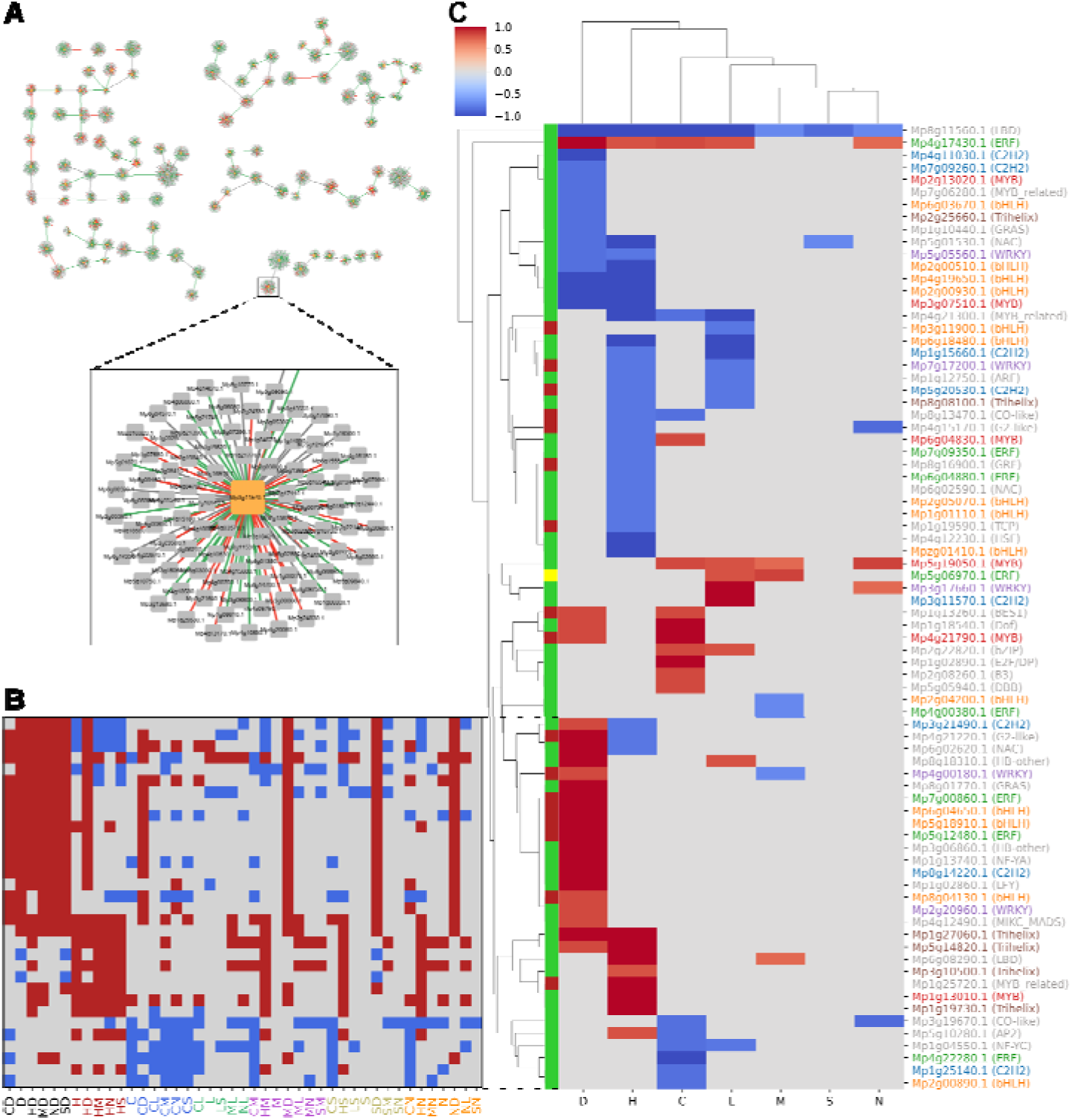
Identification of robustly-responding transcriptional activators and repressors. A) Gene regulatory network constructed from the union of the seven stress-specific networks. For each of the 6257 differentially expressed genes, we kept models with R^2^ > 0.8 and selected one transcription factor with the highest absolut coefficient. Orange and gray nodes represent transcription factors and genes, respectively, while green and re edges represent positive and negative coefficients, respectively. B) Differential expression patterns of transcription factors across stress groups. Red, blue, and gray indicate significantly up-, down-regulated, and unchange expression, respectively. C) Identification of 75 robustly-responding transcription factors across the stresses. For clarity, cells with specificity scores < 0.7 are masked. Red and blue cells indicate the degree of up- and down-regulation, respectively.

### Revealing transcription factors regulating biological pathways

To understand how specifically regulated transcription factors might be affecting certain biological processes, we took transcription factors and second-level Mapman bins that were specifically regulated in a stress group. A stress group is defined as a group of experiments sharing common stress, for example, the heat stress group contains the experiments Heat, Heat-Mannitol, Heat-Salt, Heat-Dark, and Heat-Nitrogen deficiency.

Robustly-responding transcription factors were identified based on the ratio of occurrences where it is differentially regulated in a stress group against the number of experiments in the stress group. A transcription factor is considered to be specifically expressed in the stress group if the ratio > 0.7. Stress group-specific MapMan bins were defined in the same manner.

Robustly-responding MapMan bins (Figure S13) were considered to be regulated by robustly responding TFs if at least 5% of the genes (Figure S14) in the MapMan bin were found to be associated with the TF (Figure 4B and S15).

**Figure 4.**
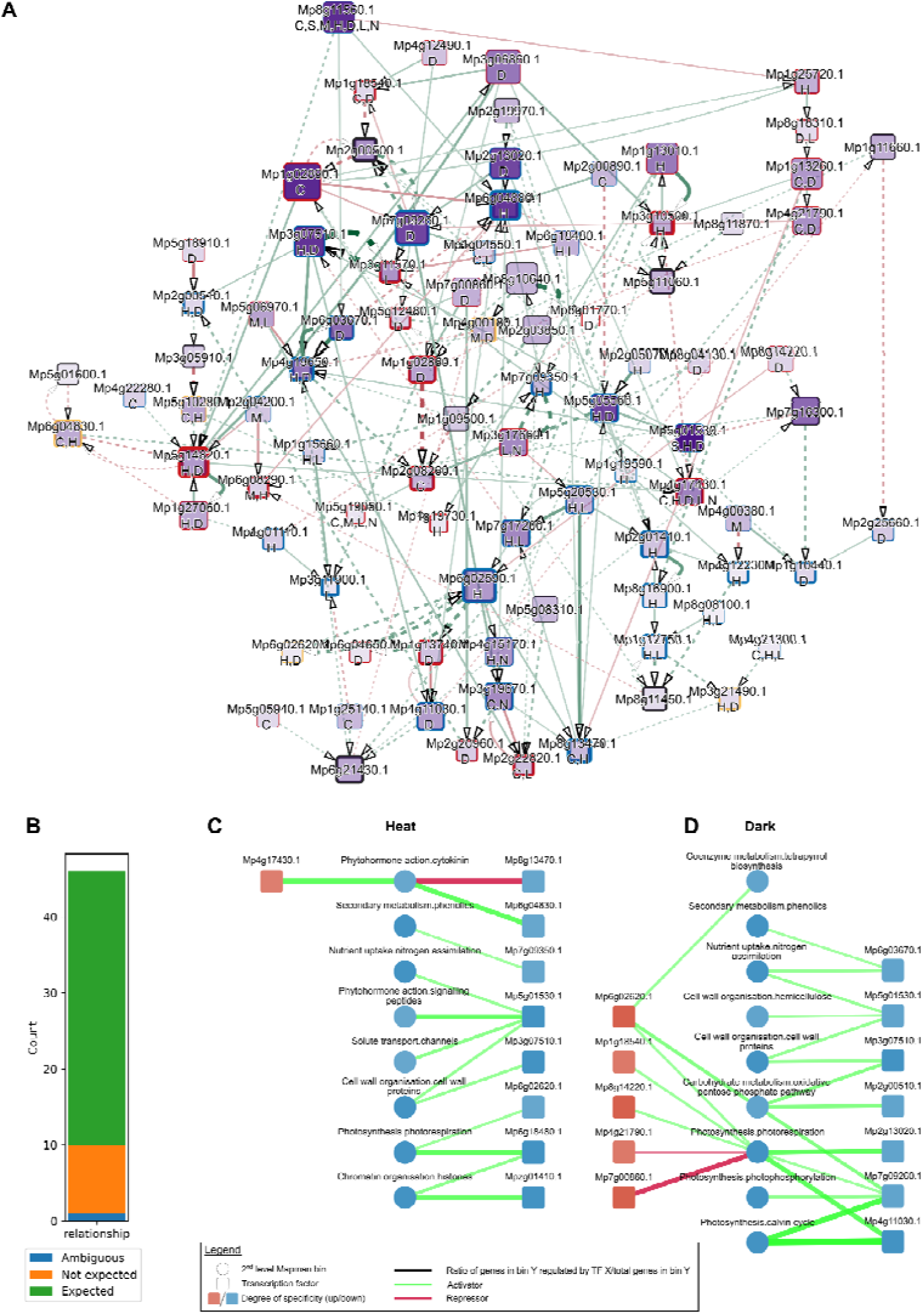
Analysis of stress-specific GRN. A) GRN comprising of TFs differentially regulated in at least five experiments. Labels below the gene name indicate robustly-responding TFs in (C)old, (D)arkness, (H)eat, (L)ight, (M)annitol, (N)itrogen deficiency, and (S)alt. Darker node colors indicate a higher number of genes a TF is regulating, while the node sizes indicate the number of transcription factors the node is regulating. Node border width indicates the number of incoming regulatory signals of the transcription factor. Red, blue, and yellow node border colors indicate TFs that are robustly up-regulated, down-regulated, or both. Thicker edges indicate higher absolute coefficients, where green and red edges represent positive and negative coefficients, respectively. Solid and dashed edges represent expected and unexpected regulations, respectively. D) The number of expected, unexpected, and ambiguous gene regulatory relationships between TFs and MapMan bins. Only edges between TFs controlling ≥5% of genes in a MapMan bin are used. C) Identification of TFs that regulate biological processes during heat stress. Red and blue nodes indicate robustly up- and down-regulated second-level MapMan bins, respectively. Edge thickness represents the percentage of genes controlled by a TF, in a given MapMan bin, with thicker edges indicating a higher percentage. Green and red edges indicate that a TF is up- or down-regulating a given process. D) Darkness-specific TF-ManMan bin network.

### Inference of TF-TF GRN

We chose the highest absolute co-efficient for each TF-TF pair to indicate the putative regulatory relationships between TFs. Next, we defined the transcription factors to be up-regulated, downregulated, or ambiguous (up- and down-regulated in more than 1 stress group) based on their specific expression across stress groups. We then defined the edges as expected if: 1) TF_A_ (up-/down-regulated, activator) regulates TF_B_ (up-/down-regulated), 2) TF_A_ (down-/up-regulated, repressor) regulates TF_B_ (up-/down-regulated). All other edges were defined as unexpected, e.g., TF_A_ (up-regulated, activator) regulates TF_B_ (downregulated). Finally, we applied an absolute coefficient cut-off that produced the highest ratio of expected / total edges (Figure S11), arriving at a cut-off value of 0.22.

### Functional analysis of *Arabidopsis* TF orthologs

The biological function of *Arabidopsis* TF orthologs was inferred from gene ontology terms with experimental evidence and literature searches. The expression responses were inferred through observation of gene expression changes on the *Arabidopsis* eFP browser using the "Abiotic stress" dataset from ^37^.

## Data availability

The RNA-seq data capturing the expression of controls, single and double stresses are available from https://www.ebi.ac.uk/ena as E-MTAB-11141.

## Script availability

Python and bash scripts used to generate the figures in the paper are available from: https://github.com/tqiaowen/marchantia-stress

## Results

### Response of *Marchantia polymorpha* to combined abiotic stresses

To capture gene expression changes caused by a single or combination of stresses, we first established the type and magnitude of stresses to use. We defined two types of stresses: i) the environmental stresses comprised heat, cold, excess light, and prolonged darkness, while ii) media stresses comprised nitrogen deficiency, excess salt (representing ionic and osmotic stress), and excess mannitol (representing osmotic/drought stress). Next, the magnitude of the stresses was modulated to identify near-lethal stress conditions, growth decrease by ∼50% (inferred from the approximate thallus area), or stresses displaying signs of necrosis. To this end, gemmae grown on sealed, sterile agar plates under constant light were subjected to varying degrees of stress, and their phenotypes were observed on days 15 and 21. For media stresses, the gemmae were subjected to the stress from day 0, while for the environmental stresses, the stress was applied on day 14 for 24 hours (Figure 1A). Prolonged darkness was an exception to this design, as plates were subjected to darkness on days 8, 9, 10, 11, 12, 13, and 14 to expose the plants to 7, 6, 5, 4, 3, 2, and 1 day(s) of darkness at day 15, respectively (Figure 1A).

Single stresses showed varying degrees of effect on plant growth on day 15 (day of harvest, Figure S1 shows growth measurements, Supplemental Data 1 shows agar plates) and day 21 (6 days post-stress for environmental stresses, Figure 1B). One day of cold stress did not affect the growth at the temperature range tested (3-12°C), and we selected 3°C for further analysis. The heat stress experiment showed that the plants abruptly died when the heat treatment temperature was increased from 33°C (no phenotype) to 36°C (death), and we selected 33°C for further study. For light stress, we selected 435 µEm^-2^s^-1^ as we observed necrosis at the next higher light intensity (535 µEm^-2^s^-1^, Supplemental Data 1). For osmotic (100 mM selected) and salt stress (40mM selected), we observed an expected negative growth gradient when the concentration of the two compounds was increased. For nitrogen deficiency, at 0 mM KNO_3,_ we observed a decrease in growth and an accumulation of a red pigment, which likely represents auronidin, a flavonoid shown to accumulate during phosphate deficiency ^38^. Finally, for darkness, we observed that the growth of plants decreased proportionately with the duration of days without light, and we selected plants grown in three days of darkness, as they showed a growth decrease of 50% (Figure S1). On day 21 (i.e., six days of normal growth condition), all dark-grown plants showed increased size, indicating that the seven days of darkness are not lethal.

Next, we determined how *Marchantia* responds to a combination of two stresses. To this end, we tested all 19 feasible pairs of stresses (cold+heat and dark+light combinations are mutually exclusive and excluded) using the same experimental outline as for single stresses (Figure 1A). We did not observe any unexpected phenotypes when combining the stresses (Figure 1D), as typically, a combination of stresses resulted in an expected combination of phenotypes (e.g., nitrogen: small, pigmented, mannitol: small, nitrogen+mannitol: even smaller, and pigmented) except for salt-nitrogen (SN), which is significantly larger than its salt counterpart but not different from its nitrogen counterpart (Figure 1E). While plants subjected to a combination of sub-lethal heat (33°C) and high-light (435 µEm^-2^s^-1^) died (Figure 1D), this was most likely due to a greenhouse effect caused by high irradiation and sealed plates, as the temperature of agar climbed to 38°C (i.e., lethal temperature, Figure 1C). Interestingly, we observed yellowing of the thalli for high light and cold, indicating that lowering the temperature makes the plants more sensitive to high light (Figure 1C).

The resulting panel of 7 single stresses, 18 combinations of two stresses, and 2 untreated controls were sent for RNA sequencing (Table S1 contains sample labels, and Tables S2 and S3 contain Transcript Per Million (TPM) values and raw counts, respectively). Overall, we observed a good agreement between the sample replicates, as samples showed expected clustering (Figure S2), and the correlation between expression profiles of replicates was >0.97 (Table S4).

### Combinatorial differential gene expression analysis reveals a hierarchy of stress responses

Plants perceive and respond simultaneously to multiple stresses when growing in nature. To better understand how *Marchantia* responds to a single of combination of two stresses, we first identified differentially expressed genes in the single stresses and the 18 combinations of two stresses. For more robust inferences, we used two controls taken at the beginning and the middle of the experiment and set the requirement for a gene to show conserved differential expression in both controls to be deemed a DEG (Table S5 and S6). Overall, we observed a good agreement between the two controls, as both volcano plots (Figure S3) and set comparisons (Figure S4) indicated a similar set of DEGs. We compared the number of up- and down-regulated genes with the single stresses and observed that the combination of stresses typically contains a similar or higher number of DEGs when compared to single stresses (Figure 2A).

**Figure 2.**
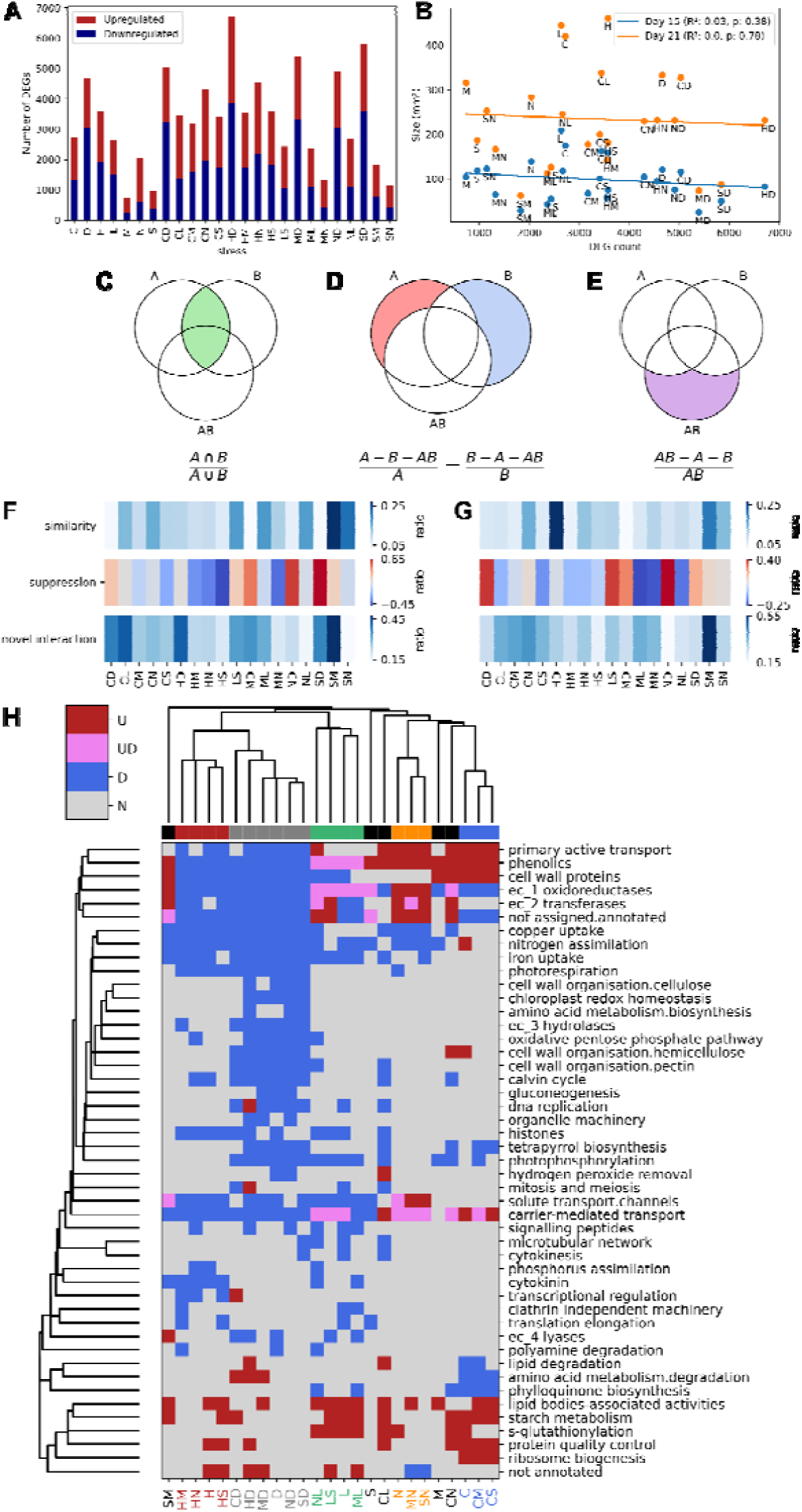
Analysis of differentially expressed genes and biological pathways. A) The number of significantly (adjusted p-value < 0.05) up-regulated (red) and downregulated (blue) differentially expressed genes. The stresses are (C)old, (D)arkness, (H)eat, (L)ight, (M)annitol, (N)itrogen deficiency, and (S)alt. B) The number of DEGs (x-axis) versus the size (y-axis) of Marchantia plants on day 15 (blue) and day 21 (orange). The R^2^ and p-values are shown i the legend. Illustration and equation of metrics used to measure A) similarity between two stresses, B) suppression of one stress when two stresses are combined, and C) novel genes that are differentially regulated when stresses ar combined. Interaction between independent and cross stresses for D) up-regulated genes and E) downregulated genes. In each stress combination, the first (’A’) and second (’B’) stress is the first and second letter of the combine (’AB’) stress, respectively. For the suppression measure, a darker shade of red and blue indicates that more genes from the first (’A’) and second (’B’) stress are not represented in the combined stress, respectively. H) Biological processes that are significantly differentially expressed (adjusted p-value < 0.05). For brevity, we only show Mapman bins that are differentially expressed in at least three stress perturbations. The groups of stresses are color-coded. Abbreviations used to describe the categories of regulation are upregulation (’U’, red), up and downregulation (’UD’, purple), downregulation (’D’, blue), and no change (’N’, gray).

Next, to investigate whether a severe growth phenotype results in a large number of differentially expressed genes, we plotted the plant size (x-axis) and the number of differentially expressed genes (y-axis, Figure 2B). We observed no significant correlation between these two variables for plants on days 15 and 21 (p-value > 0.05, Figure 2B). We concluded that there is no correlation between the severity of growth phenotype and the transcriptomic response to stress. For instance, the smallest plants (salt+mannitol, SM) also had the fewest number of differentially expressed genes.

The different single and combined stresses are likely to elicit similar and unique gene expression responses, resulting in the stresses having similar sets of DEGs. We used the UpSet plot to elucidate these similarities, which shows the intersection of multiple sets for up-regulated (Figure S5A) and down-regulated (Figure S5B) genes. Interestingly, the largest set of up- and down-regulated genes was unique to heat+darkness combined stress (HD), suggesting that HD elicits the most unique and dramatic transcriptional response among the tested stresses. Other unique stress responses comprised upregulated cold+darkness (CD), cold+high light (CL), cold+nitrogen deficiency (CN) and heat+nitrogen deficiency (HN, Figure 4C); and downregulated cold+darkness (CD), cold+nitrogen deficiency (CN), cold (C) and cold+high light (CL). Darkness alone (D) and in combination with other stresses (e.g., ND, CD, MD, SD, HD) also contained a high number of up-regulated (connected dots in columns 4, 5, Figure S5A) and down-regulated (columns 2, 4, Figure S5B) genes, suggesting a conserved, core darkness response. Similarly, we observed core responses to heat (e.g., 6th column, Figure S3A) and cold (10th column, Figure S5A). Interestingly, we also observed a high number of DEGs across heat and darkness experiments (Figure S5A & B), suggesting that these two stresses elicit a similar response to a degree.

To better understand how *Marchantia* responds to a combination of two stresses, we compared the combined response (AB) to the response to individual stresses (A and B), with three different metrics measuring the shared response, the dominance of stress, and novel responses induced by combined stress. We first produced Venn diagrams for up-regulated (Figure S6) and downregulated (Figure S7) gene sets. The first metric measures the similarity between A and B (Figure 2C, green area, Jaccard Index) and ranges from 0 (A, B do not have any DEGs in common) to 1 (A, B elicit identical DEGs). The second metric measures whether one stress suppresses the other (Figure 2D, the difference between red and blue area) and ranges from -1 (AB is similar to A but not to B, i.e., A suppresses B) to 1 (AB is similar to B but not A, i.e., B suppresses A). The third metric measures whether a combination of two stresses elicits a unique response when compared to those of the two individual stresses (Figure 2E, purple area specific to AB) and ranges from 0 (all DEGs are found in A and B, i.e., no novel response) to 1 (all DEGs in AB are unique, i.e., the combination of AB elicited a unique response).

Stresses showing the highest similarity in terms of DEG responses comprise salt and mannitol for up- and down-regulated genes (Figure 2F-G, dark area for SM), salt and nitrogen deficiency for up-regulated genes (Figure 2F), and heat and darkness for downregulated genes (dark area for HD, Figure 2F). The suppression analysis showed that darkness is a strong suppressor for many stresses (red boxes in CD, MD, ND, SD), except HD, indicating that heat stress and darkness are comparably dominant (Figure 2F-G). To support this observation, heat stress could suppress other stresses (blue boxes in HS, HN, HM, Figure 2F-G). Finally, the uniqueness analysis revealed that the salt+mannitol combination elicited DEGs that were not found in the individual stresses (dark blue boxes for SM, Figure 2F-G), suggesting that the two stresses can activate altogether different responses when combined.

To better understand the hierarchies of the stresses and how these stresses affect the activity of biological processes, we identified which processes contain significantly more up-regulated or downregulated DEGs than expected by chance for each stress combination (Figure 2H). The clustering of the rows (biological processes) and columns (stresses) revealed that darkness-containing stresses form a clear group of similar responses (six stress combinations comprising D, HD, CD, MD, SD, and ND), confirming the previous observation of the darkness suppressing other stresses. Interestingly, the darkness caused a strong decrease in gene expression of numerous pathways (Figure 2H, ∼58% blue squares). The second largest group contained nearly all heat stress combinations (four stresses: H, HD, HS, HN) and a similar but less dramatic downregulation of transcripts in many biological processes. Interestingly, despite the dramatic downregulation of biological processes in most dark and heat stresses, a subset of stresses (H, HS, HD, MD) was significantly up-regulated for uncharacterized genes (bin not annotated), suggesting that these responses employ poorly understood genes. The other groups comprised high-light (four stresses L, NL, LS, ML), nitrogen deficiency (three stresses N, MN, and SN), and cold (three stresses C, CM, and CS). In contrast, salt- and mannitol-containing stresses did not form any groups, suggesting that despite dramatic phenotypic changes, these stresses are suppressed by other stresses (Figure 2H, M, and S are not grouped with other stresses). Interestingly, salt+mannitol, cold+high light, and cold+low nitrogen were also not grouped, indicating that these combinations result in novel transcriptomic responses.

Based on these findings, we can rank the strength of dominance of abiotic stresses starting from darkness (six stress combinations), heat and light (four each), nitrogen deficiency and cold (three each), and finally, salt and mannitol (none).

### Identification of high-confidence transcription factors involved in stress response

Our results indicate that certain stresses (e.g., heat and darkness) result in a high number of DEGs (Figure 2). These DEGs are likely a result of the action of a gene regulatory network (GRN) comprising transcription factors (TFs) that are themselves differentially expressed.

To infer the stress-responsive GRN (Figure S8B), we used ElasticNet, for each of the 6257 differentially expressed genes and all 95 differentially expressed TFs that were responsive in more than five experiments employed as response and predictor variables (Figure S8A). In addition to constructing the GRNs from the whole dataset (i.e., all 81 RNA-seq experiments), we also constructed stress-specific GRNs by using the data from those experiments that included the respective stress. For instance, the darkness- specific GRN was inferred from D, DH, DS, DM, and DN expression data. Altogether, we constructed 50,056 ElasticNet models (i.e., 6275 DEGs for eight stress datasets) containing up to 95 differentially expressed transcription factors as predictors. Interestingly, the performance of the GRNs for the individual stresses was higher than for the GRN based on the whole dataset (performance measured by R^2^, Figure S9A), suggesting that there is significant variability between the datasets that can be used for linear modeling at the stress group level but not when all data sets are jointly examined. As a result, the GRNs based on the individual stress groups showed higher similarity based on the Jaccard index to experimentally-derived GRN from *Arabidopsis* than the GRN based on all data sets (Figure S9B). Furthermore, by taking the union of the stress-specific networks, we obtained a GRN with the highest similarity to the *Arabidopsis* GRN (p-value < 0.05, Figure S9B). Thus, we settled on the union of the seven stress-specific GRN, with R^2^>0.8 performance. Next, to obtain a high-confidence GRN, we selected the transcription factor with the highest absolute coefficient for each gene (Figure 3A). Based on the value of the selected coefficients, the majority of TFs are activators (75 TFs, 3355 positive coefficients, green edges), followed by repressors (19 TFs, 1338 negative coefficients, red edges) and ambiguous (1 TF, 1185 mixture of positive and negative coefficients, gray).

To identify which transcription factors are robustly responding to a given stress, we visualized the significantly up- and down-regulated transcription factors across all available combinations of stresses (Figure 3B). Interestingly, certain transcription factors show consistent expression patterns across most combinations of a given stress group (e.g., *Mp2g00890.1* is down-regulated in 5 out of 6 cold stress combinations, Figure 3B and S10, bottom row). In total, we identified 75 transcription factors that showed a consistent, robust response across >70% of combinations within a stress group (termed robustly-responding TFs). The number of robustly-responding TFs in a stress group corresponds to the number of differentially expressed genes. For example, a large number of robustly-responding TFs are found for stresses with a higher number of DEGs (Figure 3C, darkness, heat), while stresses with few DEGs had fewer robustly-responding TFs (salt, nitrogen deficiency).

We expect that the observed down-regulation of biological processes in darkness should be caused by upregulation of repressors, downregulation of activators, or both. Interestingly, in darkness, the down-regulated TFs comprise mainly of activators (leftmost column, green cells, Figure 3C). In contrast, the up-regulated TFs contained many repressors (red cells), suggesting that the large downregulation of most biological processes is due to the combined action of repressed activators and expressed repressors. Finally, most transcription factors showed specific expression in at most one stress with few exceptions, such as *Mp8g11560* (robustly downregulated in all stresses) and *Mp4g17430* (up-regulated in 5 out of 7 stresses).

### Construction of stress-responsive gene regulatory network

To gain a robust, genome-wide view of the *Marchantia* stress-responsive GRN, we set a global coefficient threshold of the Elastic Net Regression that best explained the observed DEG patterns. To achieve this, we differentiated ’expected’ from ’unexpected’ gene regulatory patterns (see methods). For example, an up-regulated transcriptional activator is expected to up-regulate its target, and conversely, an up-regulated repressor is expected to downregulate its target (Figure 4A). We then set to identify the coefficient that produced the highest ratio of expected / total regulatory edges ranging from 0 (no expected edges are observed) to 1 (all edges are expected). The analysis revealed that at a coefficient cut-off of 0.22, the ratio is highest (48.4% of edges can be explained, Figure S11A), while at the same time, most (89 out of 95) TFs are still connected to other TFs in the GRN (Figure S11B).

The resulting GRN revealed intricate regulatory relationships between the 89 TFs. TFs with the highest number of regulatory targets (dark node color) are heat- and dark-related (indicated by H, D in the node name) (Figure S12). At the same time, these TFs also regulate the highest number of other transcription factors (larger nodes indicate TFs controlling a higher number of other TFs). Interestingly, TFs with the highest number of regulatory targets are typically downregulated (dark nodes with blue borders).

Next, we set out to investigate which biological processes the TFs regulate in the different stresses. We first investigated which biological processes are robustly differentially expressed by finding MapMan bins that show consistent expression pattern changes across the stress combinations (Figure S13). Next, we calculated the percentage of target genes in each MapMan bin that a given TF regulates, based on the above GRN. The number can range from 0 (a TF regulates 0% of genes in a bin) to 1 (a TF regulates 100% of the genes). We set a threshold of 5% target genes in the MapMan bin based on the distribution of the percentages across the network (Figure S14), as at this threshold, the majority of regulatory relationships are expected (e.g., up-regulated activator results in an up-regulated bin, Figure 4B), and most TF-MapMan bin edges are removed (Figure S14), resulting in a sparse network. We observed that multiple transcription factors typically regulate each biological process (Figure S15). For example, the expression of cell wall proteins is decreased in heat (blue node ’Cell wall organisation.cell wall proteins’, Figure 4C), and this biological process is controlled by two down-regulated TFs: *Mp5g01530* and *Mp3g07510* (blue downregulated nodes). At the same time, a TF can regulate multiple biological processes, as exemplified by dark stress-specific downregulated activator *Mp7g09260* downregulating multiple processes related to photosynthesis (Figure 4D). Thus, the inferred GRN can serve as a resource to dissect how *Marchantia* can cope with abiotic stresses.

### Functional comparison of gene expression responses and GRN of Marchantia and *Arabidopsis thaliana*

Often, scientists study model organisms with the hopes of understanding the biology of other species. However, it is currently unclear how conserved the GRNs are between species. If *Arabidopsis* and *Marchantia* show a high degree of GRN conservation, we expect the TF orthologues to show conserved expression patterns and be necessary for survival under the same stresses.

To gauge how similar the stress-specific responses are between *Marchantia* and Arabidopsis, we identified the *Arabidopsis* orthologs of the 95 robustly-responding Marchantia TFs and studied their experimentally-verified biological function and stress-responsive expression patterns. Typically, each *Marchantia* TF has many *Arabidopsis* orthologs due to larger gene families in *Arabidopsis* (Table S7). For most stresses, we did not find visible congruence between the transcriptomic response of *Marchantia* TFs and the biological function and transcriptomic response of the *Arabidopsis* orthologs (Figure 5A, Table S8). For example, the heat-responsive *Marchantia* orthologs in *Arabidopsis* have experimentally-verified biological functions in cold, dark, salt, mannitol/drought, and nitrogen deficiency (Figure 5A) with the majority of functions not being involved in heat stress (Figure 5B). We also observed similar patterns for the gene expression responses of *Arabidopsis* orthologs (Figure 5C), where most observed gene expression responses were not related to heat. Taken together, the minor conservation of biological functions and gene expression responses suggest that stress-responsive GRNs between *Marchantia* and *Arabidopsis* seem poorly conserved.

**Figure 5.**
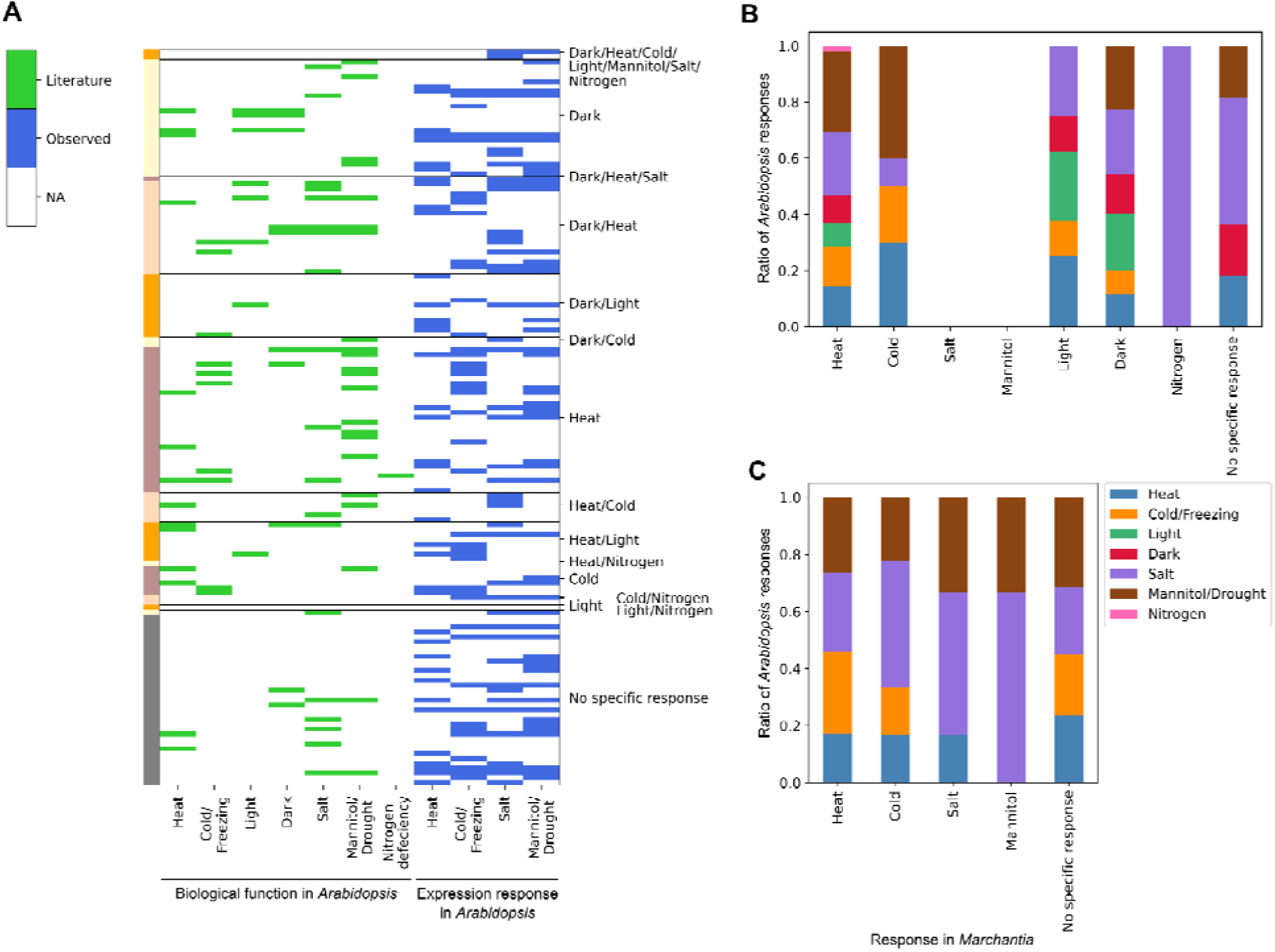
Comparison of stress-specific transcription factors in Marchantia to Arabidopsis orthologs. A) The function of Arabidopsis thaliana orthologs, inferred from the literature (green, obtained from NCBI, Arabidopsis.org) and gene expression responses (blue, eFP browser). Each row contains one Arabidopsis TF, and the rows are grouped and color-coded according to the stress response in Marchantia. The columns indicate the stresses observed in Arabidopsis. B) Ratio of evidence from the literature for *Arabidopsis* transcription factors grouped according to the stress specificity observed in *Marchantia* (x-axis). C) Ratio of evidence by observation of change in expression in *Arabidopsis* transcription factors based on expression data (source eFP browser). High light, dark, an nitrogen deficiency are omitted due to the lack of data. ’No specific response’ comprise transcription factors that were not robustly responding to a stress in Marchantia.

### Regression-based prediction of stress-responsive gene expression

Our analysis of significantly differentially regulated MapMan bins across experiments revealed that specific stresses (e.g., darkness, heat) can dominate other stresses (e.g., salt, mannitol, Figure 2H). This indicates that when two stresses (S_x_ and S_y_) are present, the combined stress S_xy_ may resemble one of the stresses more than the other. However, the rules governing how gene expression values change in a combined stress when two genes are aligned (a gene is up-regulated in S_x_ and S_y_) or conflicting (a gene is up-regulated in S_x_ and downregulated in S_y_), are still unclear.

To better understand the rules governing gene expression in combined stresses, we compared the gene expression change of single stresses and combined stresses. For each of the seven stresses, we identified significantly down- (blue), up-regulated (red), and unchanged genes, resulting in nine possible combinations of S_x_ and S_y_ (Figure 6A). Then, for each combination, we calculated the average log-fold changes in S_x_, S_y,_ and S_xy_. The resulting plot revealed simple near-additive rules governing the gene expression. For example, down-regulated genes in the cold (S_x_ log_2_ fold change - 2.3, Figure 6A, top left corner), when combined with down-regulated genes in all other stresses (S_y_ log_2_ fold change -2.5), result in more negatively downregulated genes in the combined stresses (S_xy_ log_2_ fold change -2.9). This near-additive pattern was seen for all seven stresses for down-regulated (first row) and up-regulated genes (last row, Figure 6A). Interestingly, when S_x_, S_y_ types are conflicting (e.g., S_x_ is up- and S_y_ is down-regulated), S_xy_ shows log_2_ fold change values between the two single stresses, as would be expected from additivity (Figure 6A).

**Figure 6.**
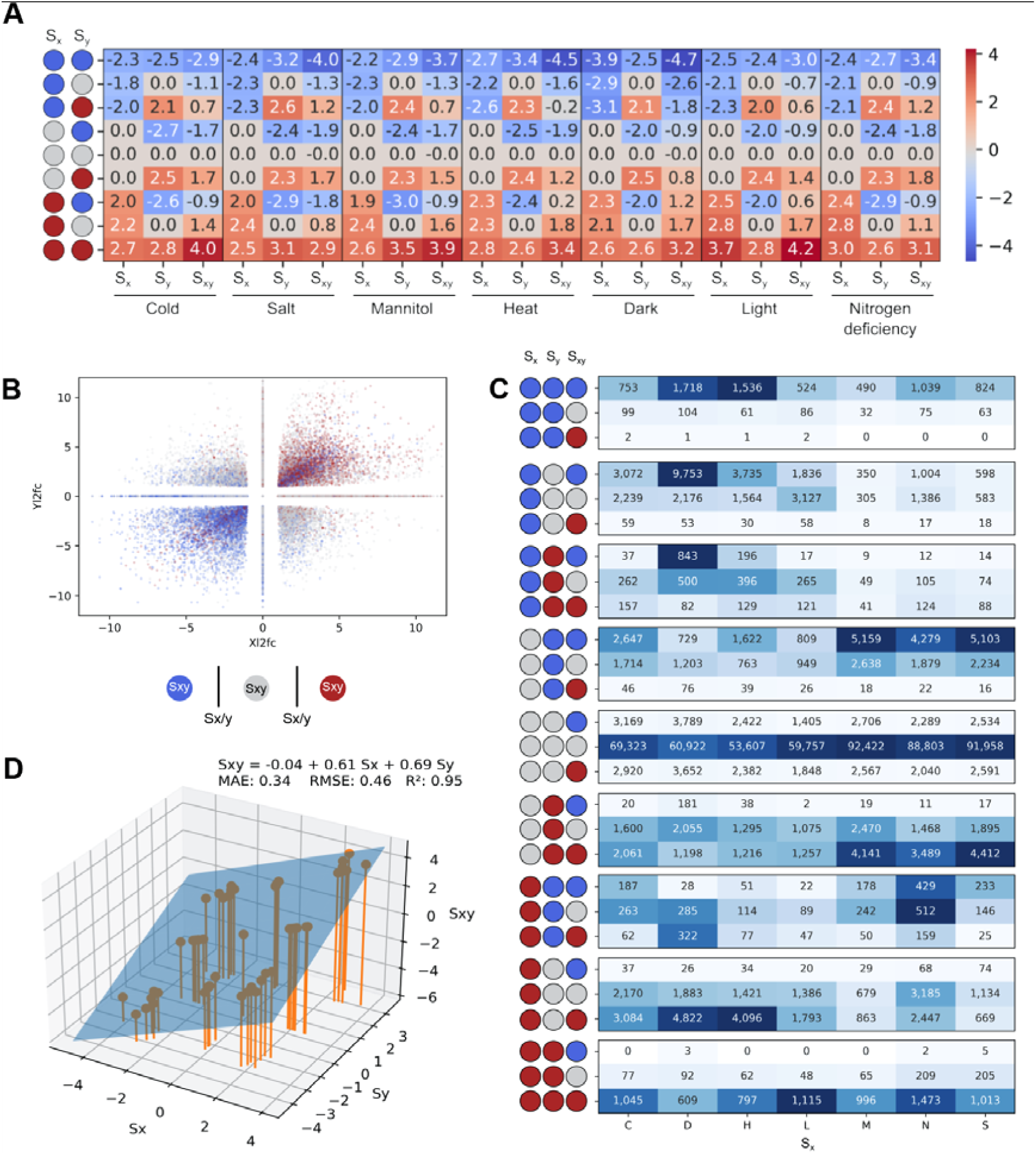
Analysis of gene expression responses in combined stress. A) Averaged log_2_ fold change of genes i S_x_ (type of stress indicated at the bottom), S_y_ (other stresses), and S_xy_ (combined stress). Blue, red, and gray circles indicate genes that are significantly downregulated, up-regulated, and not changed, respectively. B) Scatter plot depicting the outcome in combined stress, where red, blue, and gray dots indicate that the response is higher, lower, or within the range of the single stresses, respectively. C) Break down of the response observed in combined stress S_xy_. The heatmap reflects the proportion of events in given stress (column), and the colors are normalized across each category of S_x_ and S_y_. The actual number of observations is annotated in the cells. Blue, red, and gray circles indicate genes that are significantly downregulated, up-regulated, and not changed, respectively. D) Linear regression of the average log_2_ fold change values from panel A). The inferred formula is shown, together with mean absolute error (MAE), root mean squared error (RMSE), and R^2^ (goodness-of-fit measure).

The near-additive pattern is seen when all possible combinations of S_x_ and S_y_ are color-coded by the S_xy_ outcome (Figure 6B). For example, genes up-regulated in S_x_ and S_y_ tend to be more up-regulated in S_xy_ (Figure 6B, red points in upper right quadrant). Conversely, down-regulation in S_x_ and S_y_ produces an even stronger down-regulation in S_xy_ (Figure 6B, blue points in the upper right quadrant). Conversely, the conflicting log_2_ fold change values tend to produce a response between S_x_ and S_y_ (Figure 6B, gray points). Differential gene expression analysis of S_x_, S_y,_ and S_xy_ follow similar patterns, where down-regulated S_x_ and S_y_ almost exclusively result in downregulated S_xy_ (Figure 6C, top row).

Since the rules of how the different stress responses seem to follow a simple additive pattern, we investigated whether we can explain the average log_2_ fold change S_xy_ observed in Figure 6A by regressing it on the average log_2_ fold change S_x_ and S_y_. We found that the model S_xy_ = -0.04 + 0.61*S_x_ + 0.69*S_y_ can excellently explain the average log_2_ fold change (R^2^ = 0.95, Figure 6D), suggesting a simple linear mechanism of integrating gene expression changes. To further examine how well the different stresses can be explained by the log_2_ fold change values of the individual genes, rather than averages, we performed another 3-dimensional linear regression (Figure 7A-G). While the R^2^ values dropped, the resulting models could still approximate the expected values well (R^2^ = 0.57-0.67).

**Figure 7.**
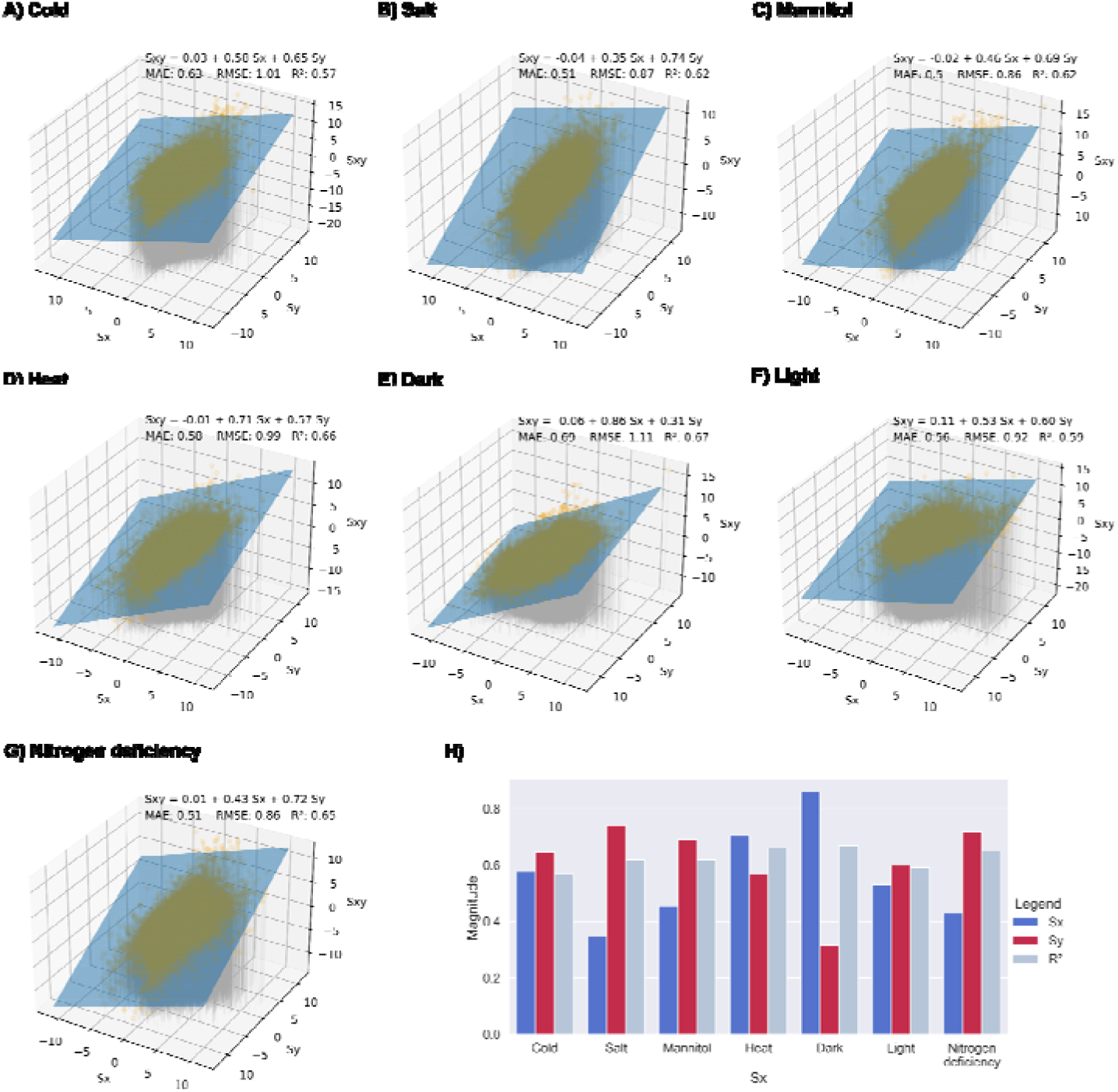
Stress-specific linear regression analysis. S_x_ indicates log_2_ fold change from the specific stress, S_y_ indicates log_2_ fold change from all other stresses, and S_xy_ represents the log_2_ fold change from the combined stresses. The S_x_ stresses are A) Cold, B) Salt, C) Mannitol, D) Heat, E) Dark, F) Light, and G) Nitrogen deficiency. H) Summary of the linear regressions coefficients and model prediction quality. The blue and red bars indicate the S_x_ and S_y_ coefficients, respectively, while the light blue bar indicates the R^2^ value.

Interestingly, we observed that the parameters S_x_ of S_y_ values differed between stresses (Figure 7H). For example, the S_x_ parameter (reflecting log_2_ fold change from the darkness experiment) in darkness is larger (0.86) than the S_Y_ parameter (0.31, capturing log_2_ fold change from all non-darkness experiments), indicating that gene expression differences resulting from the darkness have a higher influence on gene expression than other stresses, which is in line with above results (Figure 2H). Based on the S_x_ coefficients, we can rank the dominance of stresses as darkness (0.86) > heat (0.71) > cold (0.58) > light (0.53) > mannitol (0.46) > nitrogen deficiency (0.43) > salt (0.35).

### eFP browser and CoNekT database for Marchantia

Bioinformatic data is only as useful as its accessibility. To make our data readily accessible, we have constructed an eFP browser instance for *Marchantia* available at https://bar.utoronto.ca/efp_marchantia/cgi-bin/efpWeb.cgi ^37^, and updated our CoNekT database ^39^, with our stress data, available at https://conekt.sbs.ntu.edu.sg/. To exemplify how our data and these databases can be used, we provide an example with phenylpropanoids, which contribute to all aspects of plant responses towards biotic and abiotic stimuli. In flowering plants, phenylpropanoids were found to be highly responsive to light or mineral treatment, important for resistance to pests ^40^, and invasion of new habitats and reproduction ^41^. The biosynthesis of phenylpropanoids begins with phenylalanine ammonia lyases (PAL) and tyrosine ammonia lyases (TAL) that catalyze the non-oxidative deamination of phenylalanine to trans-cinnamate, which directs the output from the shikimate pathway phenylpropanoid metabolism ^40^.

To study phenylpropanoid metabolism in *Marchantia*, we started by entering ’PAL1’ into CoNekT’s search box (https://conekt.sbs.ntu.edu.sg/), which took us to the page of PAL1, a PAL gene *AT2G37040* from *Arabidopsis thaliana*. To identify PAL genes in Marchantia, we clicked on the link of the Land Plants orthogroup (CoNekT provides orthogroups for Archaeplastida, Land Plants, and Seed Plants), which revealed that all 11 land plants in the database contain PALs, while the algae *Cyanophora paradoxa* and *Chlamydomonas reinhardtii* do not. *Marchantia* contains a surprisingly large number (13) of PALs given the low redundancy of its genome, which is higher than *Arabidopsis* (4). CoNekT contains gene trees that also show the expression of genes in major organ types. The analysis revealed that the many PAL genes in Marchantia likely result from a recent duplication within Marchantia (Figure S16).

To gain insight into the function of the 13 PAL genes from *Marchantia*, we set out to study the expression of the PALs during stress conditions. First, we copied the gene identifiers into Tools/Create heatmap (https://conekt.sbs.ntu.edu.sg/heatmap/), revealing that most PALs show a high expression during cold treatment combined with nitrogen starvation (Figure 8A). Then, we clicked on one of the highly responsive genes *Mp4g14110.1*, which took us to the page dedicated to the gene. The page contains various information, such as gene description, CDS/protein sequences, gene families, found protein domains, co-expression networks, and others. The detailed expression profile confirmed the high cold + nitrogen-specific expression of *Mp4g14110* by using the CoNekT expression profile plot (Figure 8B) and eFP viewer (Figure 8C, red square at the intersection of cold and nitrogen deficiency). To better understand the function of *Mp4g14110*, we clicked on the cluster link that directed us to the co-expression cluster link of *Mp4g14110*. The cluster page contains information about the genes found in the cluster, Gene Ontology enrichment analysis, found protein domains, and other functional information.

**Figure 8.**
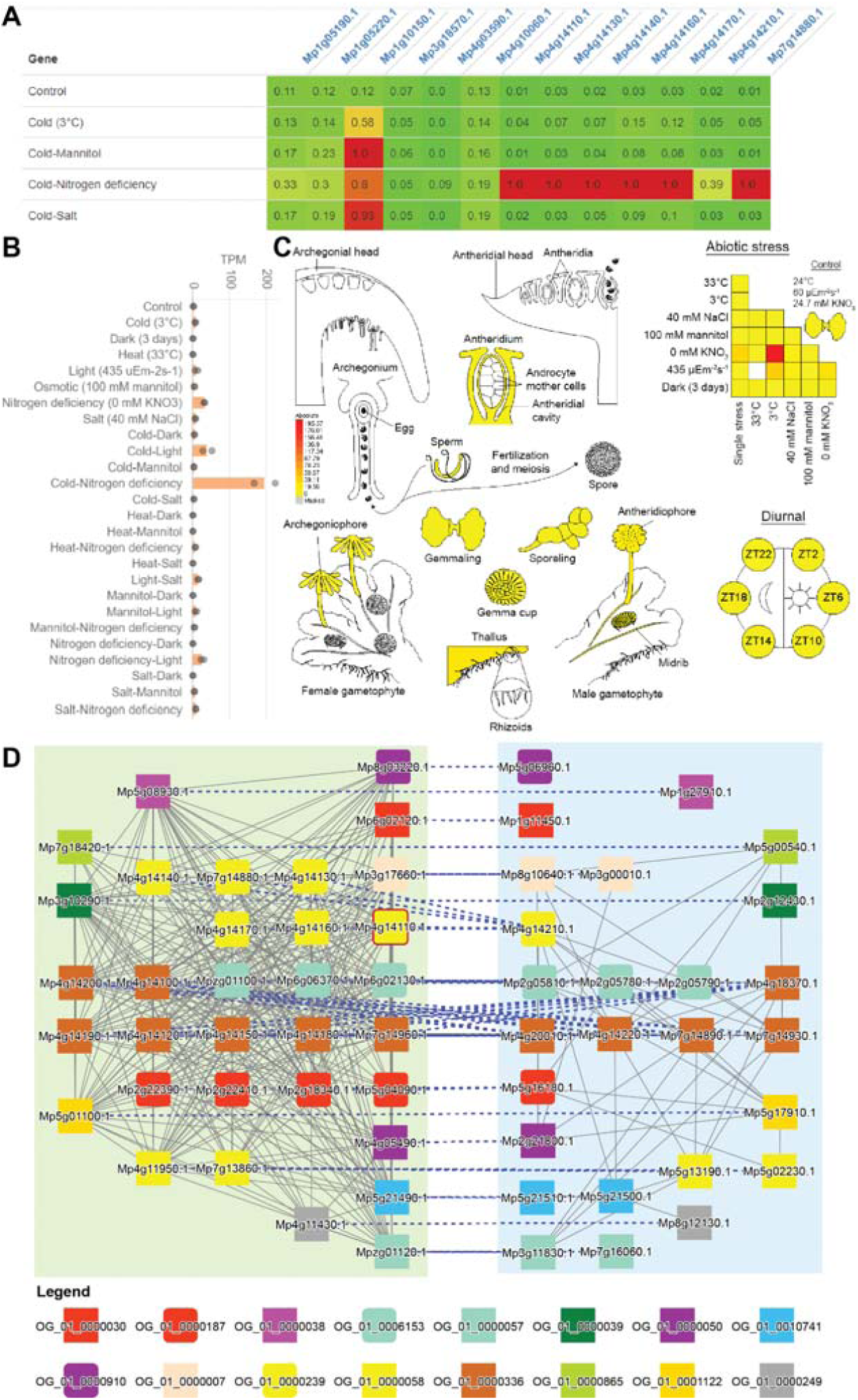
Implementation of the *Marchantia* gene expression data in CoNekT and eFP browser. A) Heatmap showing expression of PALs in representative stresses. B) Expression profile of Mp4g14110.1 (*MpPAL7*) under single and combined stress in CoNekt. C) Expression of *MpPAL7* in different organs and under different abiotic stress conditions in eFP. D) Comparison of Cluster 102 and 50 bounded by green and blue boxes, respectively. The border of node MpPAL7 is colored red, and blue dashed lines indicate homology between genes from the two clusters. A) Heatmap showing expression of PALs in representative stresses. B) Expression profile of Mp4g14110.1 (*MpPAL7*) under single and combined stress in CoNekt. C) Expression of *MpPAL7* in different organs and under different abiotic stress conditions in eFP. D) Comparison of Cluster 102 and 50 bounded by green and blue boxes, respectively. The border of node MpPAL7 is colored red, and blue dashed lines indicate homology between genes from the two clusters.

Interestingly, the ’Similar Clusters’ cluster page revealed similar co-expression clusters in other species (similarity is based on ortholog membership and defined by Jaccard Index), with another similar cluster in Marchantia. To study these duplicated clusters, we clicked on the ’Compare’ button, which revealed that the two clusters contain several gene families involved in phenylpropanoid biosynthesis (yellow rounded rectangles: PALs, brown rectangles: chalcone synthases), ABC and DTX transporters implicated in metabolite transport across membranes (purple/gray/green and red rectangles ^42^), auresidin synthases that can hydroxylate or cyclize chalcones (light blue rounded rectangles ^43^) and glutathione transferases (red rounded square ^44^) (Figure 8D). Interestingly, both clusters contained WRKY transcription factors (salmon rectangles), implicating these transcription factors in controlling the biosynthesis of the respective metabolites.

Taken together, our tools revealed evidence of duplicated modules, likely involved in the biosynthesis of related phenylpropanoids. The updated CoNekT platform contains many additional tools to predict gene function and find relevant genes ^16, 28, 39, 45^. For example, the tool found many of the described genes by identifying genes highly expressed during combined cold and nitrogen starvation (Table S9), by clicking on Tools/Search Specific Profiles, selecting Marchantia and ’Cold-Nitrogen deficiency’.

## Discussion

Plants are often exposed to multiple abiotic stresses, which requires them to perceive and integrate multiple signals and respond in a manner that allows them to survive. To understand how plants integrate and respond to the various environmental cues, we constructed a stress expression atlas capturing gene expression changes to single and combined stresses for *Marchantia polymorpha*.

To determine the response of *M. polymorpha* towards various single stresses, we tested a range of severity for heat, cold, salt, osmotic, light, dark, and nitrogen deficiency on *M. polymorpha* gemmae. As expected in most stresses, the size of the plants decreased proportionally to the severity of stresses (Figure 1B). Specific stresses, such as 3°C cold and 535 uEm^-2^s^-1^ light, caused only minor growth phenotypes, suggesting that *Marchantia* can survive under even lower temperatures and higher light intensities. Conversely, stresses such as osmotic (100mM mannitol), salt (40mM NaCl), and carbon starvation (3 days of darkness) produce strong growth phenotypes resulting in reduced growth (osmotic, salt, darkness) and discoloration (yellow-green for mannitol > 150mM, darker plants for salt > 60mM salt) (Figure 1B). Death of plants occurred for heat stress at 36°C for 24 hours and at higher salt concentrations (>200mM). Interestingly, the plants grew at 0mM KNO_3_, albeit slower and with reddish discoloration likely caused by auronidin, a flavonoid shown to accumulate during phosphate deficiency ^38^.

In most cases, the combination of two stresses resulted in additive phenotypes (e.g., salt+mannitol stress results in smaller plants than the two stresses separately, Figure 1C-E). While heat (33°C) combined with high light intensity (435 uEm^-2^s^-1^) resulted in death, this was caused by temperature built up in the sealed plates, causing the temperature to rise to the lethal 38°C. Consequently, this stress combination should be performed using an open plate setup in the future. The only exception to this was salt+nitrogen deficiency (SN) (40 mM NaCl, 0 mM KNO_3_), which was significantly larger than plants exposed only to salt stress and was observed to have extensive and dense rhizoids (Supplemental Data 1). Curiously, we did not observe any dramatic phenotypes when combining carbon/energy starvation (3 days of darkness) with any other stresses (Figure 1C). This is counterintuitive as, e.g. heat stress acclimation is a costly process requiring the biosynthesis of new transcripts and proteins to repair, replace and rebalance the affected cellular machinery ^46^. A hypothesis for the unexpectedly mild phenotype for combined dark stress could be due to the lack of photosynthetic processes in the absence of light, which increases the plant’s capacity to cope with increased ROS, a typical response to most abiotic stresses ^47^. This is also exemplified by the significant downregulation of oxidoreductases and chloroplast redox homeostasis in combined dark stress (Figure 2H). These phenotypes demonstrate that Marchantia is able to survive various adverse growth conditions and can serve as an excellent model for studying stress acclimation.

Interestingly, purely physical stresses (cold, heat, darkness, and high light) showed a higher number of DEGs than chemical stresses (salt, mannitol, and nitrogen deficiency) (Figure 2A). We speculate that the physical stresses cause more DEGs because these stresses can permeate every cell, affect every protein (heat, darkness), and/or dramatically affect the energy levels that have consequences on all processes (darkness, high light). Conversely, stresses such as mannitol and salt can be effectively contained by the action of ion transporters and osmolyte accumulation ^48^. Interestingly, we did not observe any correlation between the number of DEGs and the effect on plant growth (Figure 2B); while salt and mannitol treatments caused the most dramatic growth defects, these two stresses also showed the lowest number of DEGs (Figure 2B). This observation is in line with our study on alga *Cyanophora* ^16^, suggesting that there is no correlation between a visible phenotype and growth across stresses in other plants.

We observed that certain stresses dominate the transcriptional responses. For example, darkness+cold looks more like darkness than cold (Figure 2H), suggesting that darkness is the dominant stress (Figure 2F, G, dominance plots are red in darkness stresses). The analysis of differentially expressed pathways allowed us to rank the strength of dominance of abiotic stresses: darkness (clustered in six stress combinations), then heat and light (four combinations each), then nitrogen deficiency and cold (three each)(Figure 2H). To better understand the mechanism governing the dominance of the stresses, we performed several analyses that revealed multiple mechanisms that likely work together.

Firstly, we identified 75 robustly-responding TFs, revealing that the dominant stresses (e.g., heat and darkness) differentially express a higher number of TFs than the non-dominant stresses (Figure 3A). Secondly, the inferred GRN showed that the TFs active in the dominant stresses regulate more genes and other TFs than TFs from non-dominant stresses (Figure 4A, H, D nodes tend to be darker). Thirdly, our regression model showed that the gene expression changes in a combination of two stresses could be explained by the addition of the log_2_ fold change values. Thus, when combining stress with a high number of largely negative log_2_ fold change values (e.g., darkness down-regulated log_2_ fold change is between -3.9 to -2.9, Figure 6A, top row), with other stresses, the log_2_ fold change in combined stress will also be negative (log_2_ fold change for combined stress in darkness down-regulated genes ranges between - 4.7 to -1.8, Figure 6A). Fourth, our regression model showed that dominant stresses have higher coefficients (Figure 7H), suggesting yet another unknown component governing the integration of multiple stress responses.

Our comparison of Marchantia robustly responding TFs and their Arabidopsis orthologs revealed that the majority of Arabidopsis TFs are not involved in the same stresses as the *Marchantia* TFs (Figure 5AC). This contrasts with studies showing conservation of stress response in plants focusing on transcription factors and hormones ^49–53^. However, the lack of conservation between Marchantia and Arabidopsis is not entirely unexpected, as massive changes such as genome and gene duplications have occurred since the last common ancestor, rendering a lack of homology in genes across species and differences in gene families and regulation. This suggests that each model plant uses a different strategy to acclimate to stress, making it uncertain to what degree knowledge gained from model species such as *Arabidopsis* can be used to improve our crops. This lack of conservation of responses to abiotic stresses has been observed by us at the species level in *Cyanophora* ^16^, and at the intraspecies level by a salt stress study in six Lotus accessions, where only 1% of genes showed a conserved response ^54^, in seven *Arabidopsis* accessions, which showed a divergent response to treatment by salicylic acid ^55^ and by two strawberry cultivars, which displayed modest conservation of DEGs to the same pathogen ^56^. However, while we analyzed only gene expression, stress responses can be active at the epigenetic (methylation of genes), transcriptomic (mRNA, microRNA, lncRNA) and proteomic (posttranslational modifications and activation) levels ^57–61^.

To make our data more readily accessible, we provide an eFP browser for Marchantia, a popular tool allowing the visualization of gene expression by an ’electronic Fluorescent Pictograph’. Furthermore, we provide an updated CoNekT database with expression atlases of 13 species comprising various algae and land plants. The database provides tools to study gene expression, functional enrichment analyses of co-expression networks, and other comparative tools. These valuable tools will help further dissect the gene regulatory networks behind abiotic stress responses in *Marchantia* and other species Figure 7).

Importantly, our analysis shows that it is possible to predict gene expression of combined stresses with a simple linear regression. This paves the way to building more complex, better-performing models that can predict gene expression in any environment, given sufficient input data. This strengthens the call for more emphasis on studying combined biotic and abiotic stresses in light of future challenges posed by climate change ^62, 63^.

## Acknowledgments

We want to thank the Nanyang Technological University start-up grant for funding. We want to thank the members of the Mutwil lab for their comments on the manuscript. We thank D. Maizels (http://www.scientific-art.com/) for the illustrations used for the eFP browser.

## Author contributions

M.M. conceived the project, wrote the manuscript, Q.W.T. performed the single stress experiments and data analysis, updated the CoNekT database and wrote the manuscript, P.K.L performed the combined stress experiments, N.P. and A.P. provided the eFP browser, Z.C., provided the Marchantia cultures and feedback on growing Marchantia. M.A. provided scripts for the ElasticNet regression, while Z.N. supervised M.A. and Q.W. in ElasticNet regression. Z.N. also contributed to the manuscript writing and project design.

## Supplementary figures

**Figure S1.**
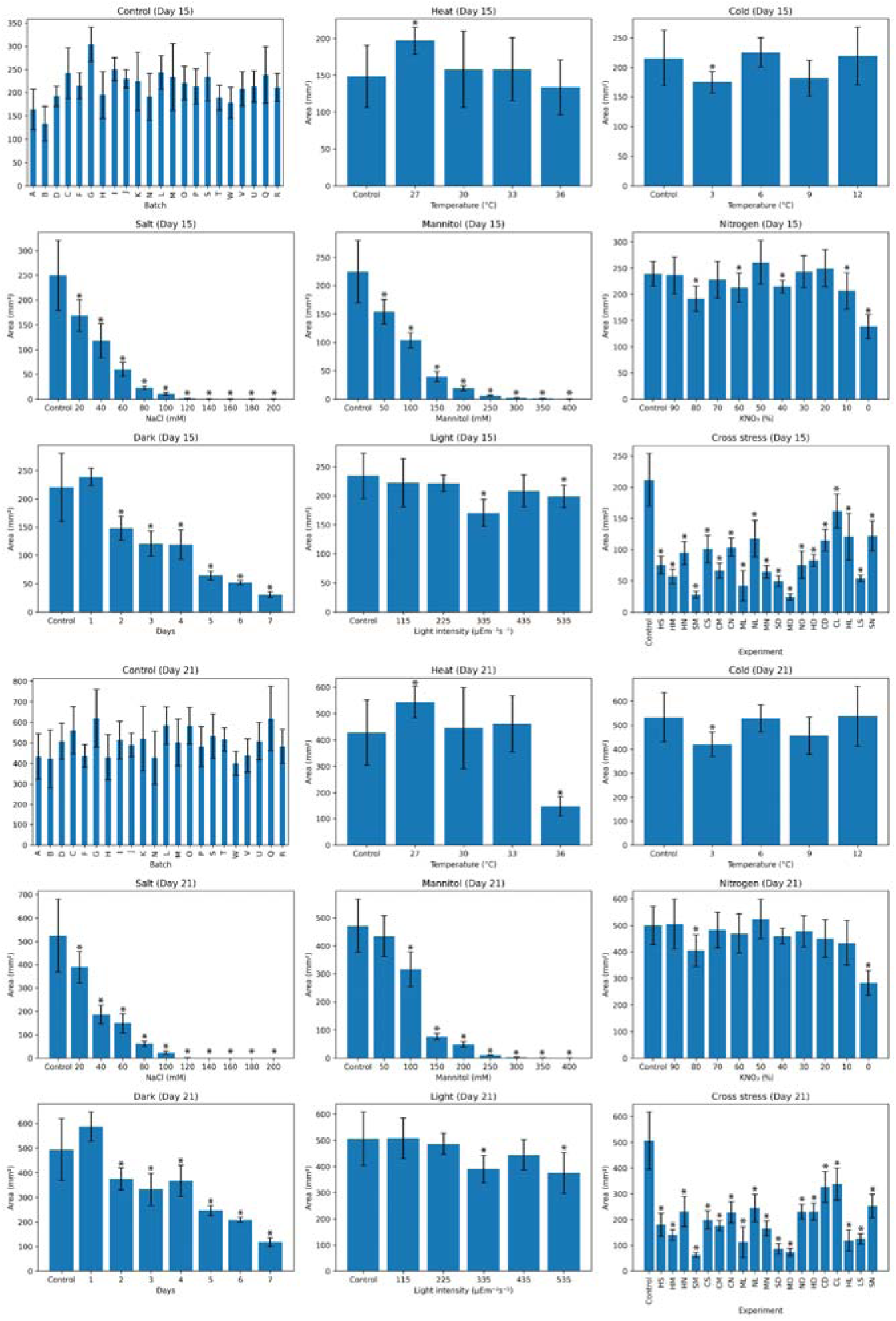
Area of plants at 15 days (day of harvest) and day 21 (6 days post-harvest) for the single and combined stresses. Each bar represents measurements from at least 4 plants. Error bars are represented by standard deviation and significance was determined by Student’s two-tailed, t-test, p < 0.05.

**Figure S2.**
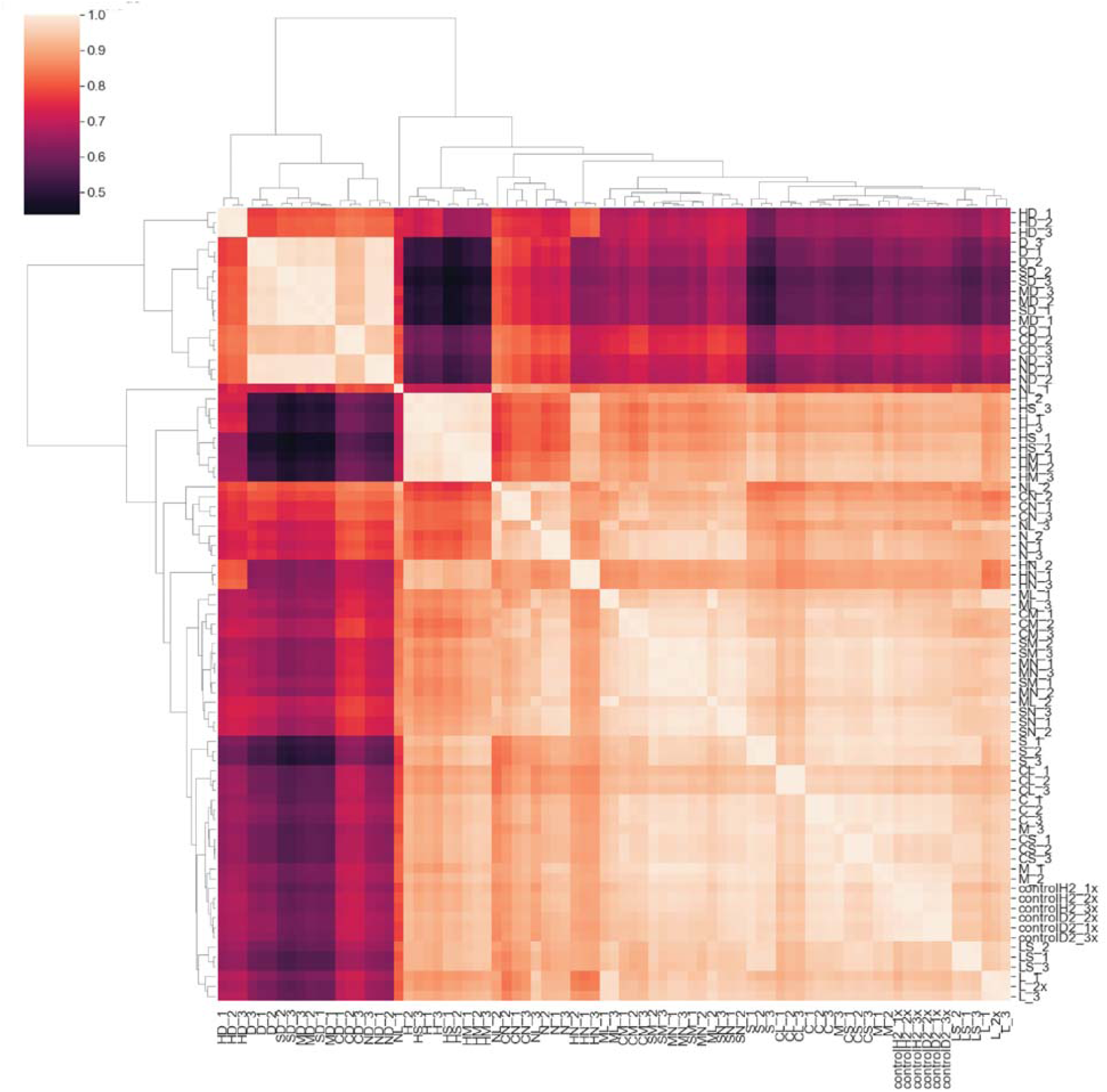
Hierarchical clustering of the Marchantia stress experiments. Transcript per million (TPM) gene expression values were scaled with standard scaler and clustered with seaborn.clustermap() function.

**Figure S3.**
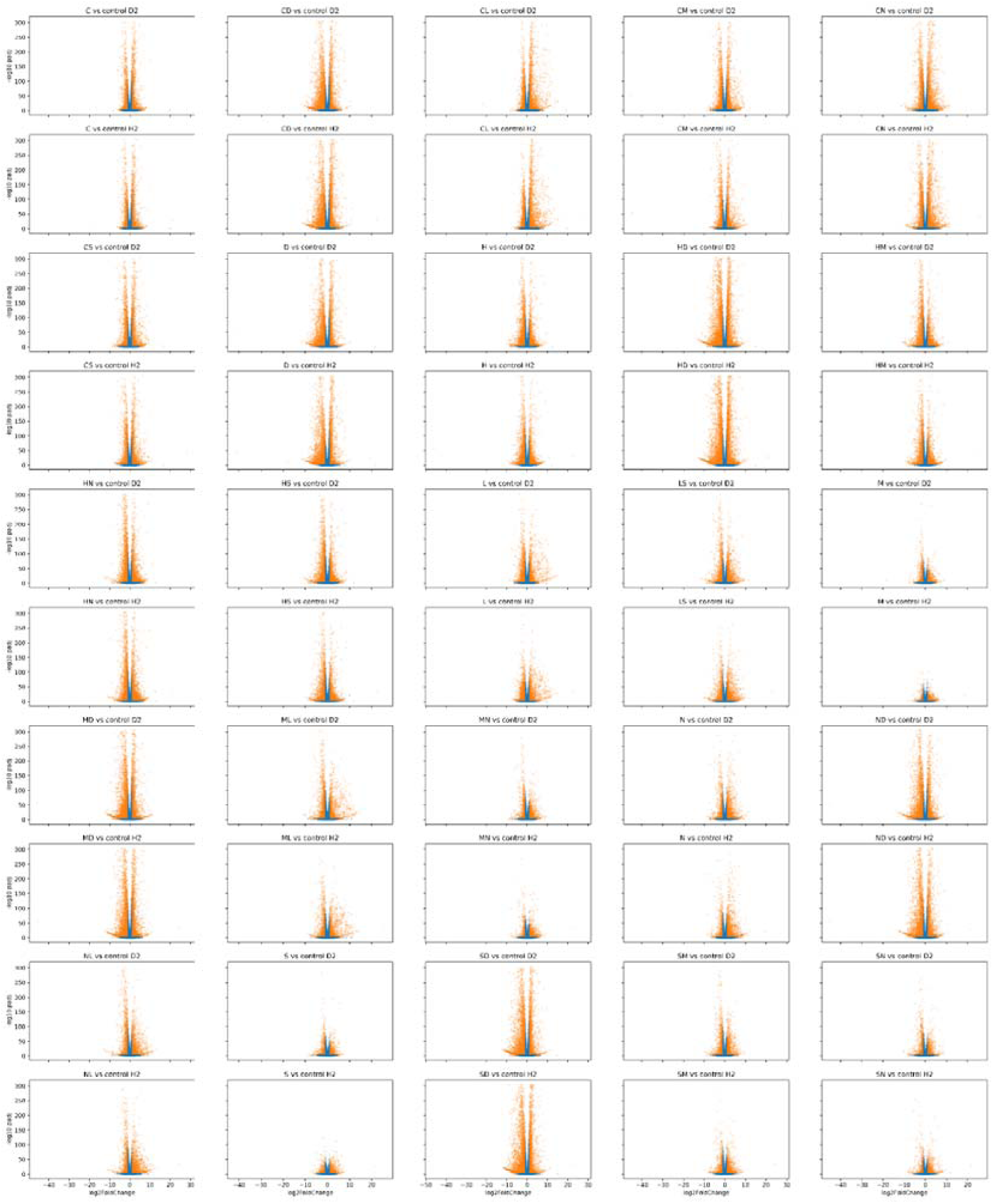
Volcano plots of stress experiments. The x-axis represents log_2_-fold change, the y-axis indicates the -log_10_ of the adjusted p-value. Each point represents a gene that has an absolute log_2_-fold change > 1 and is significantly (adjusted p-value<0.05, orange color) or not significantly (p-value>0.05, blue point) differentially expressed.

**Figure S4.**
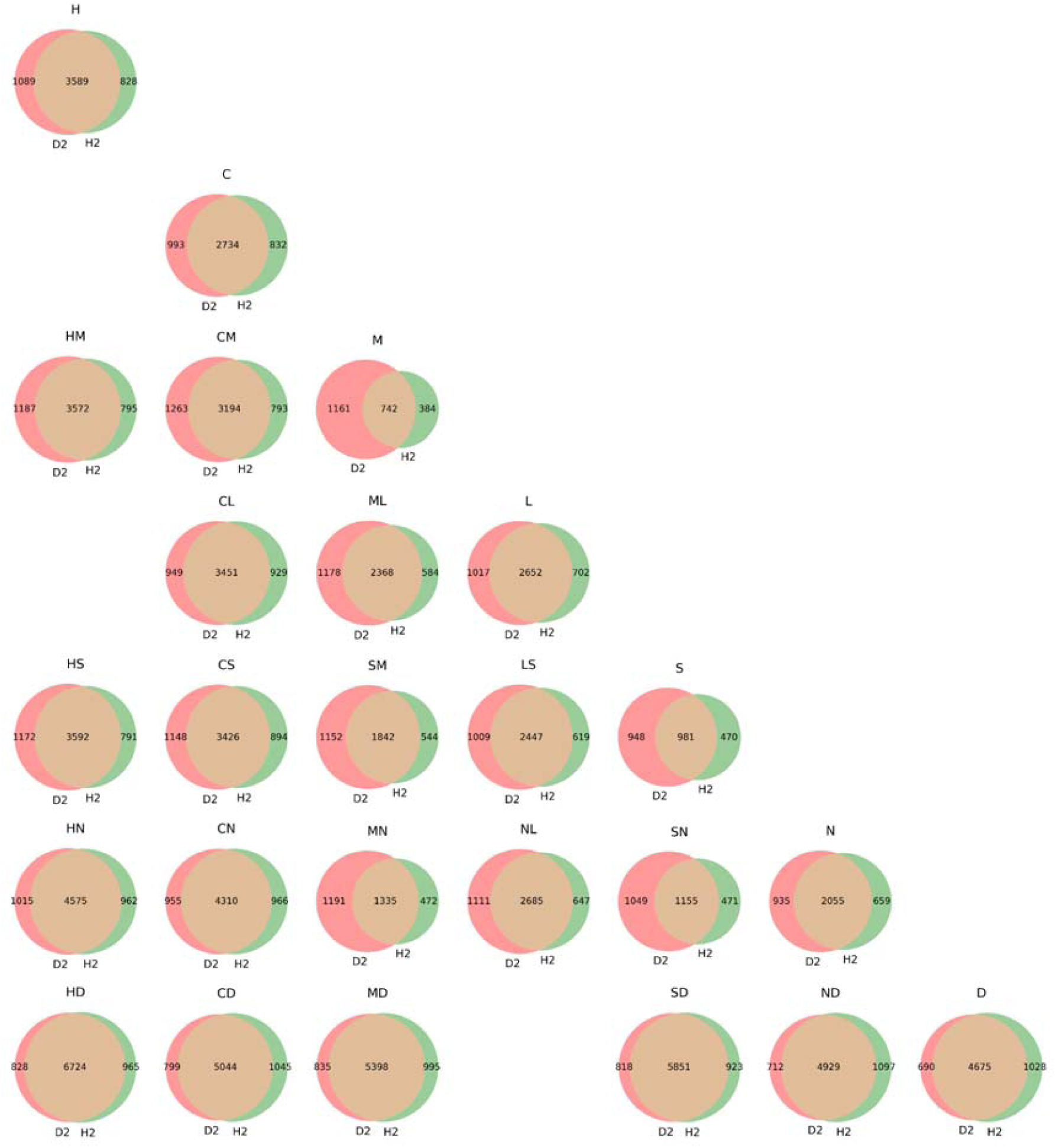
Comparison of found differentially expressed genes for the single and combined stressed. For each stress combination, we estimated significant DEGs (adjusted p-value<0.05) by using control D2 (red circle) and control H2 (green circle). The orange areas indicate the overlap between the found DEGs of the two controls. The overlapping DEGs were used for further study. The following abbreviations are used for the description of stresses: cold (C), dark (D), heat (H), light (L), mannitol (M), nitrogen deficiency (N), and salt (S). The combination of stresses is indicated by two letters, where e.g., HD indicates heat+darkness treatment.

**Figure S5.**
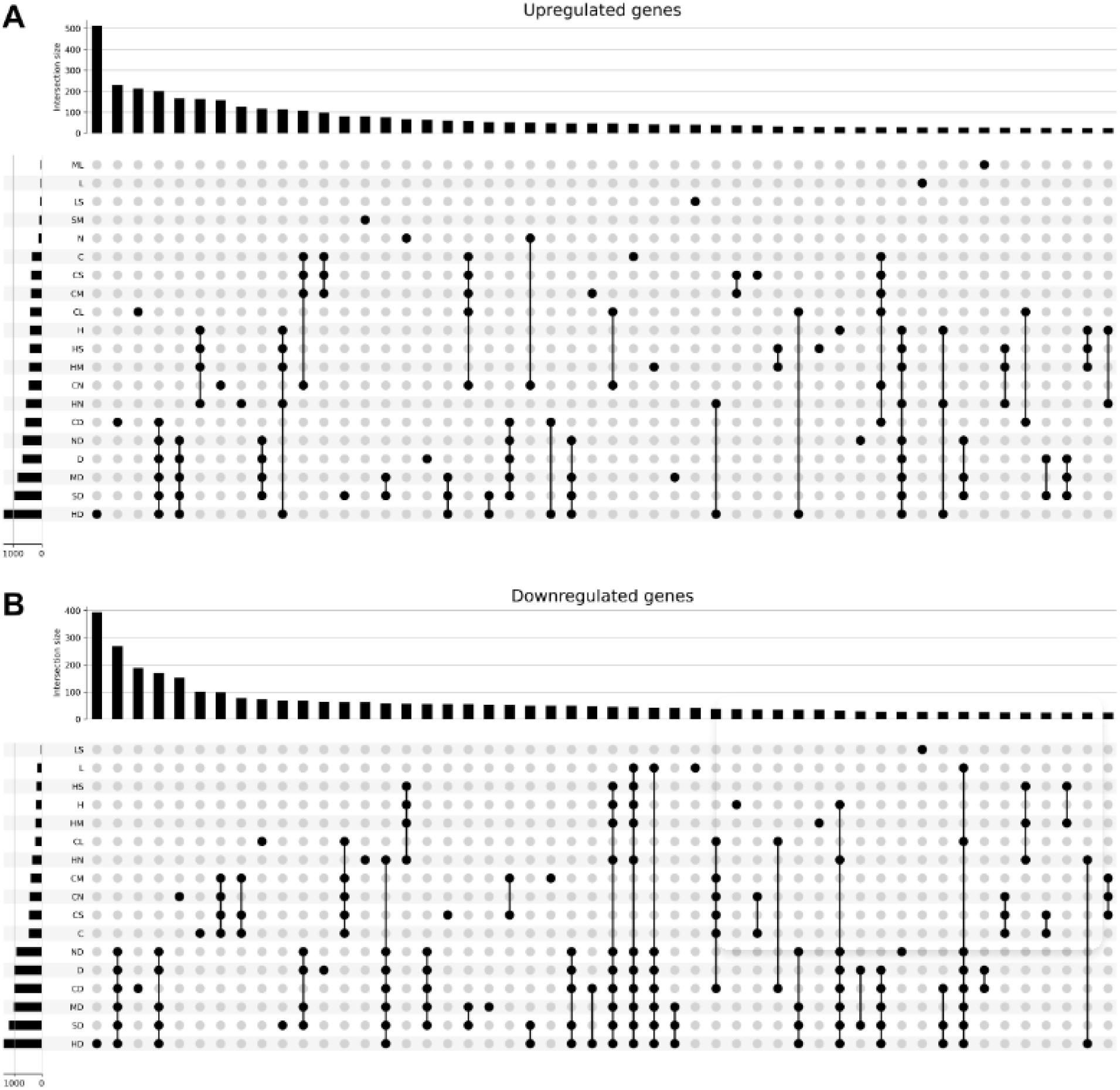
Upset plot showing the top 50 intersections of differentially expressed genes. A) Up-regulated genes. B) Downregulated genes. The package is available from htps://upsetplot.readthedocs.io/en/stable

**Figure S6.**
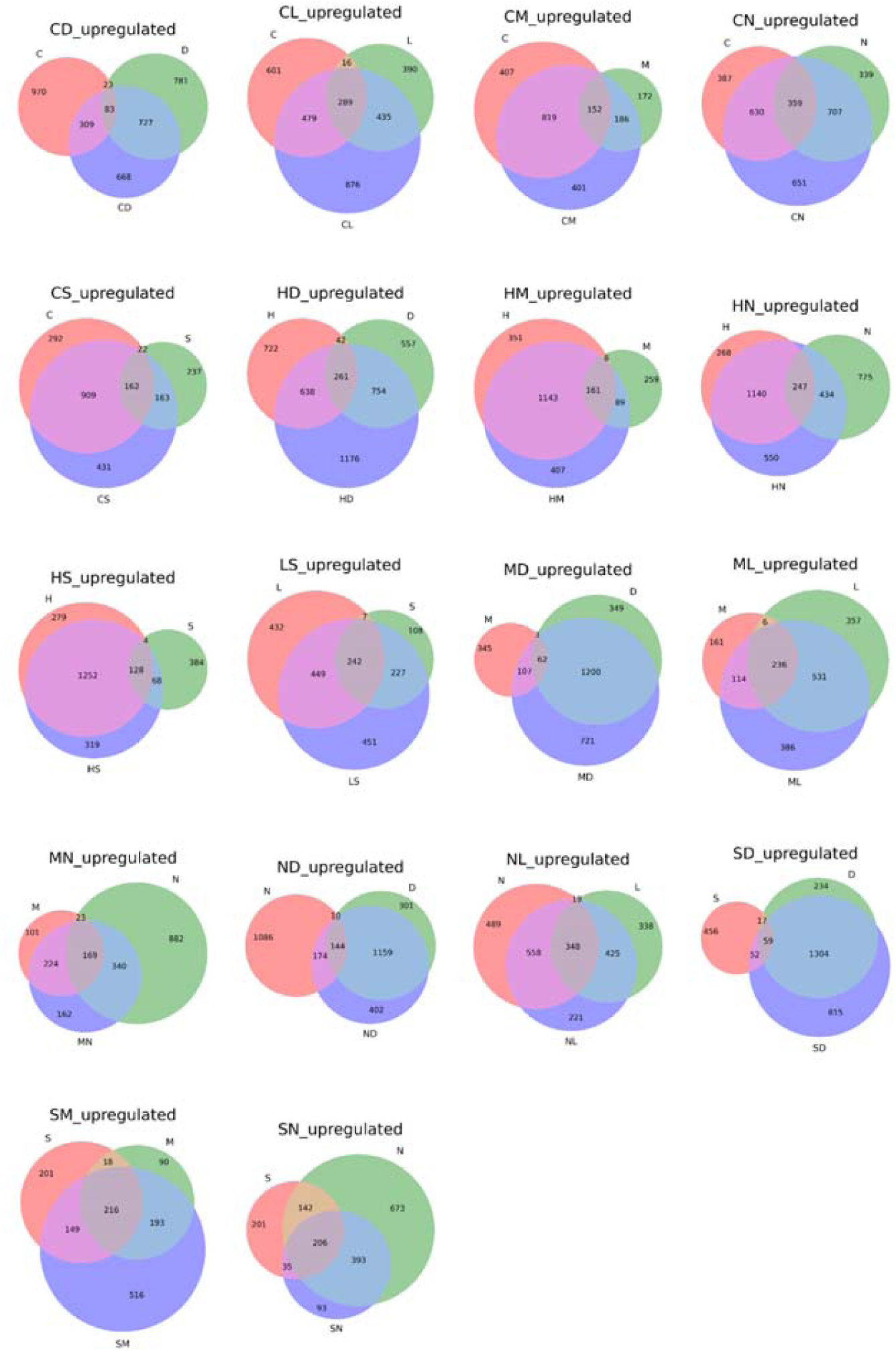
Venn diagrams of up-regulated genes in single and combined stresses. The abbreviations are cold (C), dark (D), heat (H), light (L), mannitol (M), nitrogen deficiency (N), and salt (S). A combination of stresses is indicated by two letters and purple circles. The first and second stress in a pair is colored red and green (e.g., cold is red in CD), while the combined stress is colored purple. The sizes of the circles and intersections and the numbers within indicate the number of significant DEGs.

**Figure S7.**
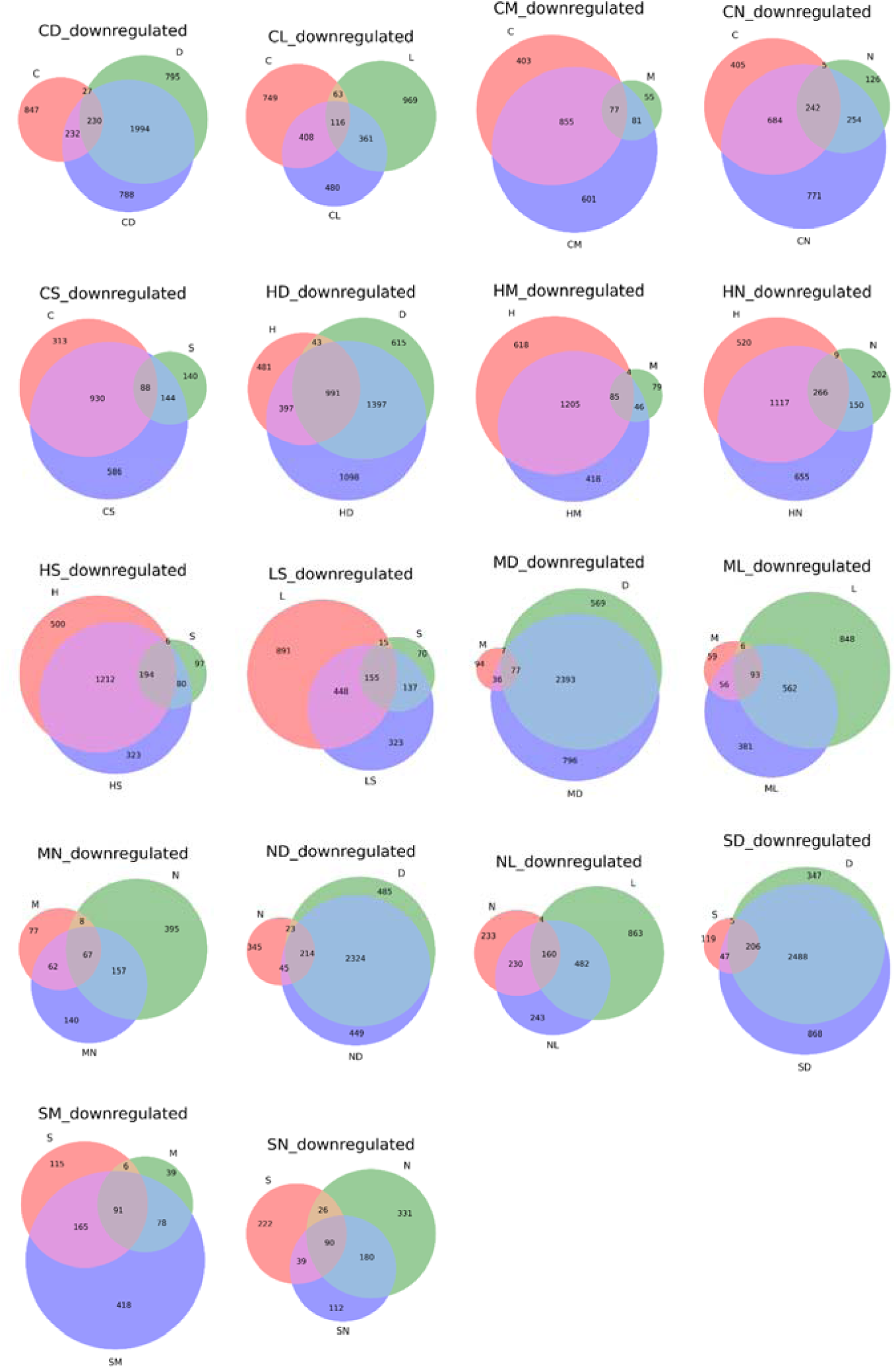
Venn diagrams of downregulated genes in single and combined stresses. The abbreviations are cold (C), dark (D), heat (H), light (L), mannitol (M), nitrogen deficiency (N), and salt (S). A combination of stresses is indicated by two letters and purple circles. The first and second stress in a pair is colored red and green (e.g., cold is red in CD), while the combined stress is colored purple. The sizes of the circles and intersections and the numbers within indicate the number of significant DEGs.

**Figure S8.**
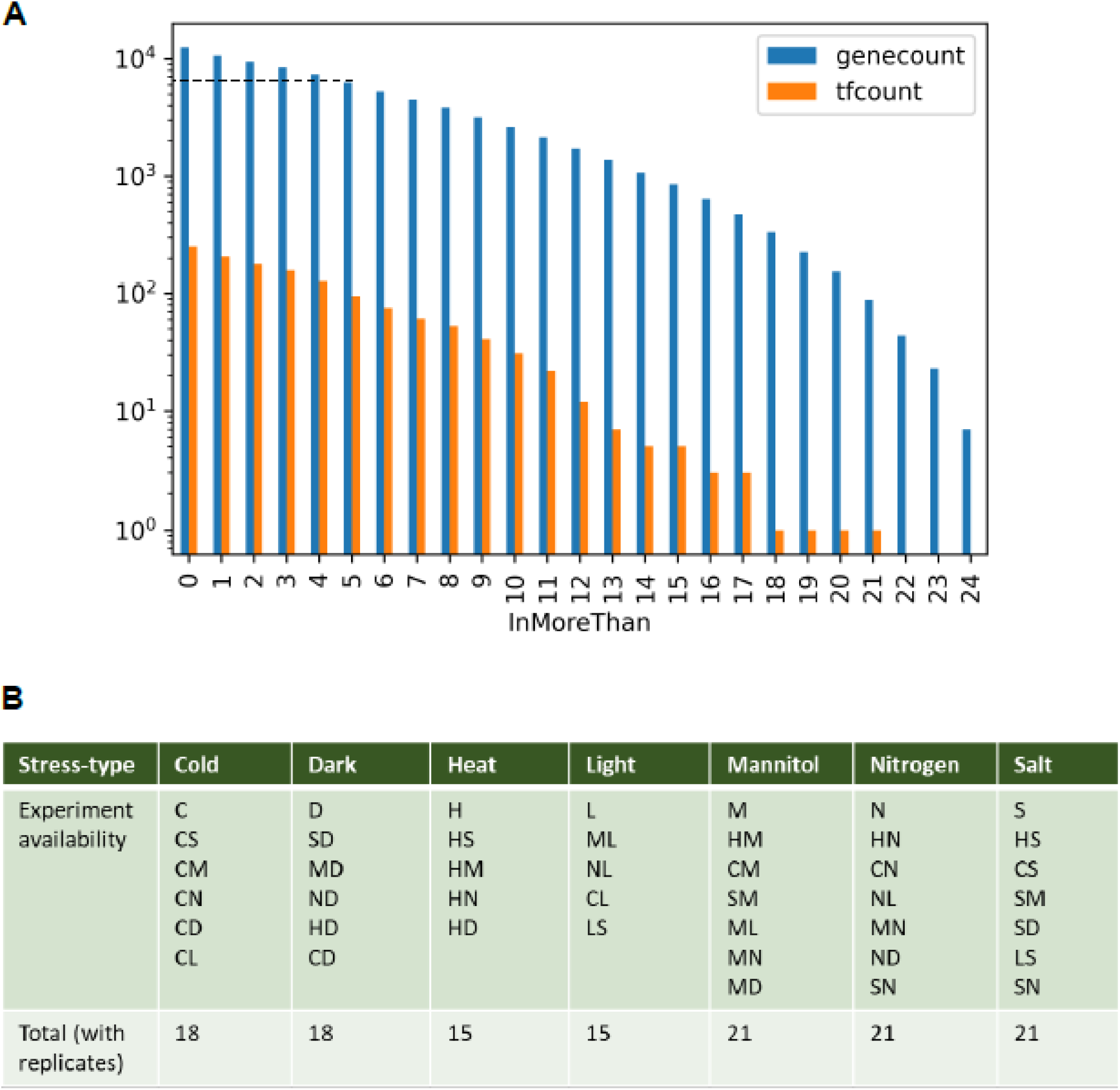
Considerations for the reconstruction of the stress-specific gene regulatory networks. A) Number of differential genes in more than a certain number of experiments. B) List of experiments in each stress specific-network.

**Figure S9.**
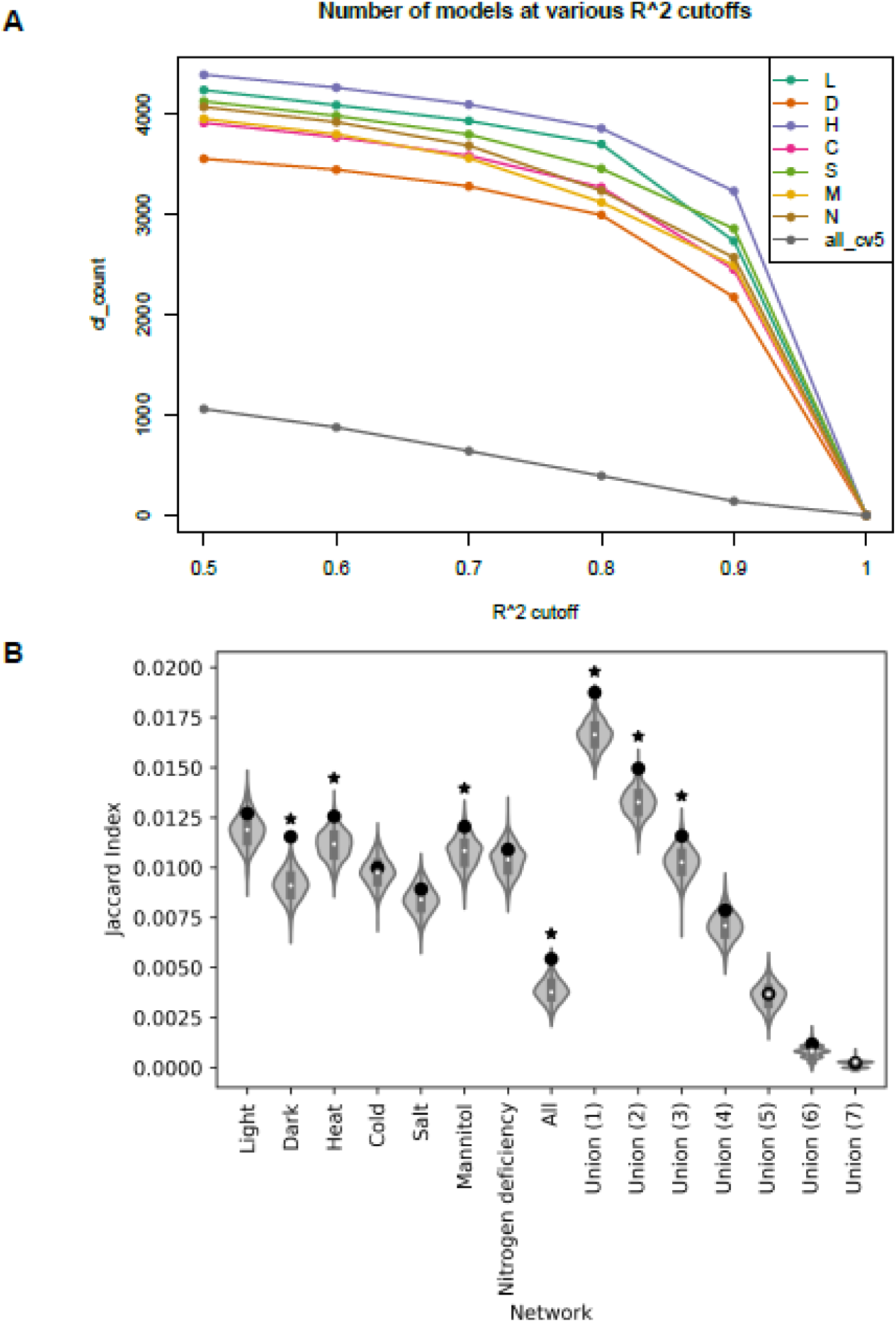
Metrics used to construct the unified GRN. A) Distribution of model counts across various R^2^. B) Similarity of various gene regulatory networks against a shuffled AGRIS gene regulatory network. Point indicates observed similarity while asterisk indicates significance, hypergeometric test, p < 0.05.

**Figure S10.**
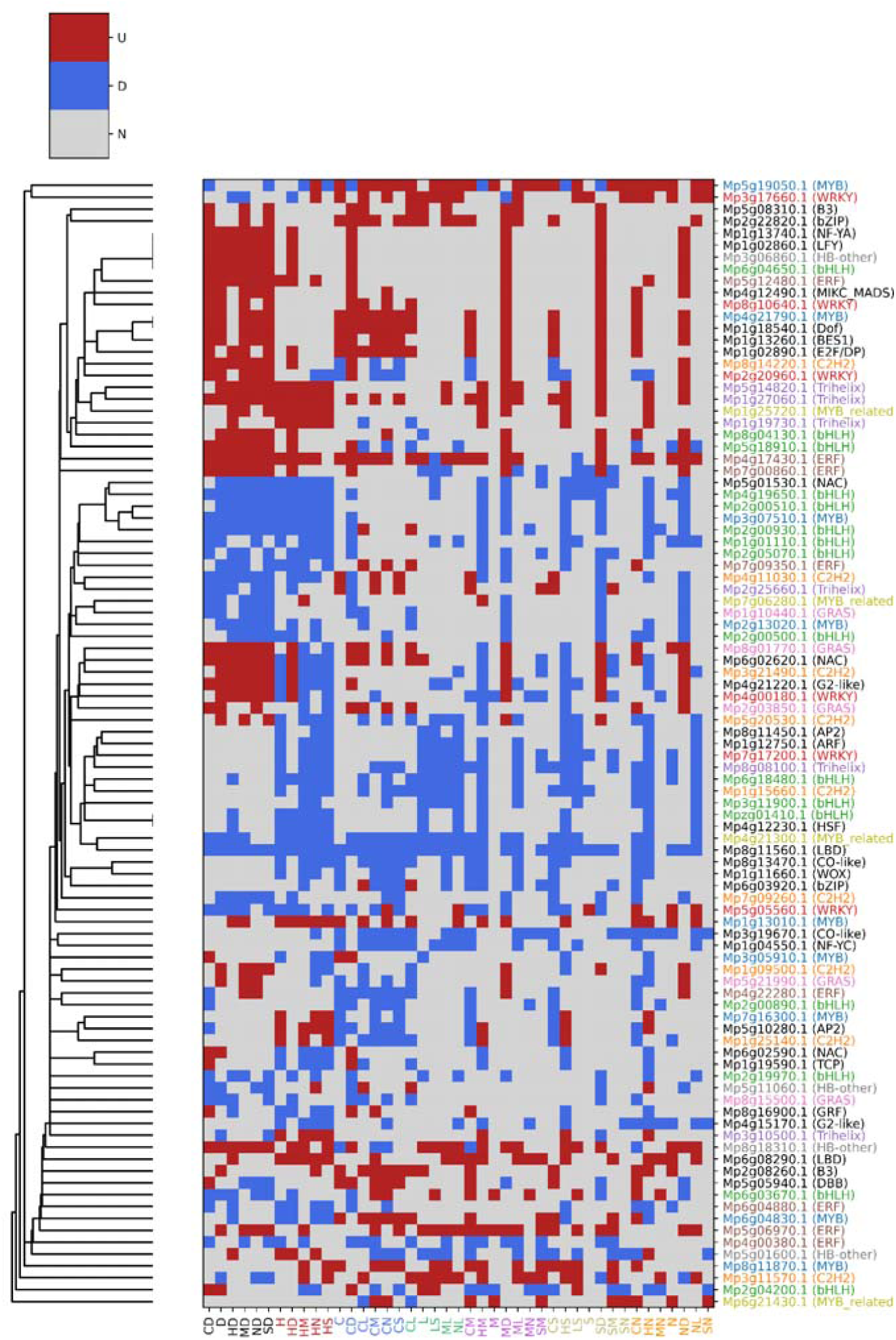
Expression profile of transcription factors involved in GRN reconstruction. Red, blue, and gray cells represent up-, down-regulated and unchanged expression respectively. The similarity of transcription factors was calculated based on the Jaccard index. Transcription factors with more than 3 members in a family are assigned a unique color.

**Figure S11.**
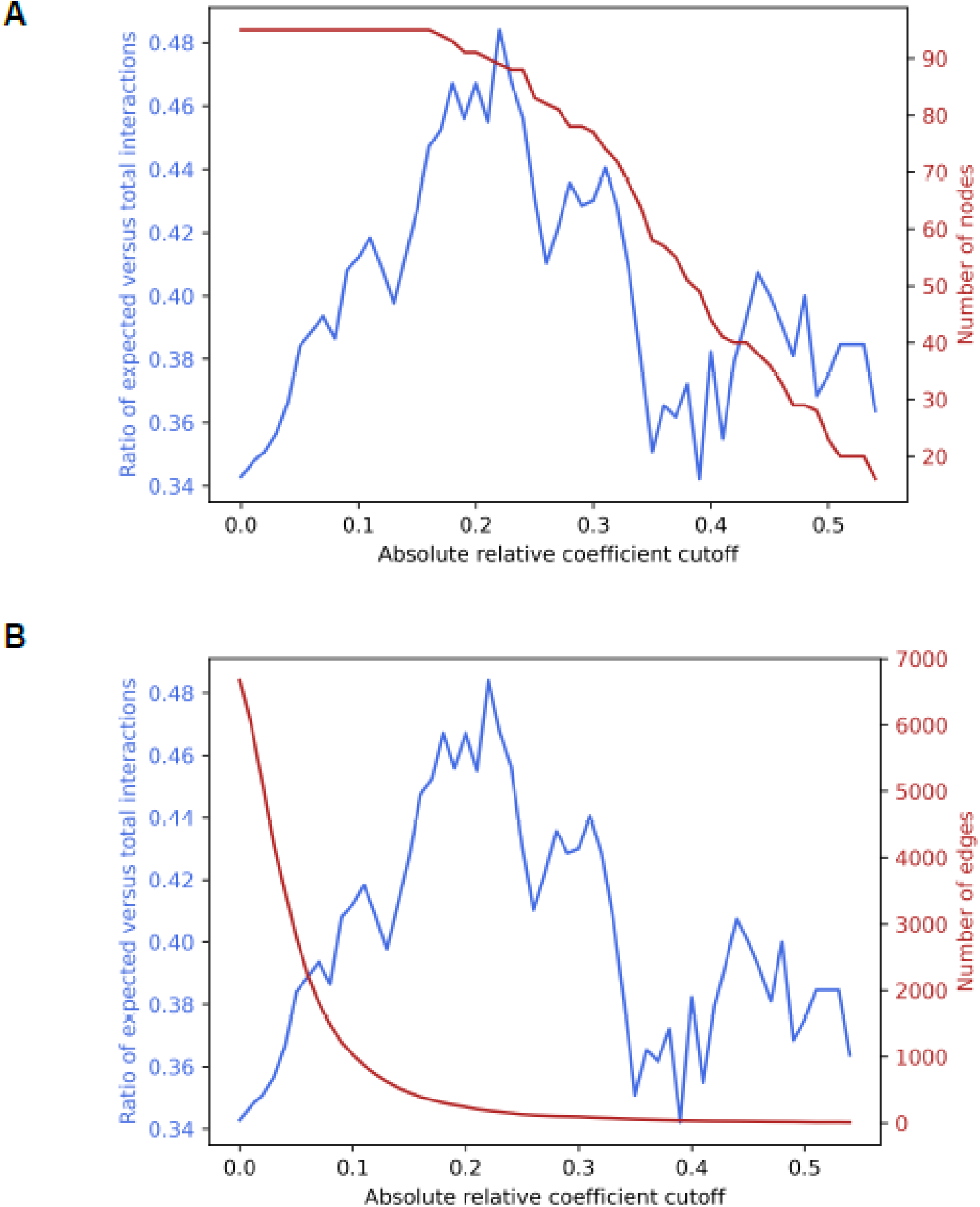
The ratio of expected interactions across absolute coefficient cut-offs. A) Number of nodes retained. B) Number of edges retained.

**Figure S12.**
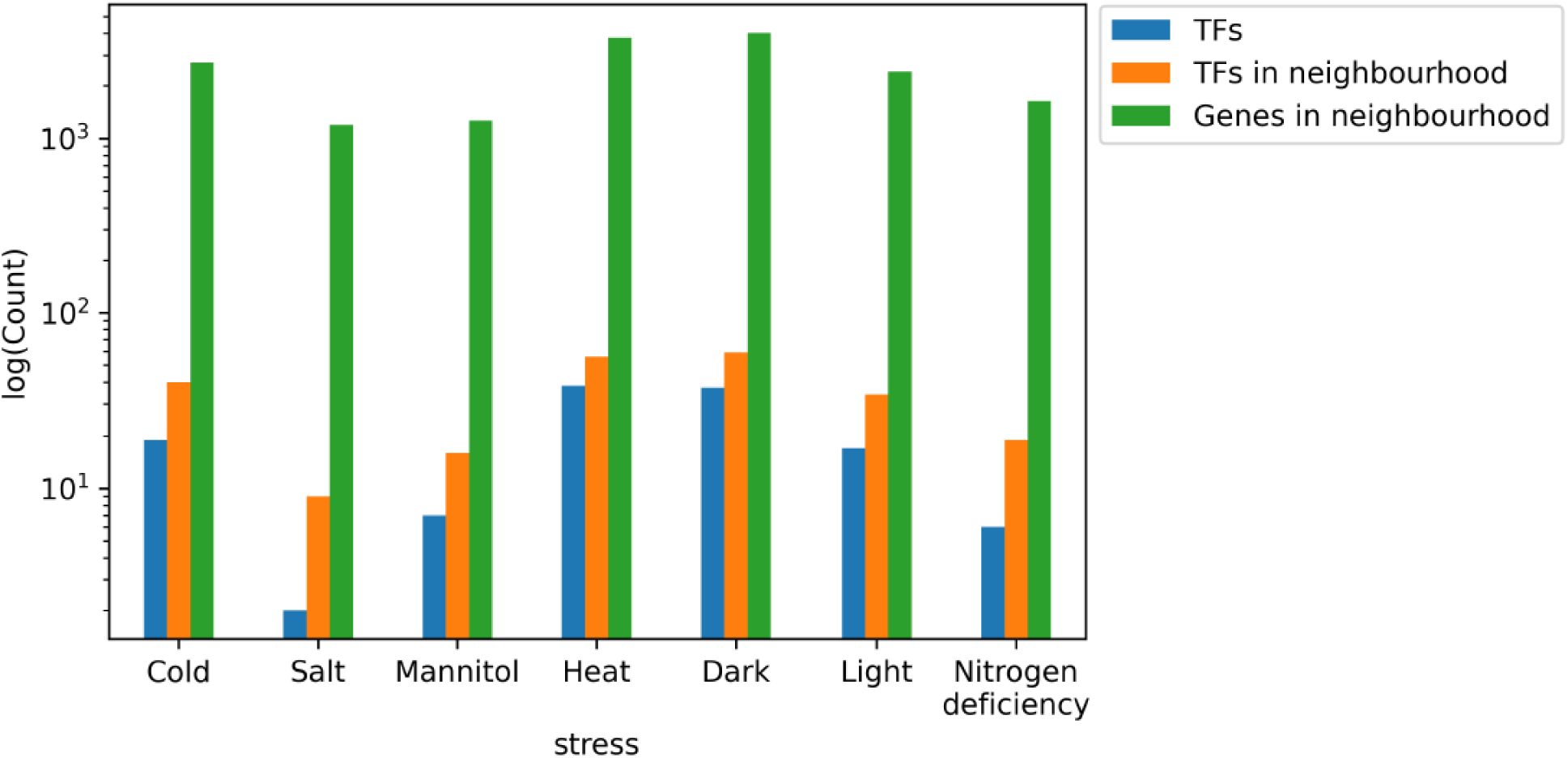
The magnitude of response in GRN for each stress. Blue, orange, and green bars reflect the number of stress-specific TFs, the number of TFs in the first neighborhood of the former, and genes associated with the first neighborhood respectively.

**Figure S13.**
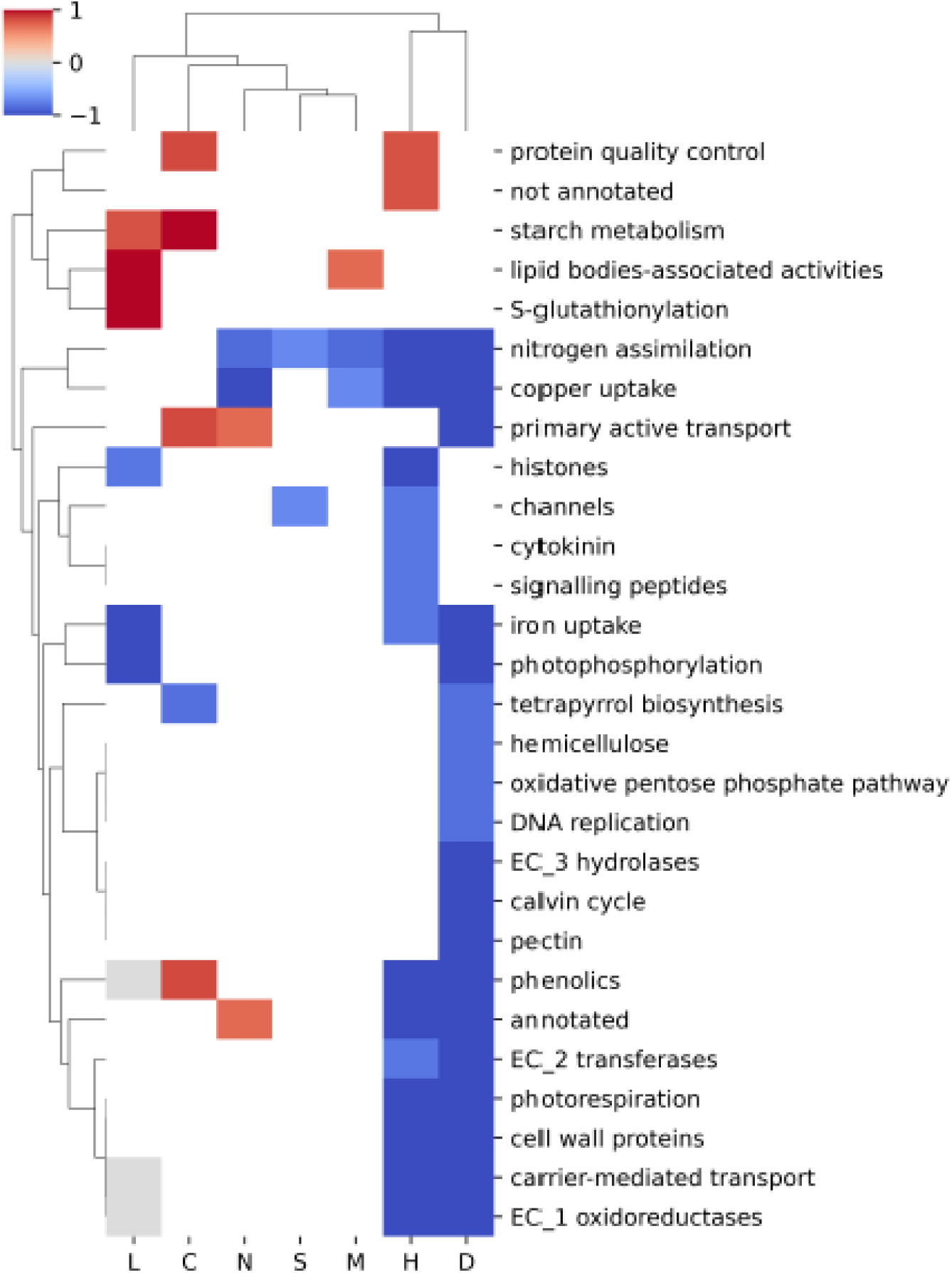
Robustly responding second-level mapman bins across the 7 abiotic stresses. Red, blue, and gray cells correspond to specifically up-, down-regulated, and no change respectively. Cells that do not contain enrichment for stress are in white.

**Figure S14.**
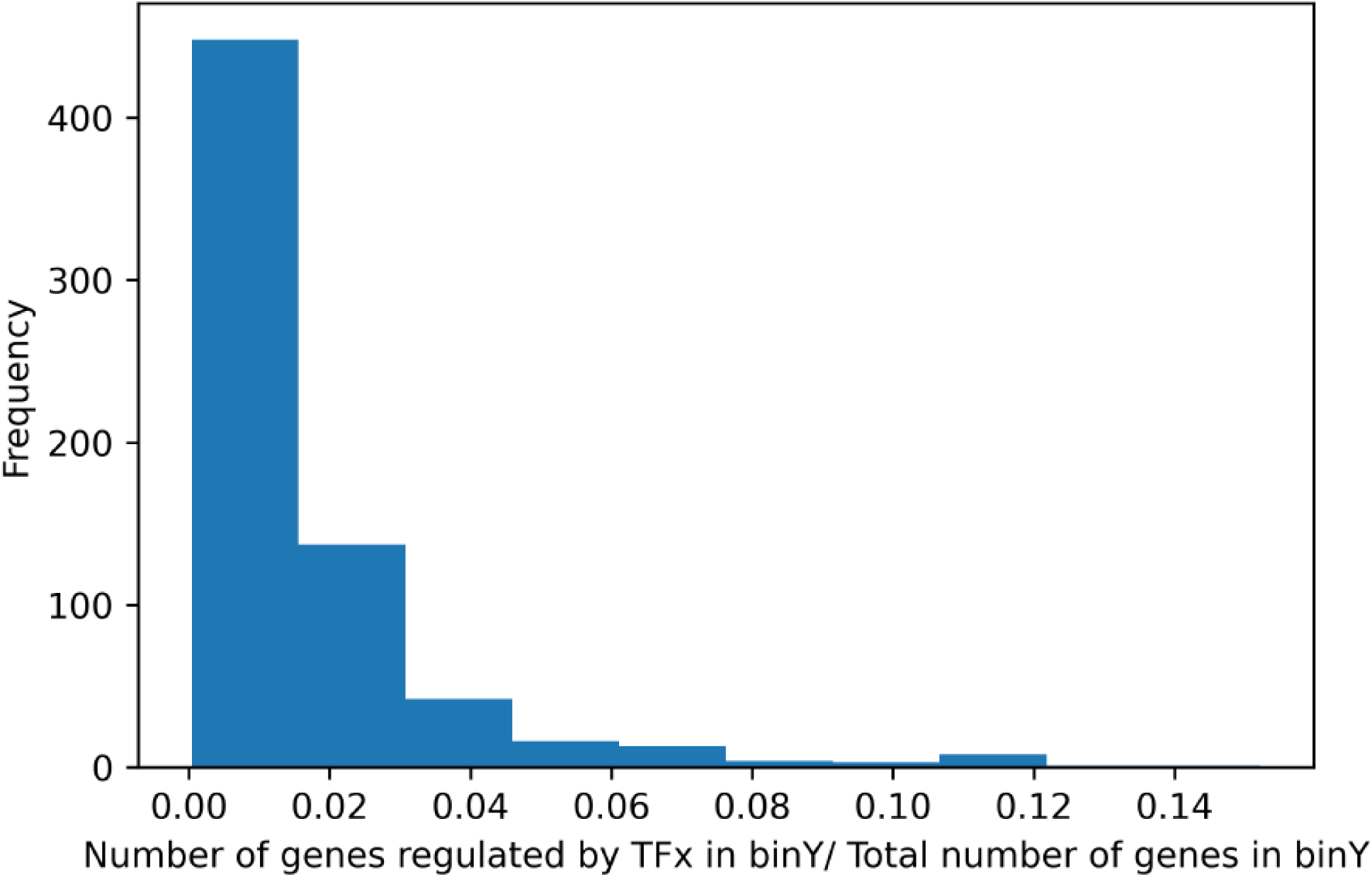
Frequency of ratio of robustly responding MapMan bins regulated by robustly responding transcription factors.

**Figure S15.**
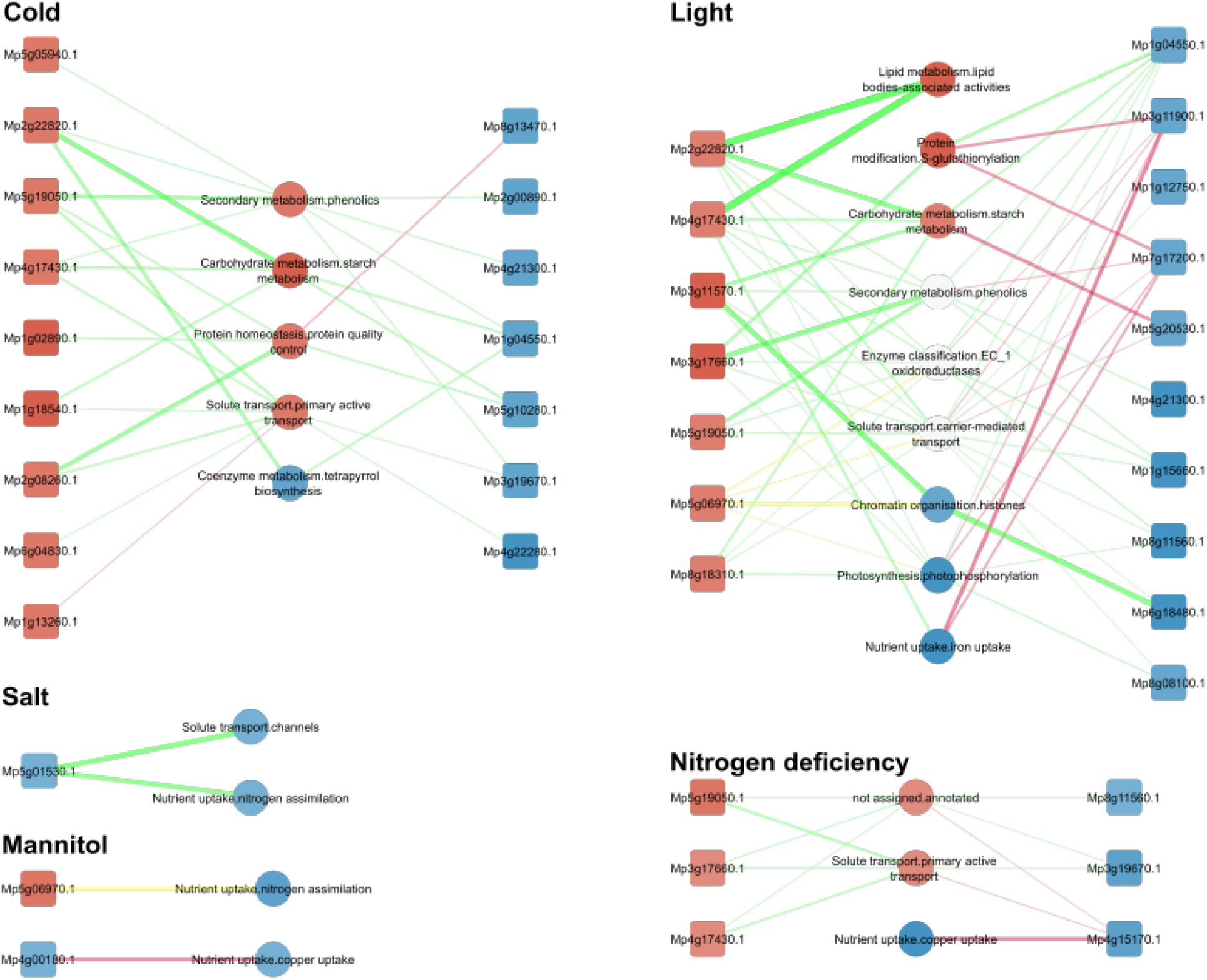
Representation of enriched second-level mapman bins under the regulation of specifically expressed transcription factors in A) Cold, B) Light, C) Salt, D) Mannitol and E) Nitrogen deficiency stresses. Node colors red and blue reflect the degree to which the nodes are specifically up or downregulated in the stress respectively. The edge width represents the ratio of genes under regulation as a fraction of the total number of genes found in that particular MapMan bin. The color of edges reflects the general trend of regulation of the TF, with green and red for positive and negative regulation respectively.

**Figure S16.**
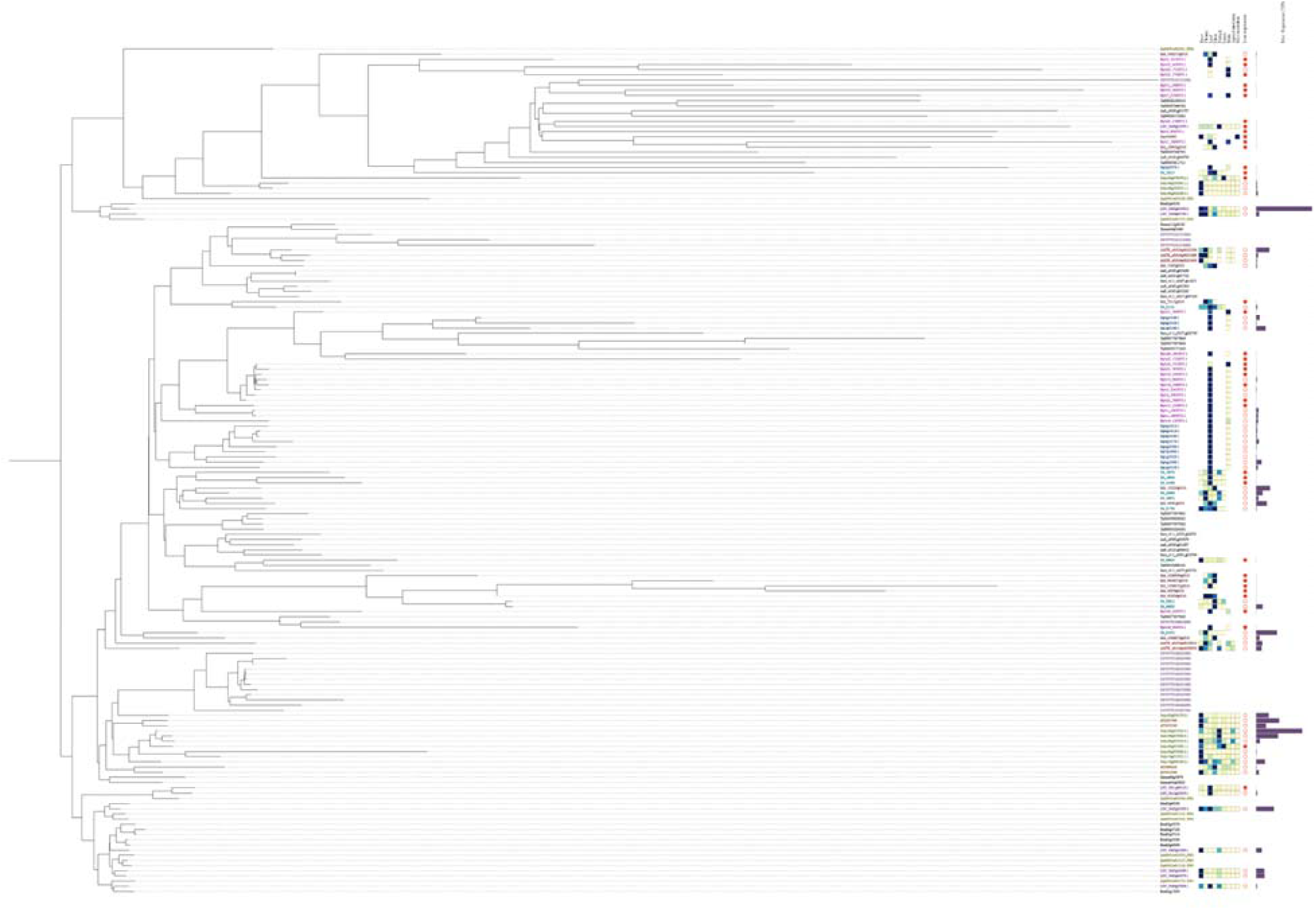
**Phylogenetic Tree of PAL OG_02_0000119 (Land Plants).**

## Supplementary Tables

**Supplementary Table 1.** Summary of experiment label to experiment conditions and mapping statistics from Kallisto.

**Supplementary Table 2.** TPM normalized gene expression matrix for *Marchantia* experiments

**Supplementary Table 3.** Raw gene expression matrix for *Marchantia* experiments

**Supplementary Table 4.** PCC values within *Marchantia* experiment replicates

**Supplementary Table 5.** Summary of DESeq2 output of *Marchantia* stress experiments

**Supplementary Table 6.** List of genes that are identified to be differentially expressed in both controls with corresponding MapMan bins and annotation. ’UP’ and ’DOWN’ in L2FC_D2 and L2FC_H2 columns indicate significant up and downregulation when compared against control D2 and H2 respectively.

**Supplementary Table 7.** Evidence of function in *Arabidopsis* orthologs of *Marchantia* transcription factors based on literature and gene expression changes.

**Supplementary Table 8.** Incidents of evidence in *Arabidopsis* (yellow) for each group of robustly responding *Marchantia* transcription factor (green)

**Supplementary Table 9.** Cold-Nitrogen deficiency specific genes as obtained from conekt.plant.tools

