## Supplemental Figure S1-17 for "Cross-stress gene expression atlas of *Marchantia polymorpha* reveals the hierarchy and regulatory principles of abiotic stress responses"

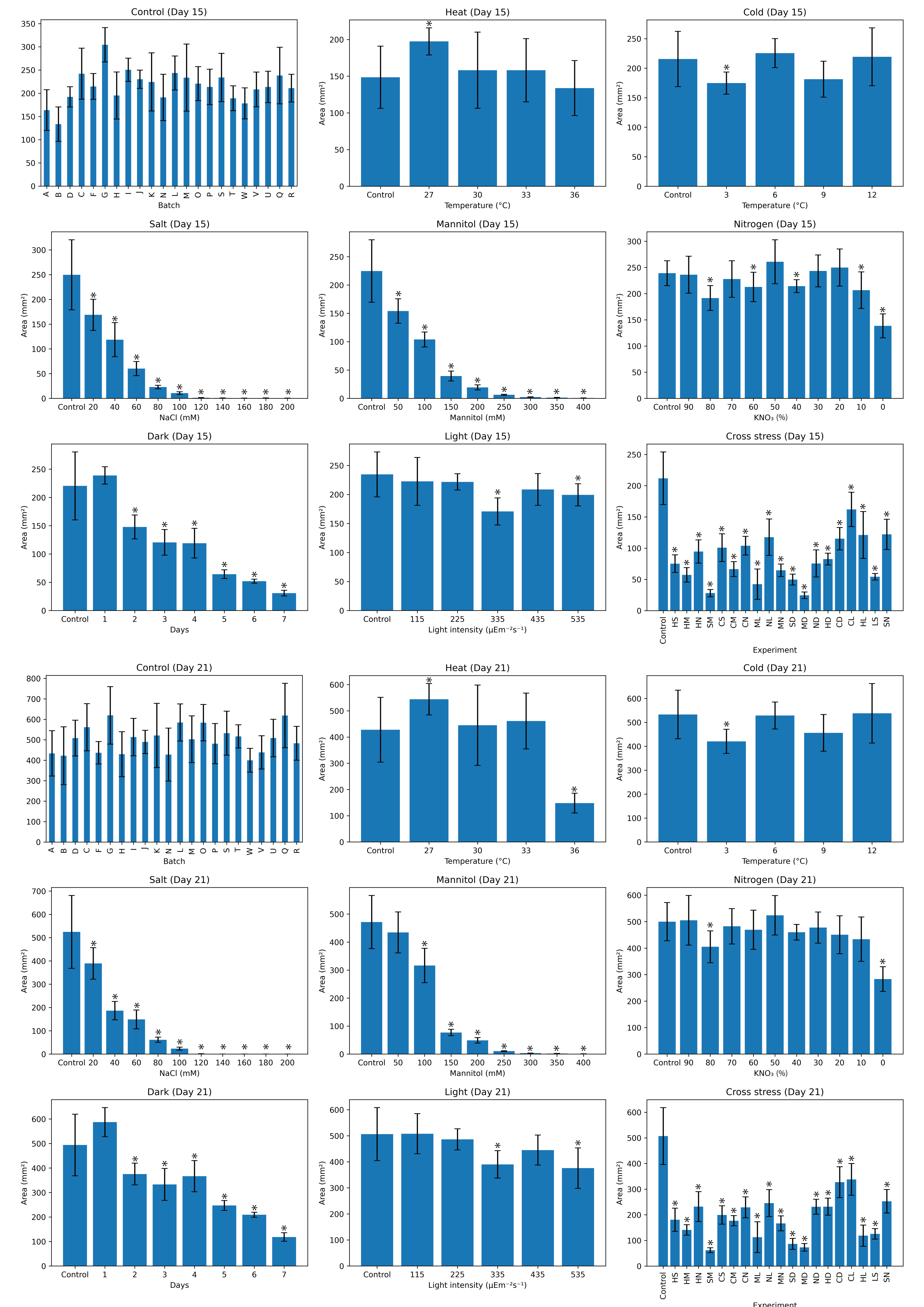

Control (Day 15)
0 50 100 150 200 250 300 350
A B D C F G H I J K N L M O P S T W V U Q R
Batch
Heat (Day 15)
Area (mm²)
0 50 100 150 200
Control 27 30 33 36
Temperature (°C)
Cold (Day 15)
Area (mm²)
0 50 100 150 200 250
Control 3 6 9 12
Temperature (°C)
Salt (Day 15)
Area (mm²)
0 50 100 150 200 250 300
Control 20 40 60 80 100 120 140 160 180 200
NaCl (mM)
Mannitol (Day 15)
Area (mm²)
0 50 100 150 200 250
Control 50 100 150 200 250 300 350 400
Mannitol (mM)
Nitrogen (Day 15)
Area (mm²)
0 50 100 150 200 250 300
Control 90 80 70 60 50 40 30 20 10 0
KNO₃ (%)
Dark (Day 15)
Area (mm²)
0 50 100 150 200 250
Control 1 2 3 4 5 6 7
Days
Light (Day 15)
Area (mm²)
0 50 100 150 200 250
Control 115 225 335 435 535
Light intensity (µEm⁻²s⁻¹)
Cross stress (Day 15)
Area (mm²)
0 50 100 150 200 250
Control HS HM HN SM CS CM CN ML NL MN SD MD ND HD CD CL HL LS SN
Experiment
Control (Day 21)
0 100 200 300 400 500 600 700 800
A B D C F G H I J K N L M O P S T W V U Q R
Batch
Heat (Day 21)
Area (mm²)
0 100 200 300 400 500 600
Control 27 30 33 36
Temperature (°C)
Cold (Day 21)
Area (mm²)
0 100 200 300 400 500 600
Control 3 6 9 12
Temperature (°C)
Salt (Day 21)
Area (mm²)
0 100 200 300 400 500 600 700
Control 20 40 60 80 100 120 140 160 180 200
NaCl (mM)
Mannitol (Day 21)
Area (mm²)
0 100 200 300 400 500
Control 50 100 150 200 250 300 350 400
Mannitol (mM)
Nitrogen (Day 21)
Area (mm²)
0 100 200 300 400 500 600
Control 90 80 70 60 50 40 30 20 10 0
KNO₃ (%)
Dark (Day 21)
Area (mm²)
0 100 200 300 400 500 600
Control 1 2 3 4 5 6 7
Days
Light (Day 21)
Area (mm²)
0 100 200 300 400 500 600
Control 115 225 335 435 535
Light intensity (µEm⁻²s⁻¹)
Cross stress (Day 21)
Area (mm²)
0 100 200 300 400 500 600
Control HS HM HN SM CS CM CN ML NL MN SD MD ND HD CD CL HL LS SN
Experiment

**Figure S1. Area of plants at 15 days (day of harvest) and day 21 (6 days post harvest) for the single and combined stresses.** Each bar represents measurements from at least 4 plants. Error bars are represented by standard deviation and significance was determined by Student’s two-tailed, t-test, $p < 0.05$.

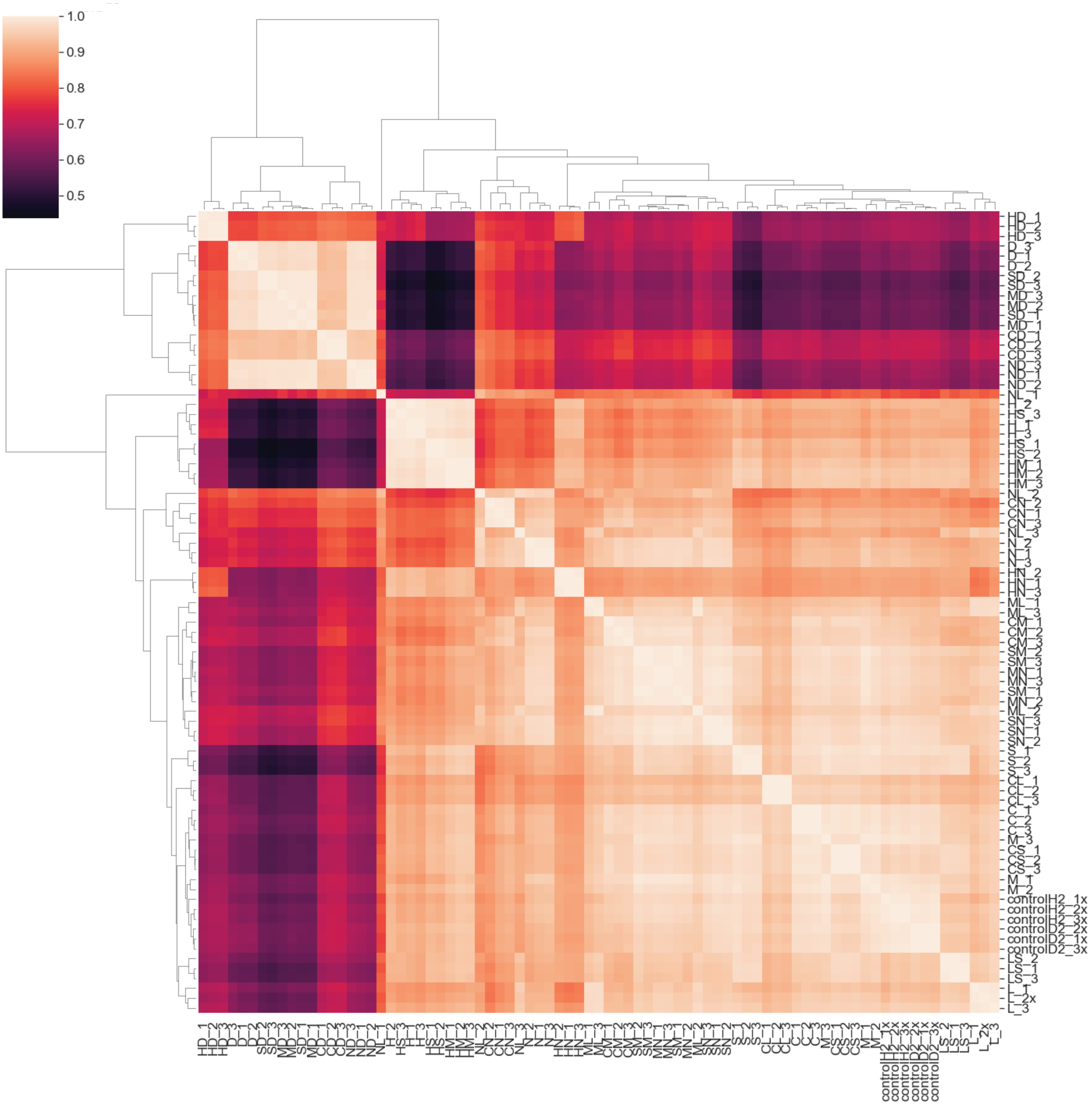


**Figure S2. Hierarchical clustering of the Marchantia stress experiments.** Transcript per million (TPM) gene expression values were scaled with standard scaler and clustered with seaborn.clustermap() function.

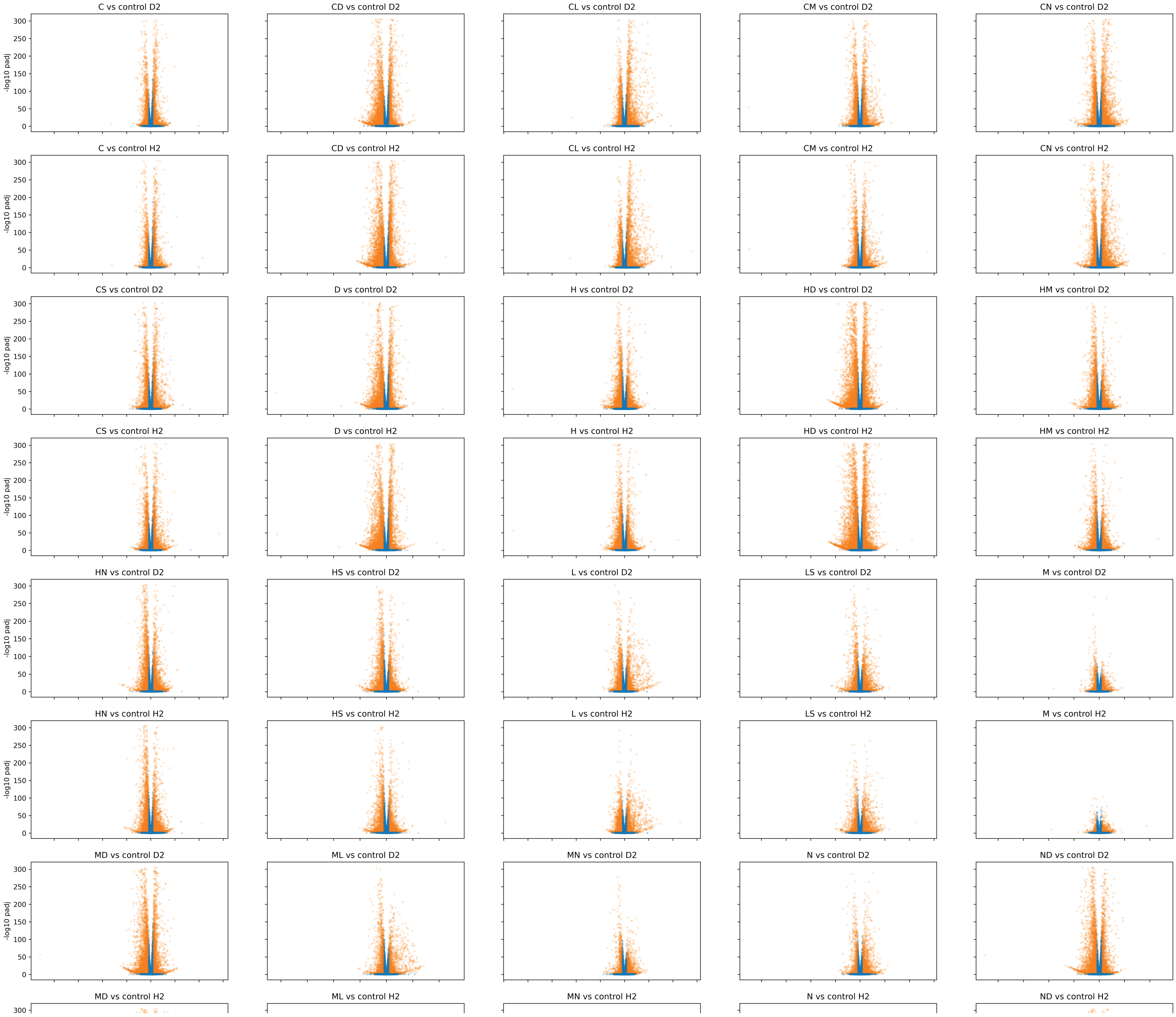


**Figure S3. Volcano plots of stress experiments.** The x-axis represents log2-fold change, the y-axis indicates the vv-log10 of adjusted p-value. Each point represents a gene that has an absolute log2-fold change more than 1 and is significantly (adjusted p-value<0.05, orange color) or not significantly (p-value>0.05, blue point) differentially expressed.

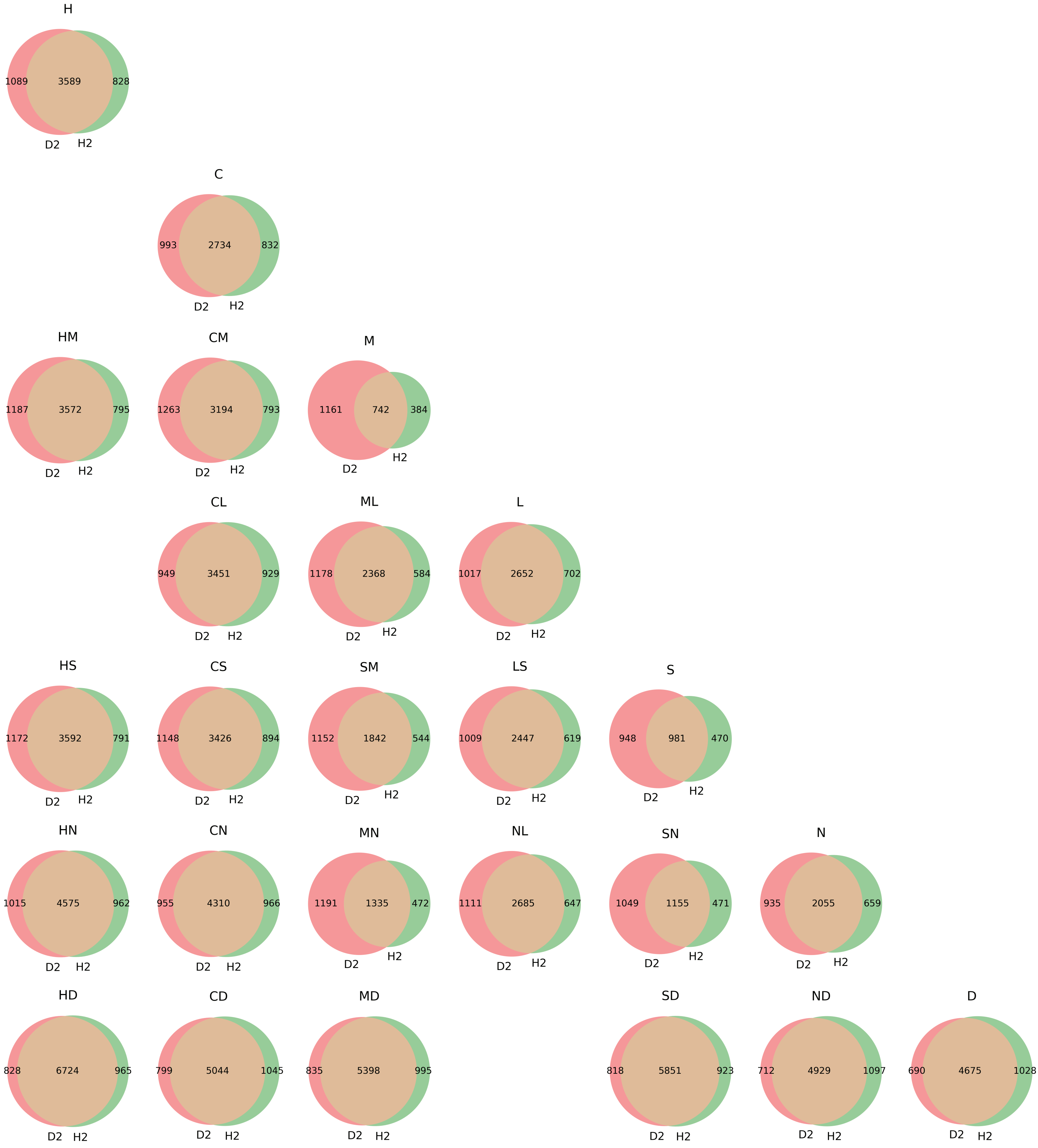


**Figure S4. Comparison of found differentially expressed genes for the single and combined stressed.** For each stress combination, we estimated significant DEGs (adjusted p-value<0.05) by using control D2 (red circle) and control H2 (green circle). The orange areas indicate the overlap between the found DEGs of the two controls. The overlapping DEGs were used for further study. The following abbreviations are used for the description of stresses: cold (C), dark (D), heat (H), light (L), mannitol (M), nitrogen deficiency (N) and salt (S). Combination of stresses is indicated by two letters, where e.g., HD indicates heat+darkness treatment.

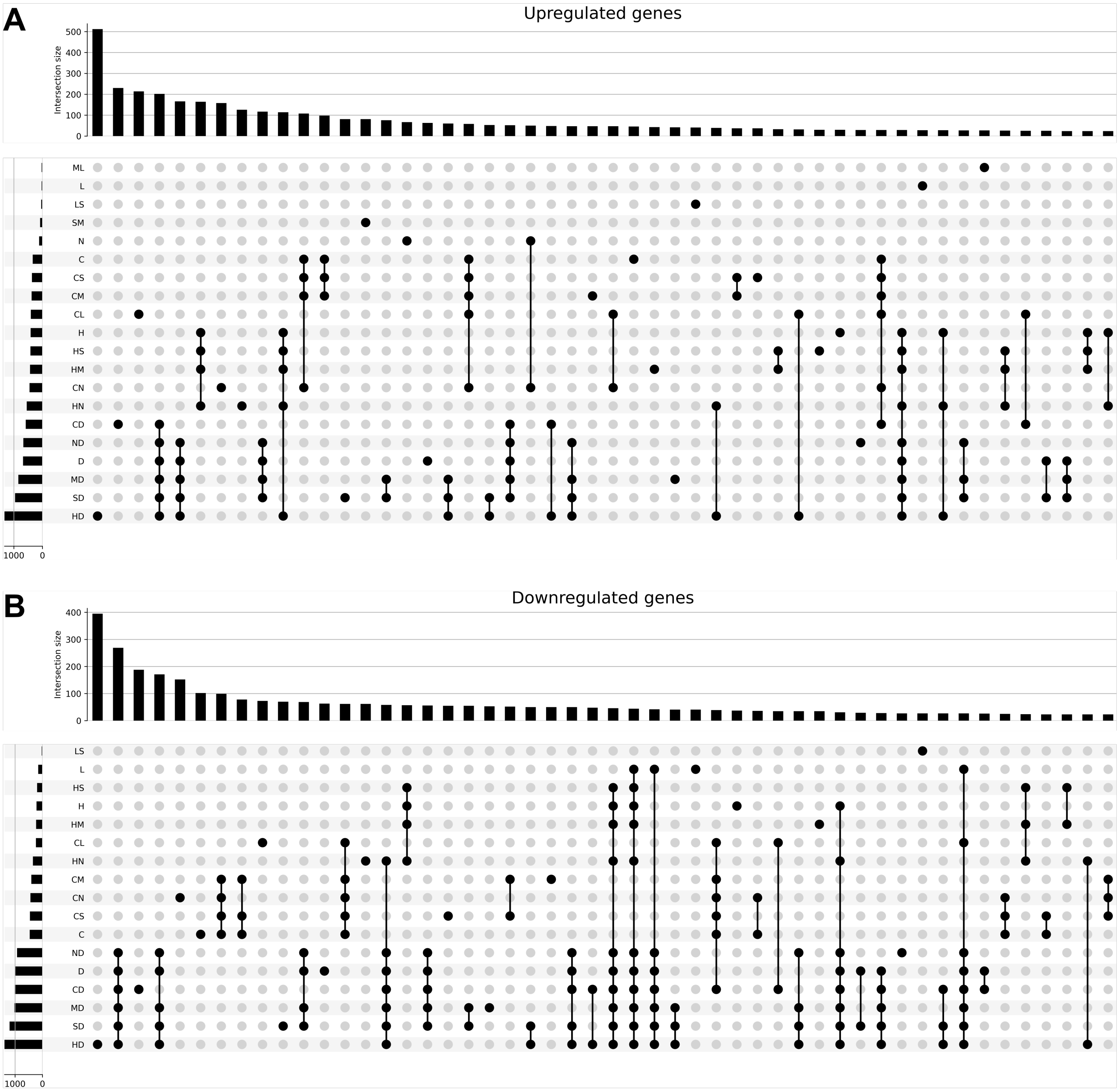


**Figure S5. Upset plot showing the top 50 intersections of differentially expressed genes.** A) Upregulated genes B) Downregulated genes. The package is available from https://upsetplot.readthedocs.io/en/stable/.

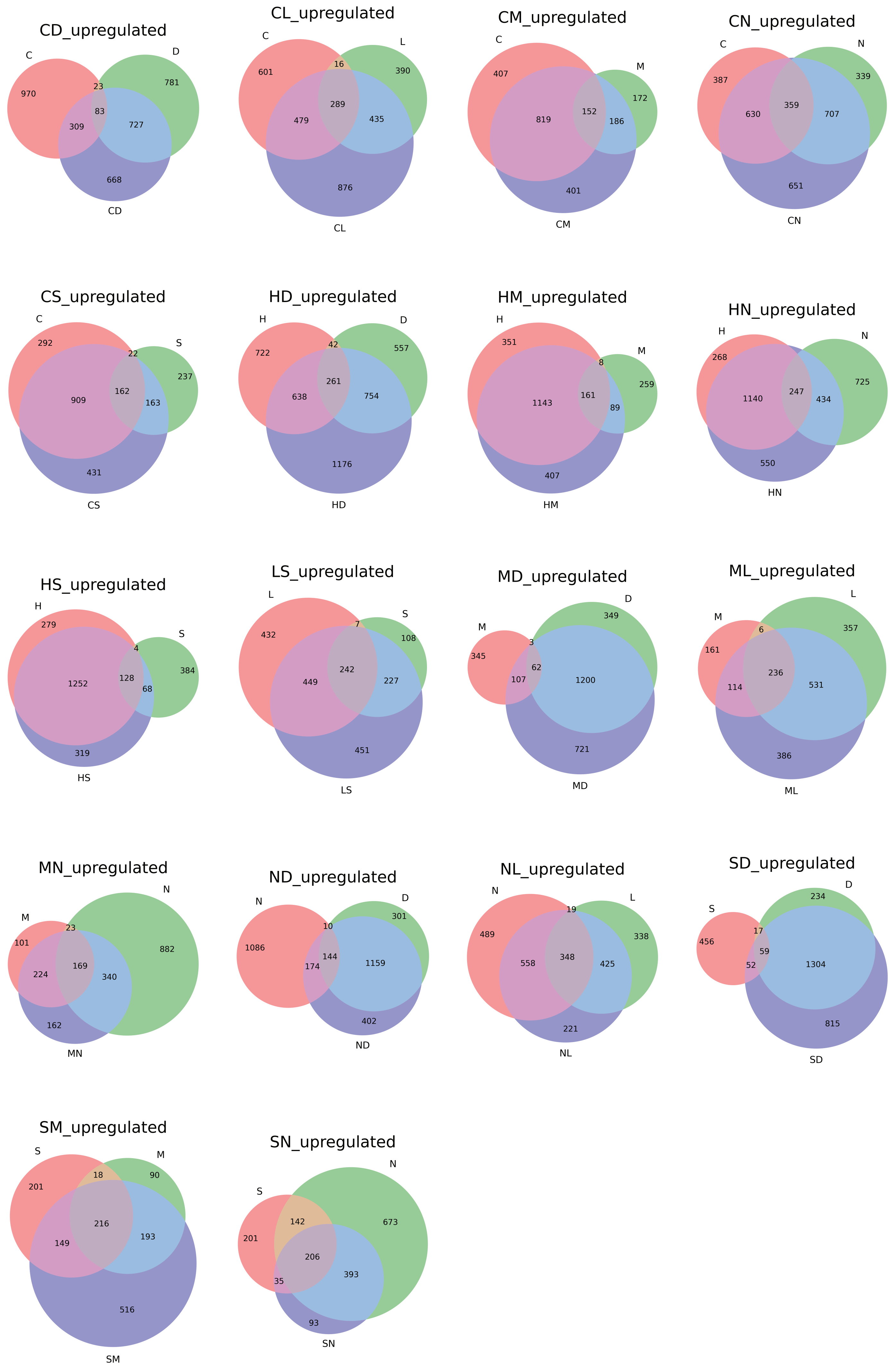
CD_upregulated
C
D
970
23
781
83
309
727
668
CD
CL_upregulated
C
L
16
601
390
289
479
435
876
CL
CM_upregulated
C
407
M
172
819
152
186
401
CM
CN_upregulated
C
N
387
339
359
630
707
651
CN
CS_upregulated
C
S
292
22
237
909
162
163
431
CS
HD_upregulated
H
D
42
722
557
261
638
754
1176
HD
HM_upregulated
H
351
M
8
259
161
1143
89
407
HM
HN_upregulated
H
N
268
725
247
1140
434
550
HN
HS_upregulated
H
279
S
4
384
128
1252
68
319
HS
LS_upregulated
L
S
7
432
108
242
449
227
451
LS
MD_upregulated
D
349
M
3
345
62
107
1200
721
MD
ML_upregulated
L
M
357
6
161
236
114
531
386
ML
MN_upregulated
N
M
23
101
882
169
224
340
162
MN
ND_upregulated
N
D
10
301
1086
144
174
1159
402
ND
NL_upregulated
N
L
19
489
338
348
558
425
221
NL
SD_upregulated
D
234
S
17
456
59
52
1304
815
SD
SM_upregulated
S
M
18
90
201
216
149
193
516
SM
SN_upregulated
N
S
142
673
201
206
393
35
93
SN

**Figure S6. Venn diagrams of upregulated genes in single and combined stresses.** The abbreviates are cold (C), dark (D), heat (H), light (L), mannitol (M), nitrogen deficiency (N) and salt (S). Combination of stresses is indicated by two letters and purple circles. The first and second stress in a pair is colored red and green (e.g., cold is red in CD), while the combined stress is colored purple. The sizes of the circles and intersections and the numbers within indicate the number of significant DEGs.

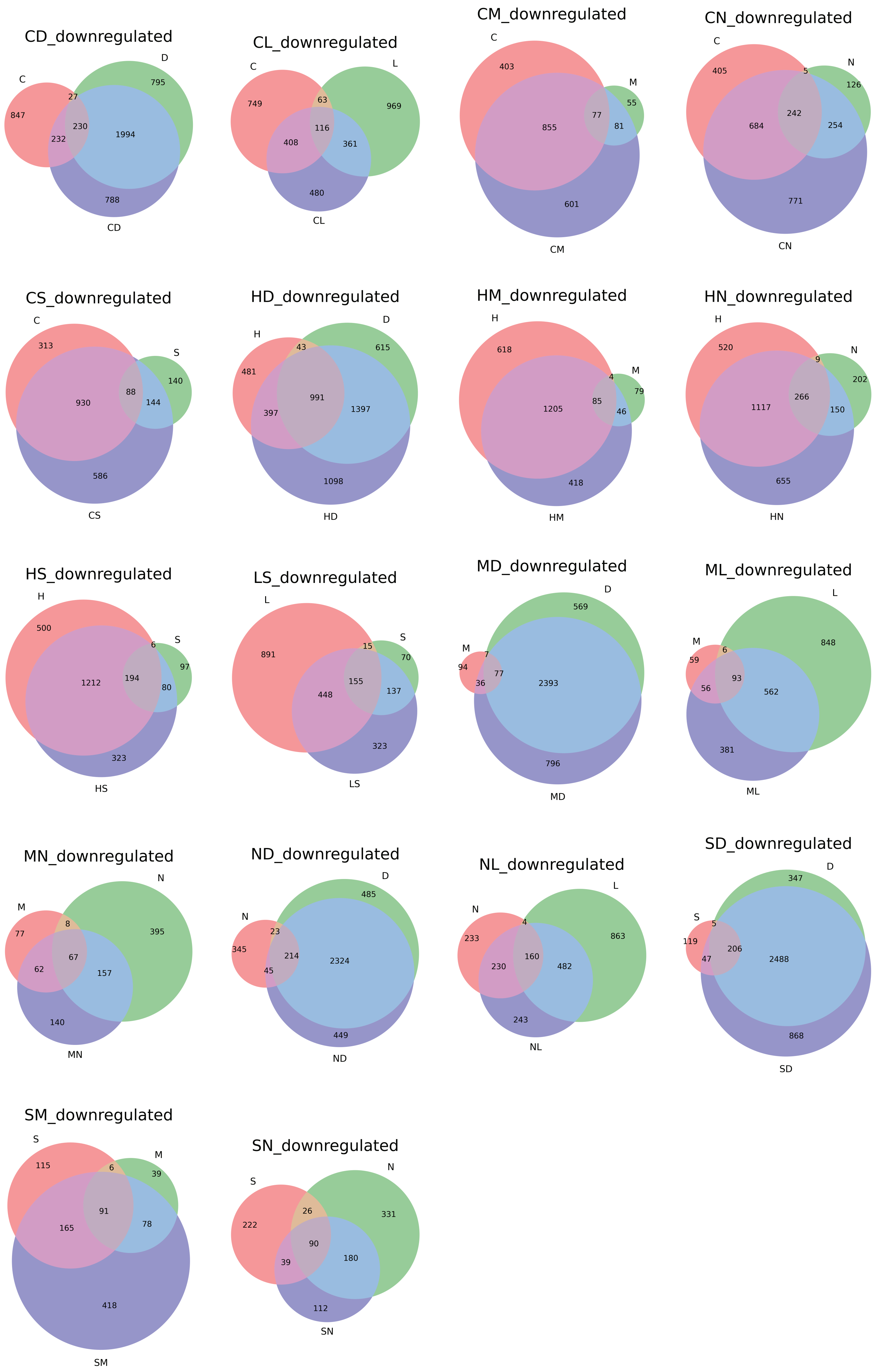
CD_downregulated
C
D
847
27
795
232
230
1994
788
CD
CL_downregulated
C
L
749
63
969
408
116
361
480
CL
CM_downregulated
C
M
403
55
855
77
81
601
CM
CN_downregulated
C
N
405
5
126
684
242
254
771
CN
CS_downregulated
C
S
313
140
930
88
144
586
CS
HD_downregulated
H
D
43
615
481
991
397
1397
1098
HD
HM_downregulated
H
M
618
4
79
1205
85
46
418
HM
HN_downregulated
H
N
520
9
202
1117
266
150
655
HN
HS_downregulated
H
S
500
6
97
1212
194
80
323
HS
LS_downregulated
L
S
891
15
70
448
155
137
323
LS
MD_downregulated
D
M
569
7
94
77
36
2393
796
MD
ML_downregulated
L
M
59
6
848
56
93
562
381
ML
MN_downregulated
M
N
77
8
395
62
67
157
140
MN
ND_downregulated
N
D
485
23
345
214
2324
45
449
ND
NL_downregulated
N
L
4
863
233
160
230
482
243
NL
SD_downregulated
S
D
347
5
119
206
47
2488
868
SD
SM_downregulated
S
M
115
6
39
91
165
78
418
SM
SN_downregulated
S
N
26
222
331
90
39
180
112
SN

**Figure S7. Venn diagrams of downregulated genes in single and combined stresses.** The abbreviates are cold (C), dark (D), heat (H), light (L), mannitol (M), nitrogen deficiency (N) and salt (S). Combination of stresses is indicated by two letters and purple circles. The first and second stress in a pair is colored red and green (e.g., cold is red in CD), while the combined stress is colored purple. The sizes of the circles and intersections and the numbers within indicate the number of significant DEGs.

**A**

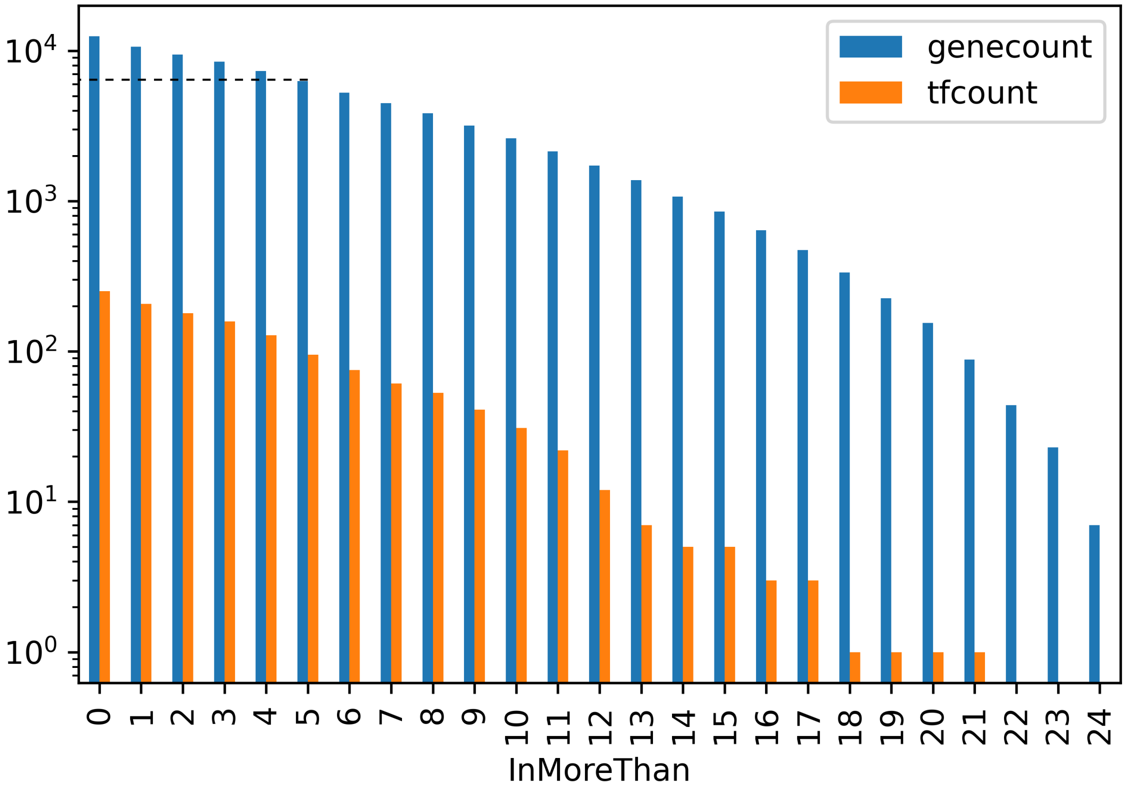


**B**

| Stress-type | Cold | Dark | Heat | Light | Mannitol | Nitrogen | Salt |
|---|---|---|---|---|---|---|---|
| Experiment availability | C<br>CS<br>CM<br>CN<br>CD<br>CL | D<br>SD<br>MD<br>ND<br>HD<br>CD | H<br>HS<br>HM<br>HN<br>HD | L<br>ML<br>NL<br>CL<br>LS | M<br>HM<br>CM<br>SM<br>ML<br>MN<br>MD | N<br>HN<br>CN<br>NL<br>MN<br>ND<br>SN | S<br>HS<br>CS<br>SM<br>SD<br>LS<br>SN |
| Total (with replicates) | 18 | 18 | 15 | 15 | 21 | 21 | 21 |

**Figure S8. Considerations for reconstruction of the stress-specific gene regulatory networks.** A) Number of differential genes in more than a certain number of experiments. B) List of experiments in each stress specific-network.

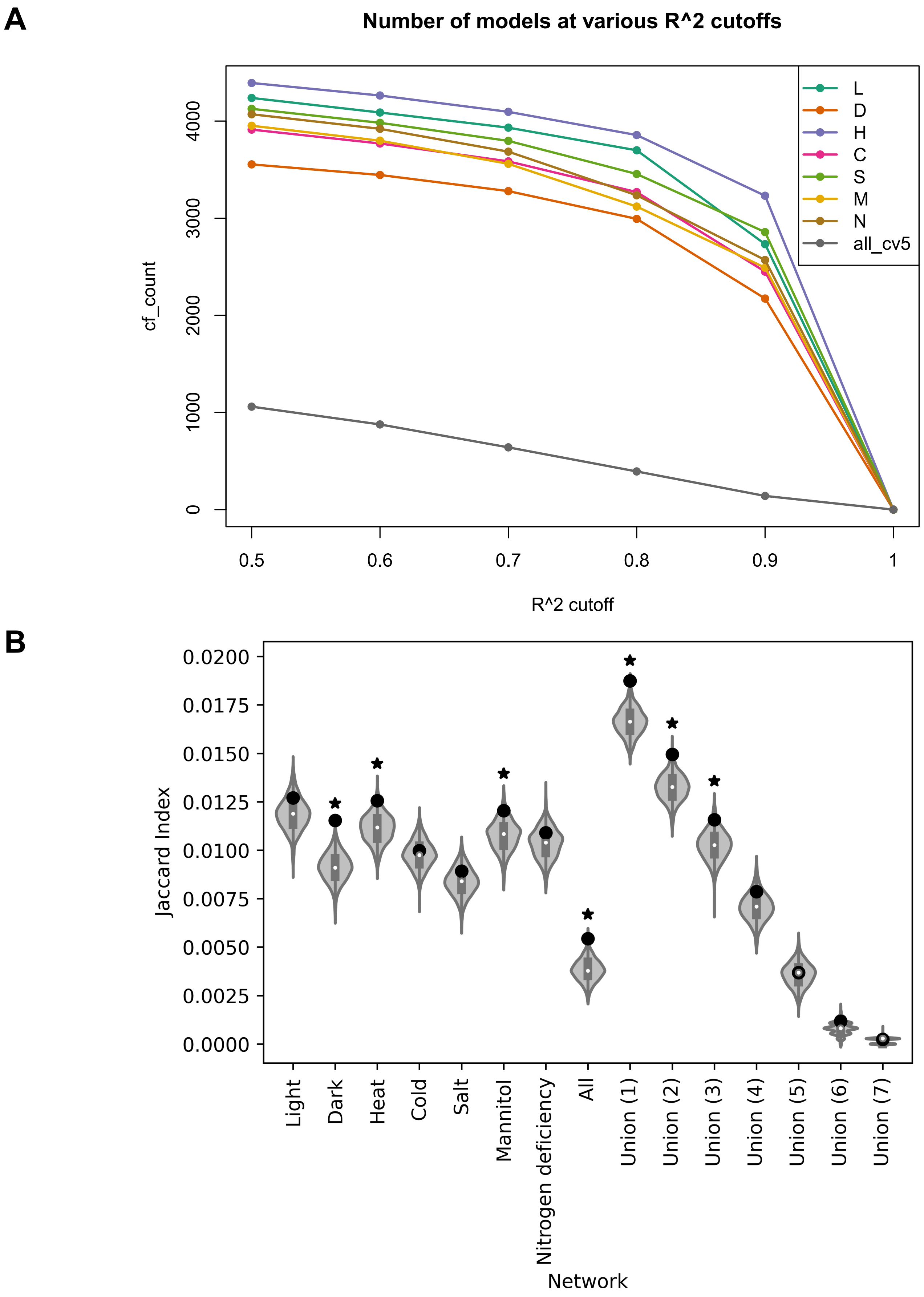


**Figure S9. Metrics used to construct the unified GRN.** A) Distribution of model counts across various $R^2$. B) Similarity of various gene regulatory networks against a shuffled AGRIS gene regulatory network. Point indicates observed similarity while asterisk indicates significance, hypergeometric test, $p < 0.05$.

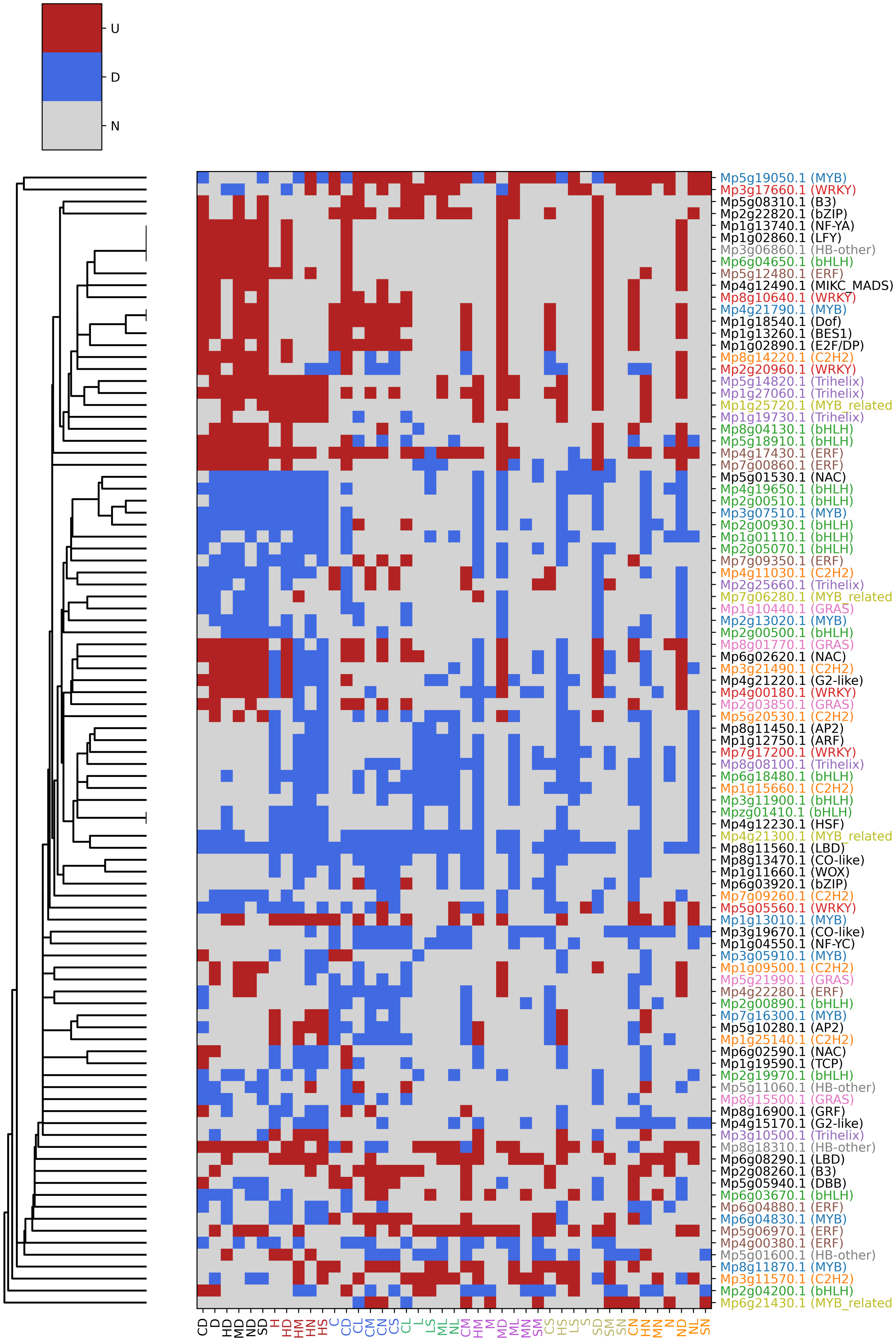


**Figure S10. Expression profile of transcription factors involved in GRN reconstruction.** Red, blue and gray cells represent up-, down-regulated and unchanged expression respectively. Similarity of transcription factors was calculated based on the jaccard index. Transcription factors with more than 3 members in a family are assigned a unique colour.

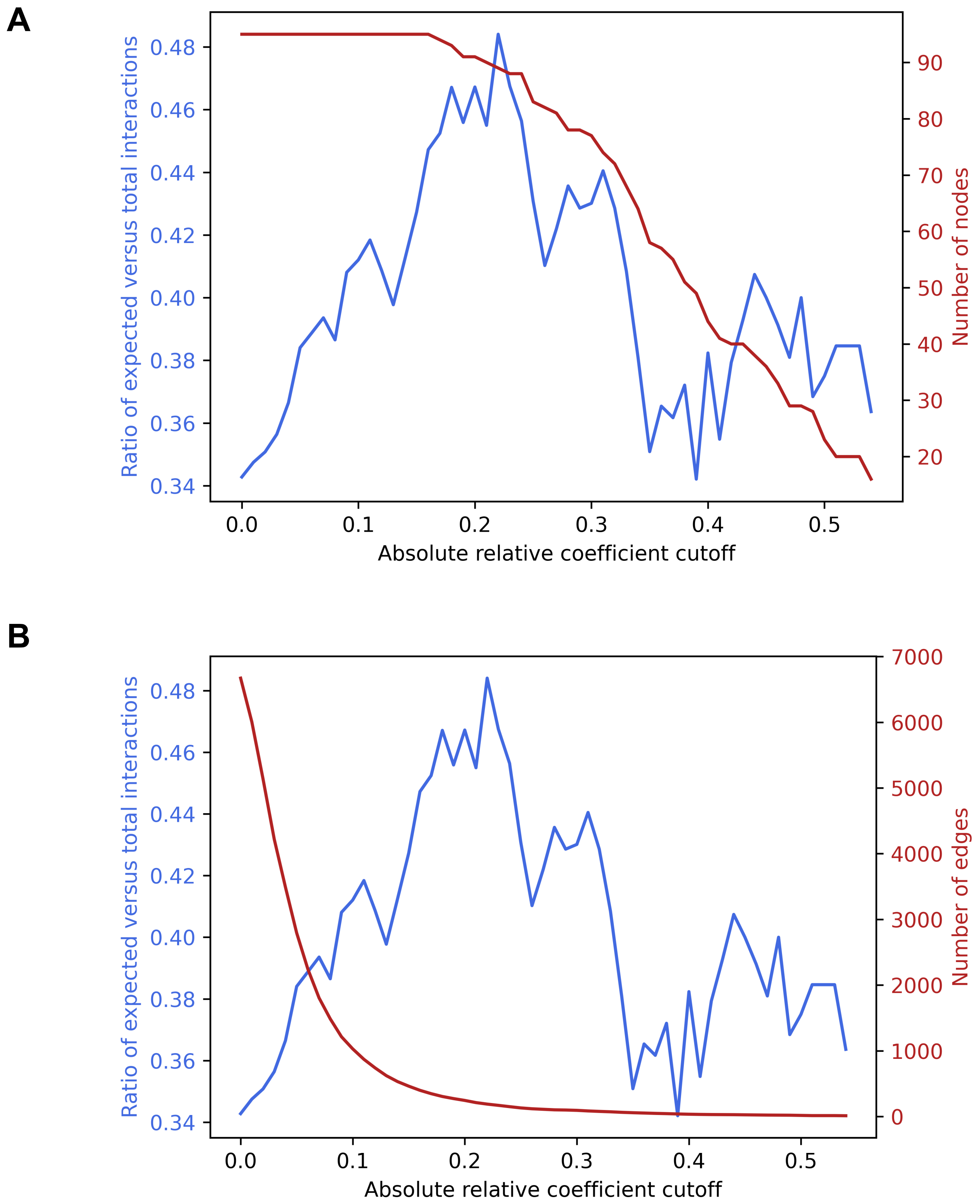


**Figure S11. Ratio of expected interactions across absolute coefficient cutoffs.** A) Number of nodes retained. B) Number of edges retained

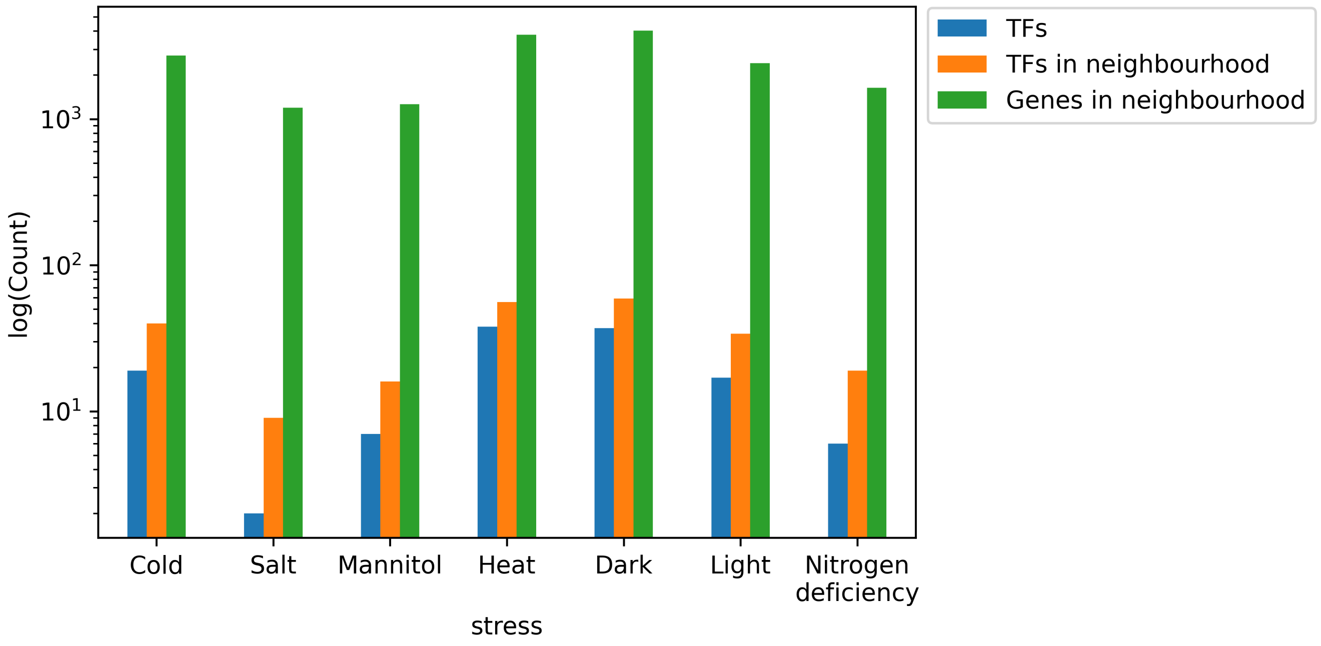


**Figure S12. Magnitude of response in GRN for each stress.** Blue, orange and green bars reflect the number of stress-specific TFs, number of TFs in the first neighborhood of the former and genes associated to the first neighborhood respectively.

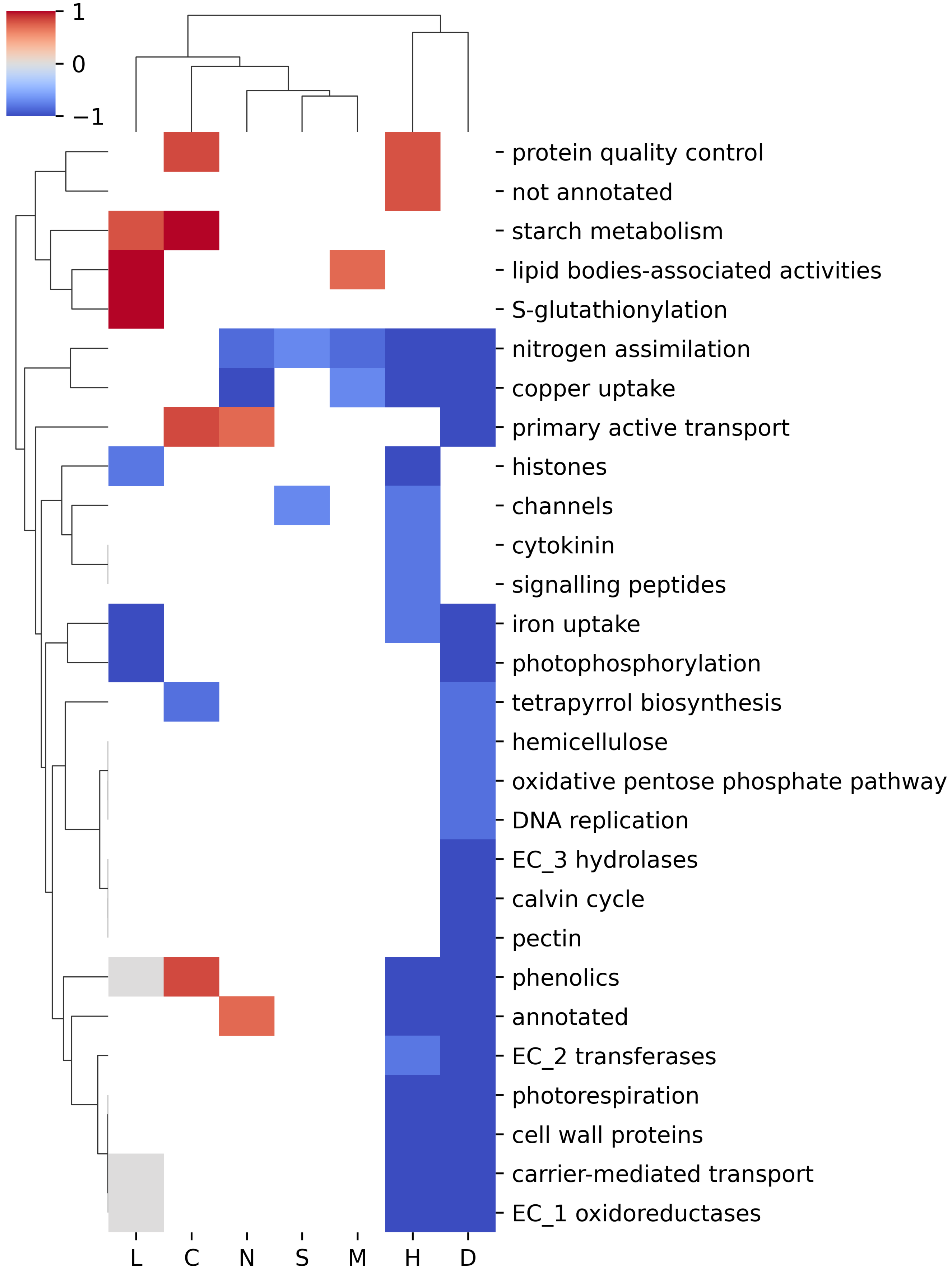


**Figure S13. Robustly responding second level mapman bins across the 7 abiotic stresses.** Red, blue and gray cells correspond to specifically up-, down-regulated and no change respectively. Cells that do not contain enrichment for stress are in white.

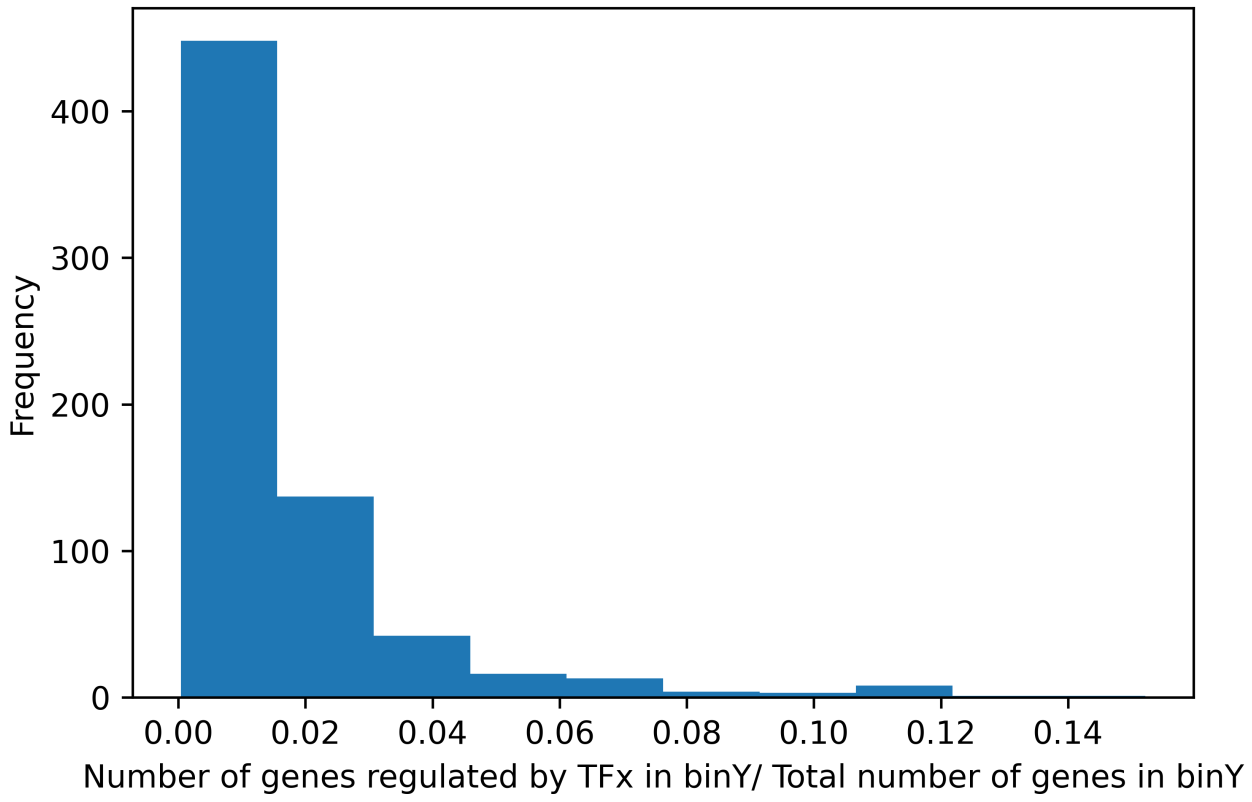


**Figure S14. Frequency of ratio of robustly responding mapman bins regulated by robustly responding transcription factors.**

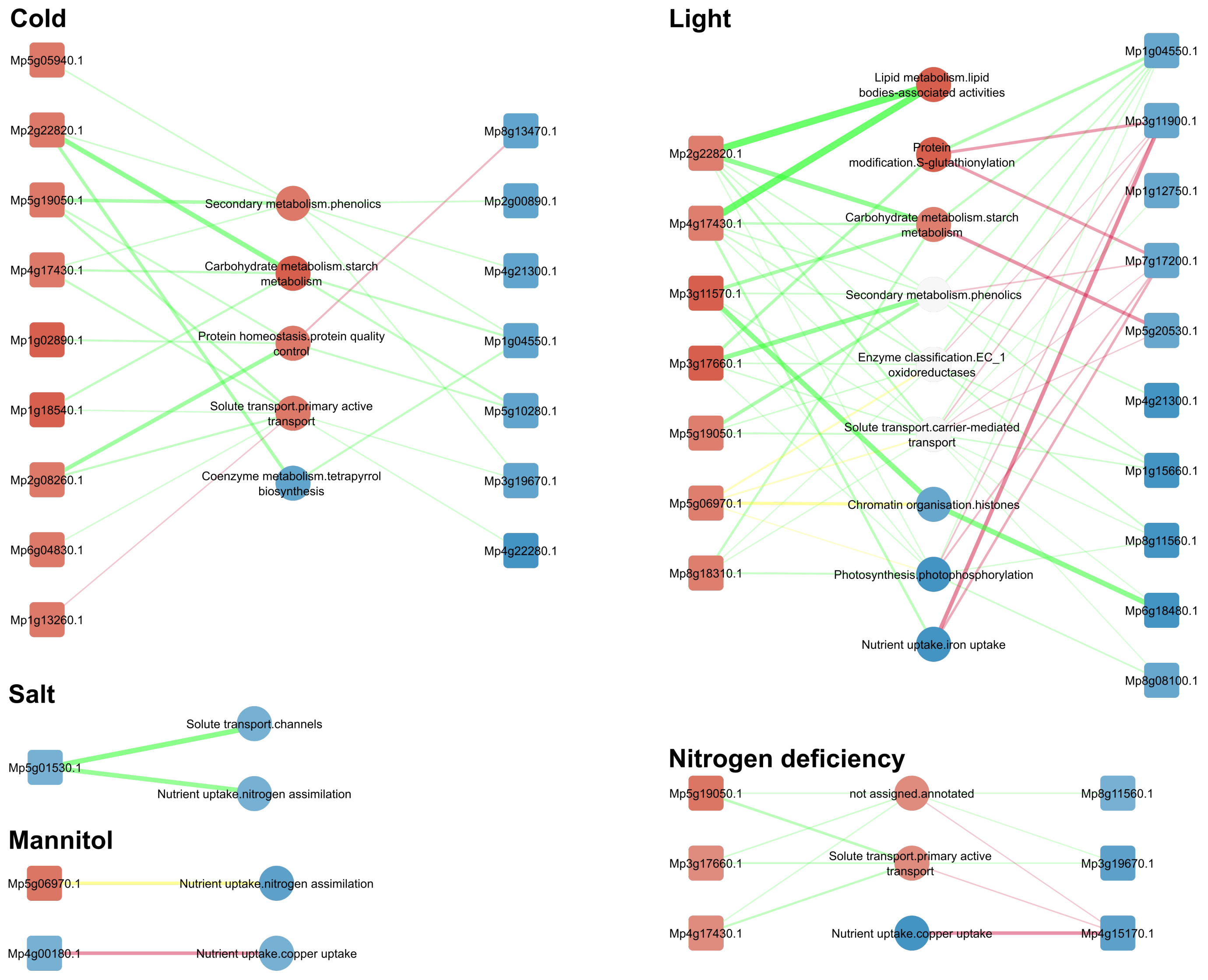


**Figure S15. Representation of enriched second level mapman bins under the regulation of specifically expressed transcription factors** in A) Cold, B) Light, C) Salt, D) Mannitol and E) Nitrogen deficiency stresses. Node colors red and blue reflect the degree where the nodes are specifically up or downregulated in the stress respectively. The edge width represents the ratio of genes under regulation as a fraction of the total number of genes found in that particular MapMan bin. Color of edges reflect the general trend of regulation of the TF, with green and red for positive and negative regulation respectively.

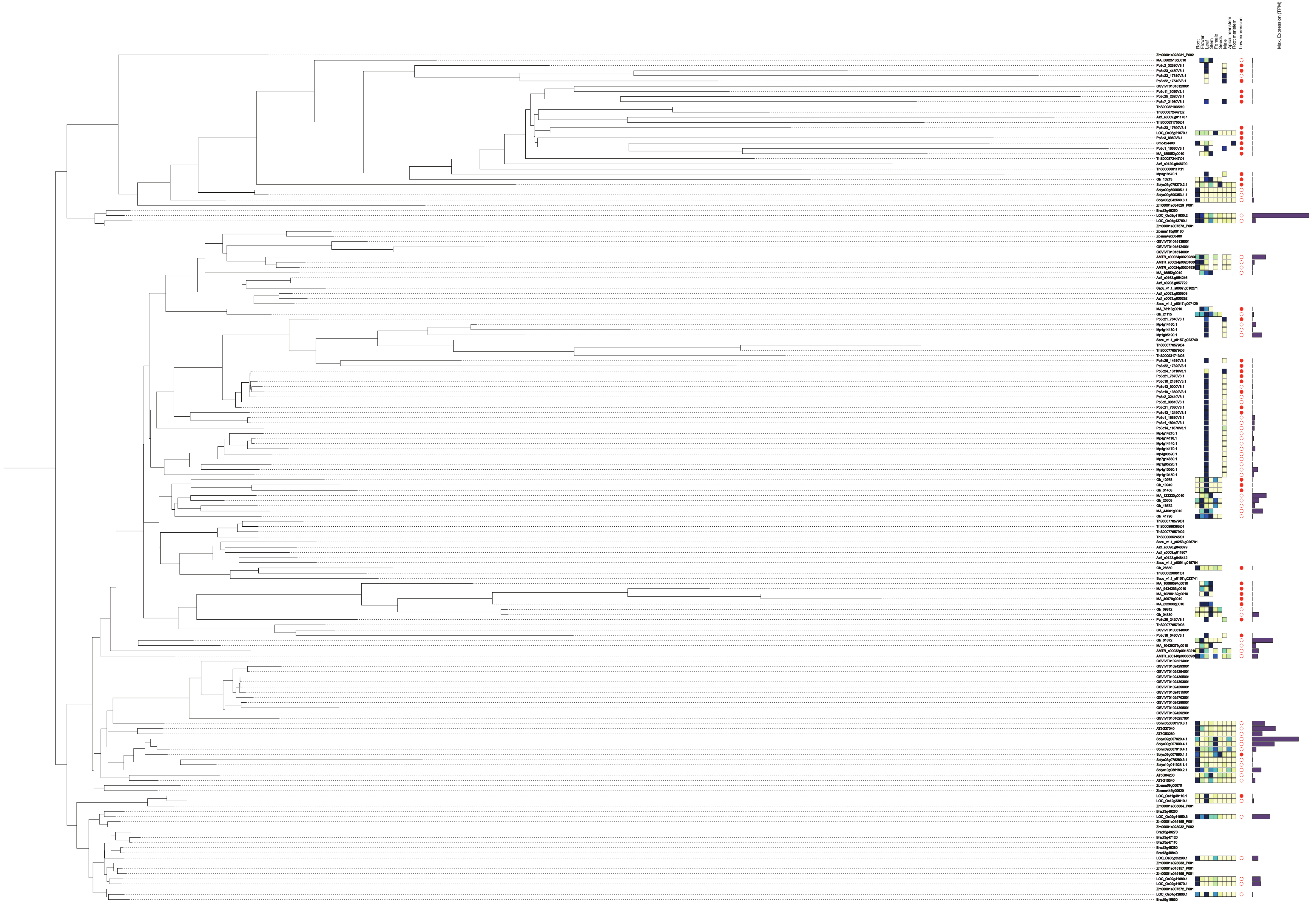


**Figure S16. Phylogenetic Tree of PAL OG_02_0000119 (Land Plants).**
